# Learning the properties of adaptive regions with functional data analysis

**DOI:** 10.1101/834010

**Authors:** Mehreen R. Mughal, Hillary Koch, Jinguo Huang, Francesca Chiaromonte, Michael DeGiorgio

## Abstract

Identifying regions of positive selection in genomic data remains a challenge in population genetics. Most current approaches rely on comparing values of summary statistics calculated in windows. We present an approach termed *SURFDAWave*, which translates measures of genetic diversity calculated in genomic windows to functional data. By transforming our discrete data points to be outputs of continuous functions defined over genomic space, we are able to learn the features of these functions that signify selection. This enables us to confidently identify complex modes of natural selection, including adaptive introgression. We are also able to predict important selection parameters that are responsible for shaping the inferred selection events. By applying our model to human population-genomic data, we recapitulate previously identified regions of selective sweeps, such as *OCA2* in Europeans, and predict that its beneficial mutation reached a frequency of 0.02 before it swept 1,802 generations ago, a time when humans were relatively new to Europe. In addition, we identify *BNC2* in Europeans as a target of adaptive introgression, and predict that it harbors a beneficial mutation that arose in an archaic human population that split from modern humans within the hypothesized modern human-Neanderthal divergence range.

## Introduction

Positive selection is one the most fundamental forces shaping the diversity of life that we can observe today (Bersaglieri et al., 2004; Hill et al., 1991). When positive selection acts on a beneficial mutation, it causes a wave-like pattern in the decrease in diversity of the genome (Smith and Haigh, 1974). As in waves found in the ocean or air, certain patterns might emerge depending on the properties of the cause and the environmental materials (genetic background). Examining these patterns might allow us to learn about the forces causing them. For example, the angle between the crest (top of a wave) and trough (bottom of a wave) might be informative for learning about the strength of selection (and concurrently the time taken for a selective event to occur). Similarly, different modes of positive selection may have on average different patterns. For example, if the crest of the wave extends above the rest position (neutrality), then this may be the signal of adaptive introgression as shown in Setter et al. (2019).

Capturing diversity patterns as they vary spatially has been the goal of a number of recent methods (Schrider and Kern, 2016; Kern and Schrider, 2018; Flagel et al., 2019). Both Schrider and Kern (2016) and Kern and Schrider (2018) attempt to recognize sweeps by learning how diversity (measured by summary statistics) changes across a number of windows encompassing the sweep. However, these methods do not explicitly model the overall patterns formed by selection events. Other methods forgo explicitly measuring diversity and transform SNP data directly to images to learn population-genetic parameters such as recombination rates (Chan et al., 2018; Flagel et al., 2019) and to identify selected regions (Flagel et al., 2019). The complementary approach of Mughal and DeGiorgio (2019) explicitly models the spatial autocorrelation of summary statistics to capture the underlying wave patterns produced by selective sweeps.

Fortunately, there exist techniques not widely applied in genomics that allow observations on continuous data (Cremona et al., 2019). Functional data analysis is a recent sub-field of statistics in which measured values are known to be the output of functions (Ramsay and Silverman, 2005; Wang et al., 2016). Relatedness between data points is inherent in this type of data analysis, which operates on values across a continuum. Transforming our measures of genetic diversity across a genomic region into functional data ensures that the spatial pattern is used to draw conclusions. Although we will be applying this method to assess how genetic diversity varies across the space of a genomic region, there are numerous and diverse potential applications of this approach in evolutionary genomics. With the deluge of ancient genome datasets emerging, it may be possible to examine how the spatial distribution of genetic diversity changes across time at different positively-selected genomic regions to learn their adaptive parameters, such as selection strength, sweep softness, and timing of selection.

We present a method termed *SURFDAWave* (Sweep inference Using Regularized FDA with WAVElets) in which we first model genetic diversity as functions, and then learn the importance of different aspects of genetic diversity across the examined genomic space in predicting selection parameters. We show that *SURFDAWave* accurately predicts parameters such as selection strength, initial frequency of mutation before becoming beneficial, and time of selection. We also demonstrate that *SURFDAWave* can be used to classify selective sweeps, while remaining robust to confounding factors. Finally, we apply *SURFDAWave* to empirical data to predict the selection parameters on regions classified as sweeps.

## Results

*SURFDAWave* is a wavelet-based regression method used to classify selective sweeps and predict adaptive parameters (Figure 1). Here we briefly present its performance in terms of both classification of selective sweeps and in estimating parameters responsible for shaping sweeps. We compare classification performance of *SURFDAWave* to *Trendsetter*, as *Trendsetter* also models the spatial autocorrelation of summary statistics, and has already been extensively compared to other leading methods (Mughal and DeGiorgio, 2019).

**Figure 1:**
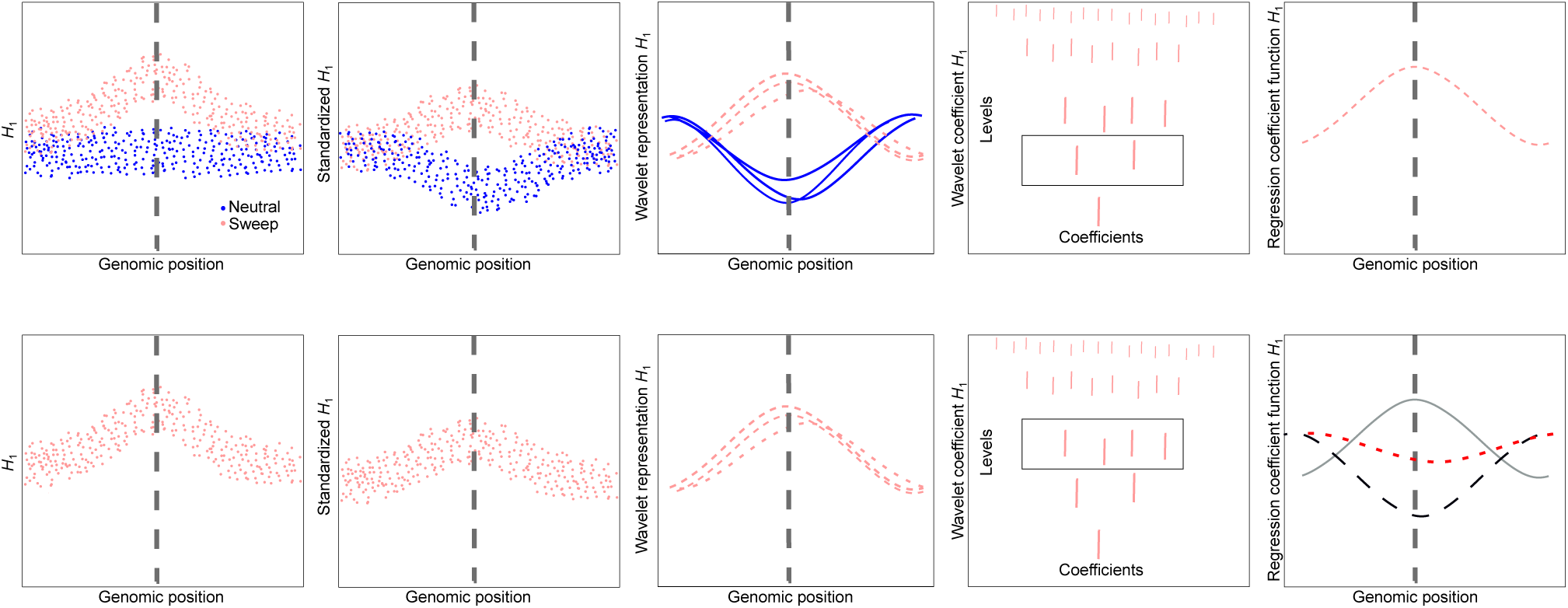
Cartoon illustrating *SURFDAWave* function. For each statistics, *SURFDAWave* standardizes values before transforming values into their wavelet representations (Middle boxes in both rows). The wavelet representation are analyzed at all possible levels from the most detail or highest level to least detail or lowest level (Right of middle boxes in both rows). The top row shows how a binary classifier chooses wavelet coefficients to differentiate between sweeps and neutrality. The bottom row shows how there is a separate model for each selection parameter we predict. In this case the three different colored lines in the right box are the regression coefficient functions for three different selection parameters.

### Classification of selective sweeps

We trained the *SURFDAWave* classifier to differentiate between sweeps and neutrality as described in *Materials and Methods*. For all sweep simulations, we drew selection start time, initial frequency of beneficial mutation, and selection strength of the beneficial allele from a distribution, such that all sweep scenarios comprise a range of hard and soft sweep settings. We initially compared *SURFDAWave* to *Trendsetter*, using the same set of summary statistics calculated from the same simulated datasets. Note that the set of summary statistics and number of windows used here is slightly different from what was originally employed by *Trendsetter* in Mughal and DeGiorgio (2019). However, we chose to focus on how the modeling of the summary statistics, rather than the number or choice of summary statistics, would affect differences in classification rates between *SURFDAwave* and *Trendsetter*. Though it is possible to apply *SURFDAWave* with many combinations of summary statistics in any number *p* = 2^*J*^ windows (where *J* is a non-negative integer), we use an implementation that employs the summary statistics 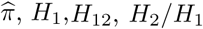, and frequencies of first to fifth most common haplotypes, all calculated in *p* = 128 genomic windows (see *Materials and Methods*).

We first train a classifier using summary statistics calculated on simulations that reflect the CEU European human demographic history (Terhorst et al., 2017). Figure 2 shows that *SURFDAWave* has similar accuracy to *Trendsetter* regardless of the regularization penalty used. This is reflected in the patterns we observe for importance of summary statistics through examining the regression coefficients (*β*s) for each model. Figures S1 and S2 respectively show how *SURFDAWave* and *Trendsetter* both identify *H*_1_ and the frequency of the most common haplotype as uninformative. We also found that the type of wavelets (*i.e.*, Daubechies’ least-asymmetric vs. Haar) used as basis functions does not dramatically influence the overall classification rates (Figure 2). However, visualizing the coefficient functions for each summary statistic shows that the overall shape is much smoother when using Daubechies’ least-asymmetric (Figure S1) compared to Haar (Figure S3) wavelets. We find smoothness of the coefficient functions to be desirable, and for this reason, most of our results are shown using Daubechies’ least-asymmetric wavelets. We also notice that although classification rates are similar regardless of whether we use ridge penalization (*γ* = 0), lasso penalization (*γ* = 1), or choosing the optimal elastic net parameter *γ* through cross validation (Figure 2), the resulting regression coefficient functions are vastly different, especially when we use *γ* = 0 (Figures S1, S4, and S5).

**Figure 2:**
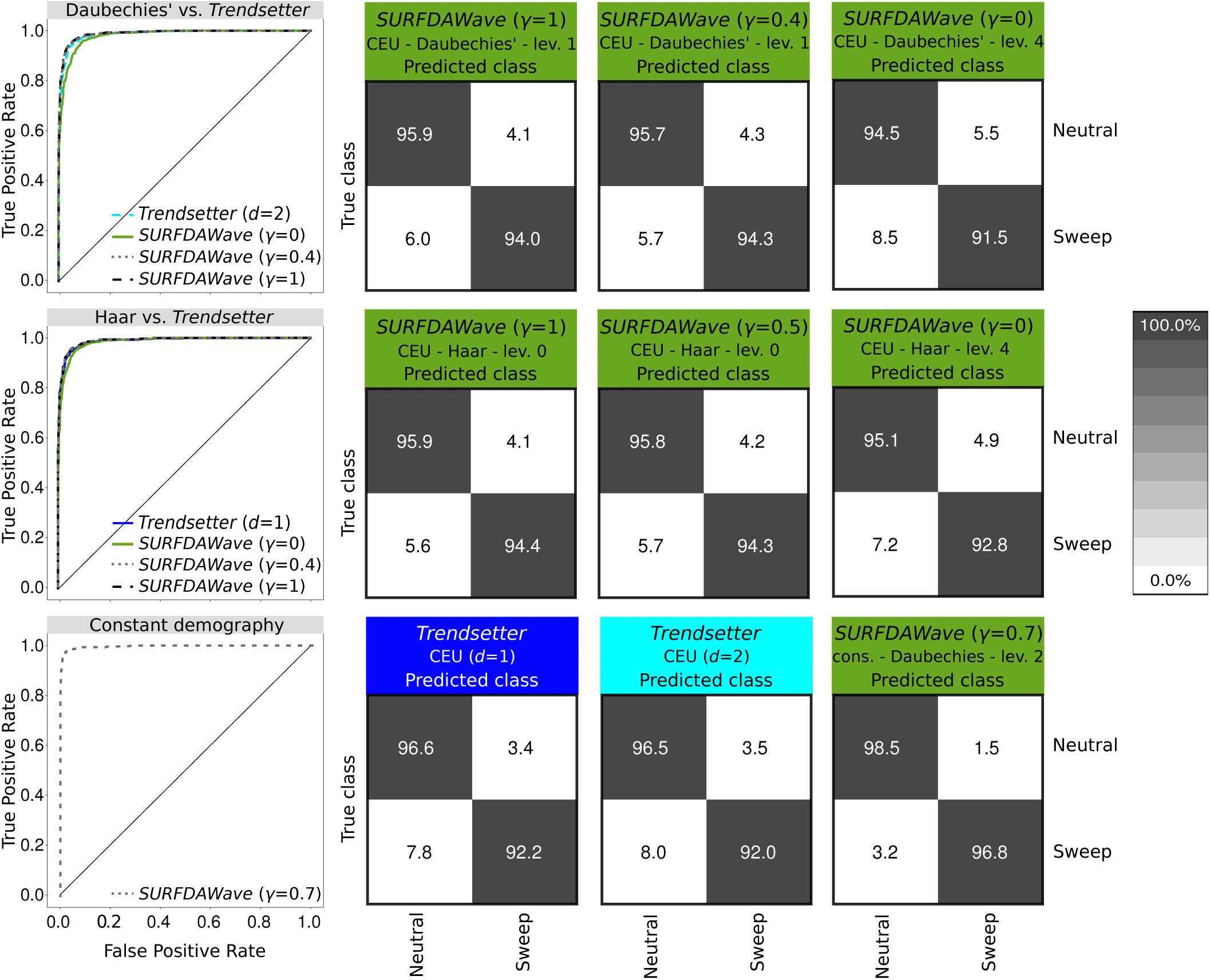
*SURFDAWave* classifier performance on simulated data. (Left column) Power to differentiate between sweep and neutrality by comparing the probability of a sweep under sweep simulations with the same probability in simulations of neutrality when using varying *γ* penalties, wavelet types, and demographic histories. (Top row confusion matrices) Confusion matrices comparing classification rates of *SURFDAWave* using Daubechies’ least-Asymmetric wavelets to estimate spatial distributions of summary statistics when using either *γ* = 1, *γ* = 0, or *γ* chosen through cross validation. (Middle row confusion matrices) Confusion matrices comparing classification rates of *SURFDAWave* using Haar wavelets to estimate spatial distributions of summary statistics when using either *γ* = 1, *γ* = 0, or *γ* chosen through cross validation. (Bottom row confusion matrices) Confusion matrices comparing classification rates of *Trendsetter* with constant (*d* = 1) trend penalty, *Trendsetter* with linear (*d* = 2) trend penalty trained and tested with simulations based on CEU demographic history and *SURFDAWave* with varying elastic net penalties (*γ*) trained and tested with constant demographic history.

We also train a classifier to differentiate between selective sweeps and neutrality using simulations of the YRI sub-Saharan African human demographic history (Terhorst et al., 2017) over a range of *γ* values. Overall, we notice an increase in the percentage of simulations classified correctly when we compare to classifiers trained under the CEU demographic history (Figure 3). However, the patterns formed by the spatial distributions of the coefficients for each summary statistic are similar in both cases (Figures S6-S8). The noisy functions resulting from the use of *γ* = 0 tend to obscure any pattern in the spatial distribution of the underlying regression functions and as a result make the function more difficult to interpret. For this reason we proceed with either *γ* = 1 or *γ* chosen through cross-validation.

**Figure 3:**
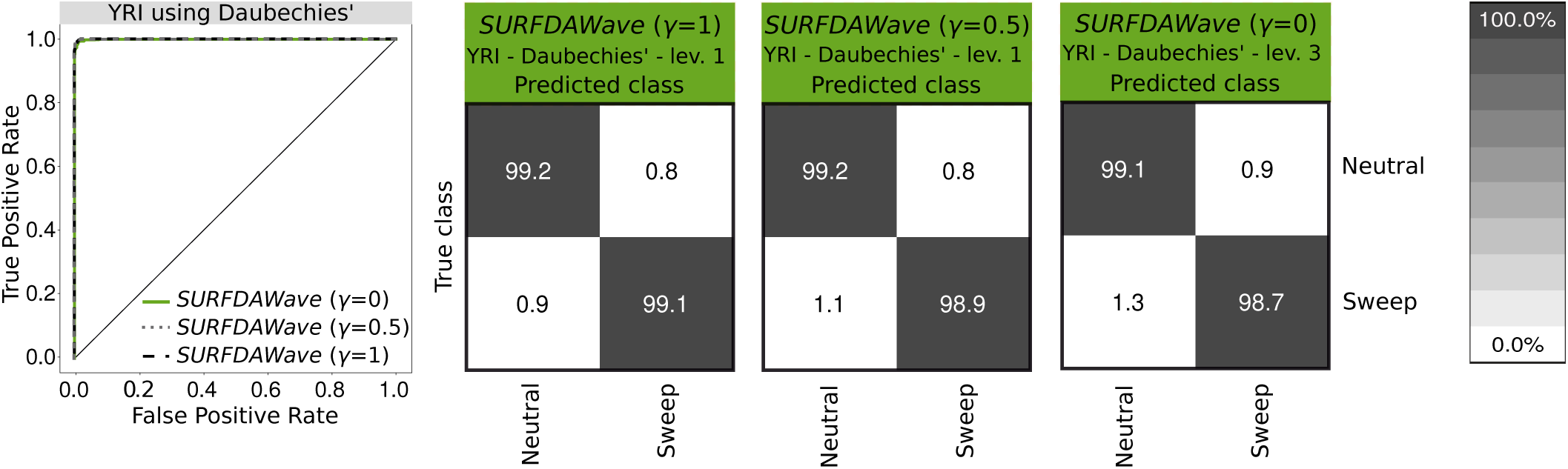
*SURFDAWave* classifier performance on simulated data trained and tested with simulations conducted with YRI demographic history parameters using Daubechies’ least-Asymmetric wavelets to estimate spatial distributions of summary statistic. (Left) Power to differentiate between sweep and neutrality by comparing the probability of a sweep under sweep simulations with the same probability in simulations of neutrality when using varying *γ* penalties. (Right confusion matrices) Confusion matrices showing classification rates using *γ* = 1, *γ* = 0, or *γ* chosen through cross validation.

Through cross-validation we also chose the level at which the discrete wavelet transform (DWT) has best performance for classification. Using wavelets as our basis functions has the advantage of allowing our regression coefficients to be represented at different resolutions, denoted by different levels *j*_0_ (see *Materials and Methods*). Choosing these levels through cross-validation allows our method to determine the smoothness of the regression coefficient function because choosing a coarser resolution (lower level) results in a smooth function, whereas choosing a finer resolution (higher level) will result in a more rugged function. As detailed in *Materials and Methods*, the total number of levels at which DWT can be applied equals log_2_(*p*) − 1, which when *p* = 128 (as is used here) means we have six different levels *j*_0_ ∈ {0, 1, 2, 3, 4, 5}. To illustrate the differences among levels, we show a model using DWT with the coarsest level (*j*_0_ = 0) compared to a model using DWT with the finest (*j*_0_ = 5), with both models employing Daubechies’ least asymmetric wavelets with a lasso (*γ* = 1) penalty (Figure S9). It is clear that the summary statistic *H*_12_ is informative for both of these models, however the noisy wavelet reconstructions seen for *j*_0_ = 5 reveals an emphasis on local features that is absent when we enforce *j*_0_ = 0.

To compare the effect of bottlenecks and expansions on classification rates to those under a constant-size demographic history, we trained and tested a classifier using simulations of a constant-size demographic model to differentiate between neutrality and sweeps. As expected, we find both neutral and sweep simulations are classified correctly more often than when classifying simulations of more complicated non-equilibrium demographic histories (Figure 2), such as those of the CEU and YRI populations.

Adaptive introgression is a complex form of natural selection that produces genetic diversity footprints distinct from typical selective sweeps (Setter et al., 2019). In both selective sweeps and adaptive introgression, diversity generally decreases surrounding the beneficial mutation. In adaptive introgression, however, diversity increases before the signal decays to the level of neutrality. This slight increase in diversity compared to the neutral background is most clearly seen when the two populations (donor and recipient) are highly diverged (Figure S10). We test how well *SURFDAWave* can differentiate among adaptive introgression, sweeps, and neutrality, using the same summary statistics discussed in the *Results* section (see *Material and Methods* for simulation details). Similar to previous sweep simulations, for adaptive introgression simulations we drew selection start time, initial frequency of beneficial mutation, selection strength of the beneficial allele, and the donor and recipient divergence time from a distribution, such that all adaptive introgression scenarios comprise a range of hard and soft sweep settings. As shown in Figure 4, we see that *SURFDAWave* is only able to correctly classify sweep simulations in 52.5% of cases, misidentifying them as adaptive introgression 43.2% of the time under the CEU-based simulations with similar results for YRI. As we saw in Figure S10, this is because when divergence times for donor and recipient populations are recent, then the signature of adaptive introgression looks more like a selective sweep.

**Figure 4:**
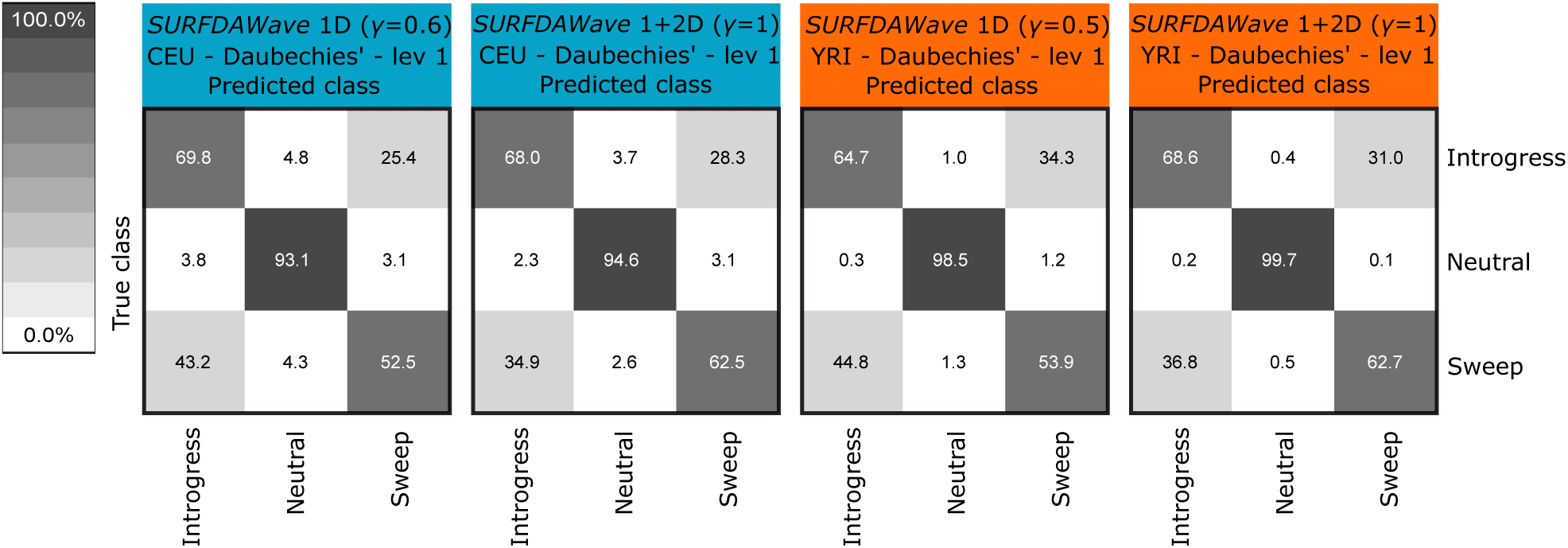
*SURFDAWave* classification rates when differentiating among adaptive introgression, sweeps, and neutrality. The two left columns are showing classification rates when trained and tested with simulations conducted under CEU European demographic history specifications, and the two right columns show the same for with simulations for the YRI Yoruban population. The first and third confusions matrix show results using summary statistics 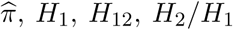, and frequencies of first to fifth most common haplotypes, while the column on the second and fourth confusion martix show results including the preceding statistics as well as the mean, varaiance, skewness, and kurtosis of *r*^2^. The *γ* for the classifiers trained with only the one dimensional statistics is chosen through cross validation, but is specified *γ* = 1 for the results in the column on the right. The level is chosen through cross validation for all models.

To investigate whether the inclusion of other summary statistics, which may better assess genomic variation, boosts our performance we include an additional set of summary statistics, specifically adding the mean, variance, skewness, and kurtosis of the squared correlation coefficient *r*^2^ (Hill and Robertson, 1968) calculated between all possible SNPs sampled from each pair of windows (see *Materials and Methods*). Because visualizing these statistics in square matrices is informative, we refer to them as two-dimensional statistics, and refer to 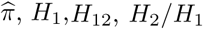, and frequencies of first to fifth most common haplotypes as one-dimensional statistics. We see that the inclusion of two-dimensional statistics increases the correct classification of selective sweeps substantially for both populations to 62.5% in CEU and 62.7% in YRI (Figure 4). The percent of adaptive introgression simulations classified correctly also increased for YRI going from 64.7% to 68.6%. We can see how the inclusion of the two dimensional statistics affects the model by directly comparing the reconstructed wavelets across the spatial distributions of the nine summary statistics included in both models. By examining the coefficients of the two-dimensional statistics for the model using both types of statistics, we can see that the skewness and kurtosis of *r*^2^ are informative in separating neutrality from the other classes (Figures 5 and S11). Interestingly, the statistic *H*_1_ is important in separating neutrality from both types of selection in the model including two-dimensional statistics for the CEU demographic history, but clearly does not serve this purpose in the model trained with only one-dimensional statistics (Figures S12 and S13).

**Figure 5:**
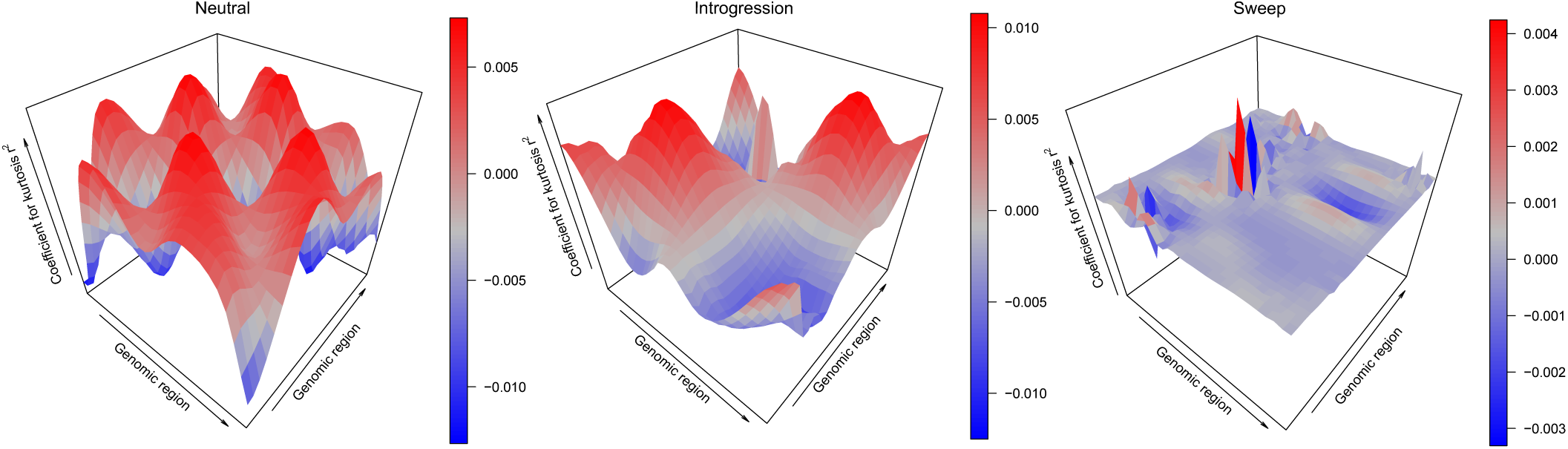
Three dimensional representations of reconstructed wavelets from regression coefficients (*β*s) when differentiating among adaptive introgression, sweeps, and neutrality scenarios for summary statistics kurtosis of pairwise *r*^2^ for *SURFDAWave* when *γ* = 1, when trained with statistics 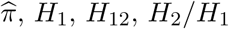, frequencies of first to fifth most common haplotypes, and mean, variance, skewness, and kurtosis of pairwise *r*^2^. *SURFDAWave* was trained on simulations of scenarios simulated under demographic specifications for European CEU demographic history. Note that the wavelet reconstructions for all summary statistics are plotted on the same scale, thereby making the distributions of some summaries difficult to decipher as their magnitudes are relatively small. *SURFDAWave* results shown are using Daubechies’ least-asymmetric wavelets wavelets to estimate spatial distributions of summary statistics. Level 1 chosen through cross-validation.

### Classification with confounding factors

Testing *SURFDAWave* on simulations of biological events that might confound classification is necessary to ensure that it can be applied under diverse empirical scenarios. For this reason we test classification performance of *SURFDAWave* under simulations with extensive missing data. To simulate missing data as we might find it in genome sequences due to technical issues, such as alignability and mappability (Mallick et al., 2009), we remove large randomly spaced chunks of the simulated data (see *Materials and Methods*). We show that missing data does not substantially affect the performance of *SURFDAWave* when classifying neutral simulations with missing data (Figure 6). We do, however, observe a slight decrease in performance in the classification of selective sweeps when sweep simulations are missing data, with an increase in the percentage of sweep simulations missing data being classified as neutral. This robustness to missing data can be attributed to the types of summary statistics applied and the manner in which they are calculated using SNP-delimited windows (Mughal and DeGiorgio, 2019). Another common confounding factor is background selection, in which deleterious mutations cause a loss of diversity which might be confused for selection signatures (Charlesworth, 2013). For this reason we test *SURFDAWave*’s performance on background selection, which we simulate based on the distribution of effect sizes and spatial distribution of coding elements in the human genome, as in Schrider and Kern (2017) and Mughal and DeGiorgio (2019) (see *Materials and Methods* for details). We find that 93.4% and 94.2% of background selection scenarios under the CEU and YRI demographic histories, respectively, are classified as neutral (Figure 6).

**Figure 6:**
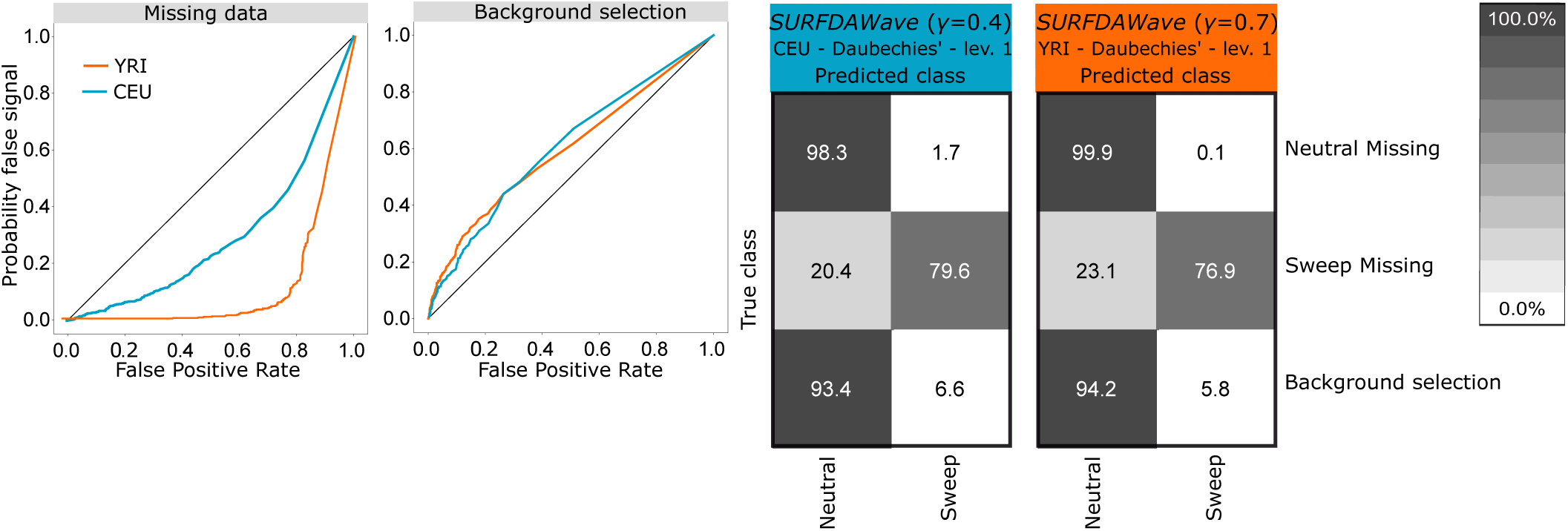
*SURFDAWave*’s robustness to missing data and background selection. (Left) Probability of mis-classifying neutrally-evolving genomic regions missing data as a sweep. Comparing probability of sweep in simulations missing data (probability of false signal) with the probability of any sweep in neutral simulations. (left-middle) Probability of mis-classifying background selection simulations as sweep. Comparing the probability of a sweep in simulations of background selection with probability of sweep in neutral simulations. (Right) Confusion matrices showing classification rates of *SURFDAWave* when classifying simulations of each class with missing data, and when classifying background selection simulations. *SURFDAWave* is trained to differentiate sweeps and neutrality using Daubechies’ least-Asymmetric wavelets. Optimal *γ* and and level were chosen through cross validation.

Along with issues of background selection and missing data, there are also known difficulties with establishing accurate demographic histories of present populations. Similar to the results shown in (Mughal and DeGiorgio, 2019), we again show that *SURFDAWave* loses performance when demographic specifications are less accurate. With a classifier trained to differentiate between sweeps and neutrality in CEU European populations, we mis-classify 37.9% of sweep simulations conducted under YRI sub-Saharan African demographic history as neutral. However, the percentage of neutral YRI simulations classified as neutral increases to 99.1% when tested with a CEU trained demographic history. In the opposite case, with the classifier trained to differentiate sweeps from neutrality with simulations of YRI, when we test sweep simulations conducted under the CEU demographic history we classify 99.2% correctly, however we mis-classify simulations of neutrality as sweeps 26.0% of the time (Figure S14). These mis-classifications should be rescued if *SURFDAWave* was trained across a diverse set of demographic histories (*e.g.*, Mughal and DeGiorgio, 2019).

### Prediction of selection parameters

Classification of selective sweeps provides a limited understanding of the evolutionary processes shaping genomic regions. To gain deeper insight about the underlying adaptive processes, we also tested the ability of *SURFDAWave* to predict the selection parameters involved in shaping sweeps. We trained a multi-response linear regression model to jointly learn the log-scaled initial frequency of the adaptive allele prior to it becoming beneficial, the log-scaled selection coefficient, and the time at which the mutation becomes beneficial (see *Materials and Methods*) using demographic specifications for the CEU and YRI populations. We include the same set of *m* = 9 summary statistics as used to train the sweep classifier in the preceding section, each computed across *p* = 128 windows. Prediction of initial frequency, selection coefficient, and time of selection is accurate (Figure S15) with the root mean squared error (RMSE) equal to 0.49 for the log-scaled selection coefficient, 0.43 for the log-scaled initial frequency, and 20.3 for time at which selection began for unstandardized log-scaled selection coefficient, unstandardized log-scaled initial frequency, and the unscaled and unstandardized time of selection, respectively (Figure 7). The RMSE for the YRI population is similar to that of the CEU (Tables S1 and S2). Visualizing the coefficient functions after regularized regression conveys that most summary statistics are informative in predicting parameters (Figures S16 and S17), with the exception of the frequency of the most common haplotype, which is flat across the entire spatial distribution in both models.

**Figure 7:**
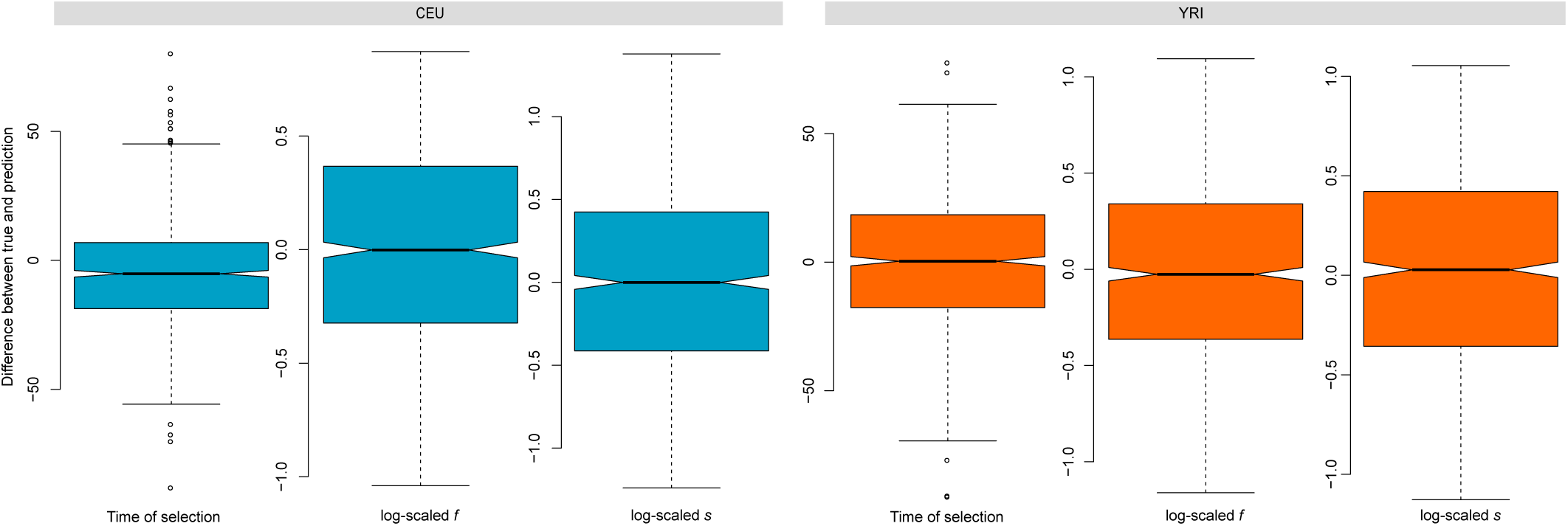
Difference between unstandardized selection parameters with *SURFDAWave* for the CEU and YRI demographic models. (Left box plot) Difference in prediction and truth of log scaled time at which mutation became beneficial. (Middle box plot) Difference in prediction and truth of log scaled frequency reached by mutation prior to it becoming beneficial (*f*). (Right box plot) Difference in prediction and truth of log scaled selection coefficient (*s*).

To test the influence of confounding factors such as missing data on the prediction model, we simulate missing data as in the *Classification with confounding factors* section above. We find that predicting parameters with missing data increases RMSE slightly (Tables S3 and S4). We also test robustness of the *SURFADAWave* prediction model to demographic mis-specification, by considering test simulations performed under CEU demographic specifications with a model trained with simulations performed under YRI demographic specifications, and vice versa (Tables S5 and S6). Again, we find that the RMSE increases compared to training and testing with the same population demographics for both experiments, but the RMSE is less than the error due to missing data. We also simulate selective sweeps including background selection, simulated as described in *Classification with confounding factors*, with the exception of including a beneficial mutation in the center of the simulated chromosome. We find that the RMSE values are very close to the RMSE with no confounding factors (Tables S7 and S8).

In addition to being able to predict the selection parameters responsible for shaping classical selective sweeps, we also probed whether *SURFDAWave* could predict selection parameters important in shaping sweeps due to adaptive introgression. An interesting parameter specific to adaptive introgression is the time at which the donor and recipient populations diverged. Instead of predicting the time at which a mutation became beneficial, as we show above in *Prediction of selection parameter*, we train models to predict the donor-recipient split time, along with the selection strength and initial frequency of the mutation before it became beneficial (Table S9 and Figure S18). The RMSE values for the selection strength and time of selection are similar to the values predicted for regular selective sweeps (Table S1).

### Application to empirical data

Using variant calls in the CEU and YRI populations from the 1000 Genomes dataset (The 1000 Genomes Project Consortium, 2015), *SURFDAwave* recapitulated many of the classical sweep candidates observed by other studies, and moreover classified the vast majority of the CEU and YRI genomes as neutral (Tables S10 and S11), with a greater percentage of the YRI genome being classified as neutral than the CEU genome. This is the result of a combination of factors, including our classifiers reduced ability to distinguish sweeps from neutrality in populations with complex demographic histories, such as the CEU population (see *Classification of selective sweeps*). To make our method more conservative, we applied a probability threshold for selective sweeps. If the probability of a selective sweep is less than or equal to 0.7, then we consider this region to be neutral. Figure S19 shows that the *SURFDAWave* classifier predicts probability distributions close to actual probability distributions, which validates our use of a probability threshold. Among the genes classified as selective sweeps in the CEU population, we found *LCT, OCA2*, and *SLC45A2*, which were previously hypothesized as targets of selection (Voight et al., 2006; Bersaglieri et al., 2004; Wilde et al., 2014; Sulem et al., 2007) (Figures S20-S22). In the YRI population we classify the genes *SYT1, HEMGN, GRIK5*, and *NNT* as under positive selection, recapitulating the work of Harris et al. (2018), Fagny et al. (2014), and Pickrell et al. (2009) (Figures S23-S26).

As we have already trained models to jointly predict the selection strength, the time at which the mutation became beneficial, and the frequency of the adaptive mutation before becoming beneficial, we next use all of the human genome regions classified as sweeps to learn about the underlying parameters shaping variation at these candidates. We first examined *OCA2*, a gene that is involved in eye coloration (H. Brilliant, 2001; Zhu et al., 2004; Eiberg et al., 2008), and predicted that the time at which a mutation on this gene became beneficial was 1,802 generations ago, and that the beneficial mutation had a selection strength of *s* = 0.06 and an initial frequency of *f* = 0.02. This prediction is made on the set of statistics classified as sweep with the highest probability in the region containing the gene *OCA2* with 0.978 probability. Using a generation time of 29 years for humans, implies the mutation became beneficial about 52,258 years ago, a time during which modern humans were relatively new to Europe (Hublin, 2012). *SLC45A2*, another gene involved in pigmentation (Cook et al., 2009), harbors a test window with a sweep probability of 0.694 and the predicted selection strength, initial frequency, and selection time are *s* = 0.04, *f* = 0.02, and 2,000 generations ago, respectively. In the YRI population we predict that a mutation on *HEMGN*, a gene that regulates the development of blood cells (Li et al., 2004), first became beneficial 1,960 generations ago and has a selection coefficient of *s* = 0.03 and frequency at which it became beneficial of *f* = 0.016. We predict that the selective sweep occurring on the region around *SYT1*, mutations on which are associated with neurodevelopmental disorders (Baker et al., 2018), began 2,260 generations ago with a selection coefficient of *s* = 0.04 and an initial frequency of *f* = 0.02.

In the list of 444 genes in YRI classified as sweep with probability greater than or equal to 0.7, we examine the range of predictions for each parameter and the genes predicted to have selection parameters at the fringes of each range. For each gene, we only include the prediction for the feature vector where the predicted probability of classification as sweep is the highest within that gene. We find that the gene with the minimum selection coefficient within this list is *HCG23*, with an inferred coefficient of *s* = 0.018. We inferred that a sweep initiated on this gene 1,259 generations ago when the initial frequency of the beneficial mutation was *f* = 0.017. The highest probability of this gene being classified as a sweep is 0.986. We also predict that the gene with the highest initial frequency also had the most recent sweep initiation time. This gene, *STPG2* (Sperm-tail PG rich repeat containing protein 2), is highly expressed in the testis (Uhlén et al., 2015), and we predict that this had a mutation reach a frequency of *f* = 0.039 about 666 generations ago, at which point it was predicted to become beneficial.

In a similar examination of the CEU population, we find 2,265 genes are classified as sweep with a probability greater than 0.7. The oldest selection time we predict (2,922 generations or 84,738 years, with a generation time of 29 years) occurred on *VPS35*, a gene on which mutations are associated with Parkinson’s Disease (Vilario-Gell et al., 2011). We infer that strong selection (*s* = 0.05) began on this gene when a selected mutation reached a frequency of *f* = 0.016. *HLA-DRB1* plays an important role in the immune system and has been previously predicted to be under balancing selection (Bronson et al., 2012). We find that this gene has the highest inferred selection coefficient of *s* = 0.14 and lowest inferred initial frequency of *f* = 0.004 out of our set of genes for the CEU. This may be indicative of a mutation around this region becoming immediately beneficial after it occurred, which we predict was about 1,718 generations ago.

We compare the distributions of selection parameters for the sets of likely selected genes discussed above. In Figure S27, we can see that while some genes are predicted to have more recent times of selection, most are predicted to have a time of selection greater than 2000 generations ago in both populations. Among the more recent sweeps, we also find a greater range of predicted initial frequencies than those genes that were predicted to have an earlier selection start time. Overall, the distributions for predicted parameters in both populations overlap extensively for all selection parameters (Figure 8). We also observe that *SURFDAWave*’s prediction of the initial frequency (*f*) is not dependent on the probability of a sweep (Figure S28), but as the probability of sweep increases *SURFDAWave* is are more likely to predict stronger selection coefficients (*s*) and slightly more recent selection start times.

**Figure 8:**
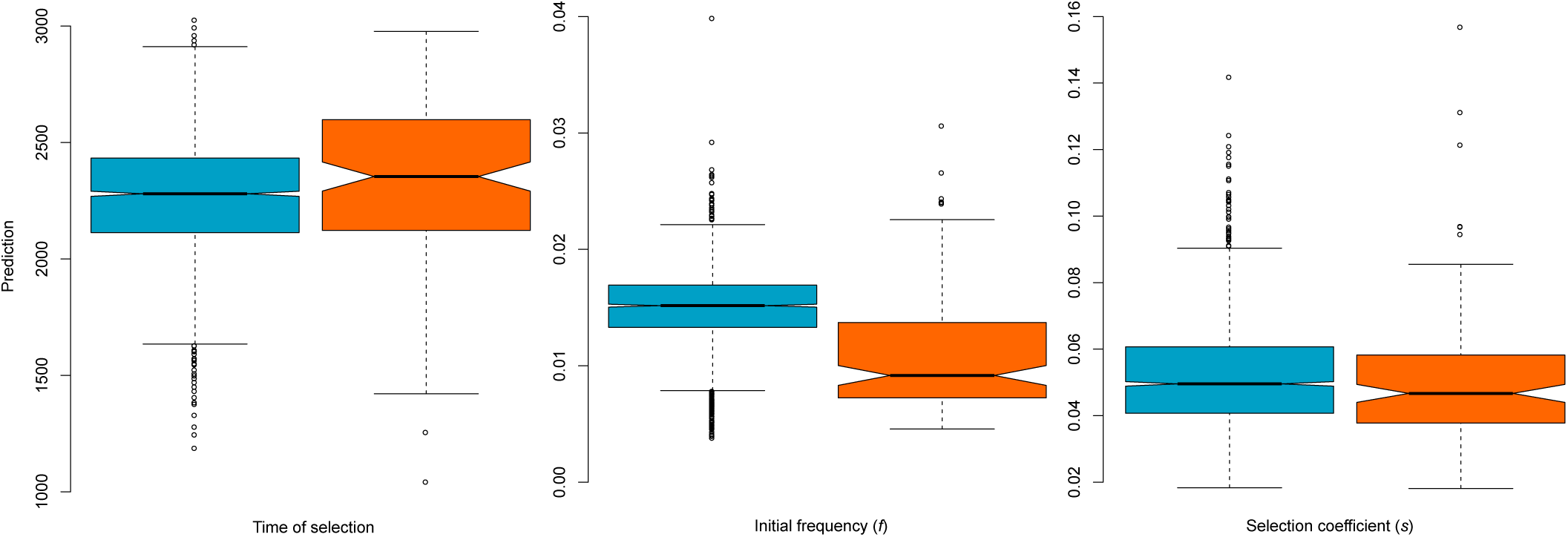
Predicted distribution of selection parameters for all genes in YRI (orange) and CEU (blue) with probability of being classified as sweep greater than 0.7 (Left) Distribution of predicted time of selection. (Middle) Distribution of predicted frequency reached by mutation before becoming beneficial (initial frequency (*f*)). (Right) Distribution of predicted selection coefficient (*s*).

Finally, we apply the classification and prediction models to locate adaptive introgression and learn the adaptive introgression parameters. We find that regardless of the types of statistics used, we classify the majority of the genome as neutral, and classify more of the genome as sweep than as adaptive introgression (Tables S12-S15). Importantly, we find that we are able to recapitulate signals of previously-identified regions of adaptive introgression in the CEU population with *SURFDAWave*, such as *BNC2* (Sankararaman et al., 2014; Racimo et al., 2015) and *APOL4* (Setter et al., 2019) (Figures S30 and S31). *BNC2* is another gene thought to play a role in human skin color determination (Visser et al., 2014), whereas the gene *APOL4* is significantly up-regulated in people diagnosed with schizophrenia (Monajemi et al., 2002). By applying the *SURFDAWave* prediction models to the summary statistic computed at these genes, we estimate that the beneficial mutation in *APOL4* reached an initial frequency of *f* = 0.05 and had a selection strength of *s* = 0.01, with the donor and recipient populations splitting 19,760 generations, or about 573,000 years, ago (using a generation time of 29 years). We also estimate that the selection strength on the *BNC2* gene is stronger and harder than the signature on *APOL4*, with *s* = 0.04 and *f* = 0.01. Moreover, the predicted donor and recipient split time of 20,180 generations (585,220 years) ago from variation at *BNC2* is similar to the estimate from *APOL4*.

## Discussion

In this article, we demonstrated that *SURFDAWave* is able to locate selective sweeps, and also predict selection parameters responsible for shaping those sweeps. Moreover, we showed that *SURFDAWave* is capable of differentiating between sweeps and neutrality, and is also able to accurately predict the time at which the selected mutation became beneficial, the frequency a mutation reached before becoming beneficial, and the selection coefficient. In addition, using image-based feature vectors increased our ability to differentiate among neutrality, adaptive introgression, and sweeps. We were able to recapitulate earlier findings by predicting genes as adaptive that were previously hypothesized to be under positive selection.

Our results show that capturing the spatial distribution of selective sweeps is informative for identifying adaptive regions and learning about the evolutionary parameters that shape them. Moreover, our *SURFDAWave* approach is not restricted to application on adaptive introgression and selective sweep scenarios, and can be implemented for probing genomic variation of other evolutionary processes that leave a spatial or temporal signature in genomic data. Such examples include the identification of genomic targets of balancing selection (*e.g.*, DeGiorgio et al., 2014; Siewert and Voight, 2017; Bitarello et al., 2018; Cheng and DeGiorgio, 2018; Siewert and Voight, 2018; Cheng and DeGiorgio, 2019), complex forms of adaptation such as staggered selective sweeps (Assaf et al., 2015) that have yet to be interrogated for in genomic data, and non-adaptive processes such as recombination rate estimation (*e.g.*, Chan et al., 2018; Flagel et al., 2019).

There are a number of potential applications of our methodological framework. For one, it is possible to naturally extend *SURFDAWave* to incorporate genomic data from ancient samples, and several recent studies have employed ancient DNA to directly examine temporal allele frequency fluctuations to identify positively-selected loci (*e.g.*, Bollback et al., 2008; Ludwig et al., 2009; Mathieson et al., 2015; Fehren-Schmitz and Georges, 2016; Schraiber et al., 2016; Loog et al., 2017). *SURFDAWave*’s framework would allow examination of changes in the spatial distribution of genetic diversity over time by incorporating information from ancient genomes of a single population at various time points throughout history, and summarizing patterns of variation using two-dimensional wavelet bases. However, a specific limitation of the implementation of *SURFDAWave* as we describe it here is that, for each dimension, its application is restricted to using feature vectors of length *p* in which log_2_(*p*) is a non-negative integer. We acknowledge that this constraint may make it difficult for *SURFDAWave* to be widely applied, especially when incorporating information from ancient DNA. Though we choose to use wavelets in our implementation, other basis functions that do not have such limitations on numbers of features, such as B-spline and polynomial basis functions (Ramsay and Silverman, 2005), can be used instead. However, unlike wavelets, these bases do not form orthonormal basis functions, and using them results in more complicated functional regression models.

Along with *SURFDAWave*’s flexibility in terms of classification problems, we also demonstrated that this framework can be adapted to predict different selection parameters. Our results suggest that *SURFDAWave*’s ability to predict the time of selection is better than its ability to predict the split time of the donor and recipient populations. This may be because the spatial distribution of genetic diversity is more affected by the time at which the examined mutation became beneficial, than the split between donor and recipient populations. It is possible, however, that introgression patterns in species in which donor and recipient populations have greater divergence times would leave a more prominent footprint, and allow better predictions of their divergence time to be made (Figure S10).

In addition, we showed how incorporating different types of features, specifically two-dimensional statistics improves the classification ability of *SURFDAWave* (Figure 4). Several recent innovative approaches have explored the use of image-based or two-dimensional features to predict population-genetic processes. For example, Flagel et al. (2019) use the derived or ancestral states from population simulation data directly rather than extracting information from these simulations through the use of summary statistics, and convert this information to images. This raw information can also be converted into wavelet data prior using it as a feature in classification or prediction models. Along with the flexibility that *SURFDAWave* provides in terms of feature input (*e.g.*, one- or two-dimensional statistics), other potential enhancements may increase its prediction and classification accuracy. In our application we assume a linear model. However, it is possible that a linear model is not an accurate representation, and instead employing a more flexible model would enhance our predictions if the actual relationship is non-linear. Therefore, using non-linear model such as a neural network with at least one hidden layer (Cybenko, 1989; Gao et al., 2019) in place of simple linear and logistic regression models may be able to improve the performance of *SURFDAWave*. An implementation of *SURFDAWave* along with results for genome wide scans for sweeps discussed in this article can be downloaded from http://degiorgiogroup.fau.edu/surfdawave.html.

## Materials and Methods

### Wavelet estimation of summary statistic spatial distribution

Consider a sample of *n* training examples, in which *m* summary statistics are computed at *p* positions along a genomic region. Let **x**_*i,s*_ = [*x*_*i,s*,1_, *x*_*i,s*,2_, …, *x*_*i,s,p*_]^*T*^ denote the vector of values for summary statistic *s, s* = 1, 2, …, *m*, for training example *i, i* = 1, 2, …, *n* caculated at each of the *p* positions in a genomic region, where *x*_*i,s,j*_ is the value of the summary statistic at position *t*_*j*_, *j* = 1, 2, …, *p*. For convenience, define the vector 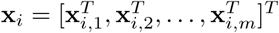 containing the values of each of the *m* summary statistics calculated at the *p* positions.

Each vector of summary of summary statistics **x**_*i,s*_ is the result of some unknown function *f*_*i,s*_(*t*) defined on genomic position *t*. The relationship between the function and the summary statistic data points can be represented as

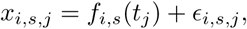

where *f*_*i,s*_(*t*_*j*_) is the function *f*_*i,s*_(*t*) evaluated at position *t*_*j*_ of summary statistics *s* in observation *i*, and where *ϵ*_*i,s,j*_ is an error term associated with observation *i* that is normally distributed with mean zero and standard deviation one. As in Ramsay and Silverman (2005), we can approximate this function *f*_*i,s*_(*t*) as a linear combination of a set of *B* orthonormal basis functions {*φ*_1_(*t*), *φ*_2_(*t*), …, *φ*_*B*_(*t*)} as

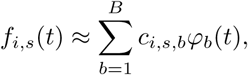

where *c*_*i,s,b*_, *b* = 1, 2, …, *B*, denotes the coefficient of the *b*th basis function *φ*_*b*_(*t*) associated with summary statistic *s* of observation *i*. Note by definition of the *B* basis functions being orthonormal, we have

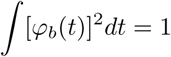

for *b* = 1, 2, …, *B* and

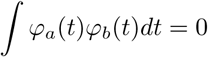

for *a* ≠ *b* (Daubechies, 1988). Orthonormal basis functions commonly used in functional data analysis include wavelets (Nason, 2008) and the Fourier functions (Ramsay and Silverman, 2005). The number *B* of basis functions is a parameter, and is chosen through cross validation. Basis functions are independent functions that can be combined to approximate more complex functions. Here we approximate the function *f*_*i,s*_(*t*) using wavelets at a detail level of *j*_0_ (Zhao et al., 2012) as

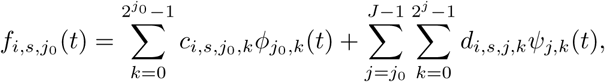

where *J* = log_2_(*p*) is the number of detail levels, *ϕ*_*j,k*_(*t*) and *ψ*_*j,k*_(*t*) are the the father and mother wavelet basis functions at scale *j* and location *k*, respectively, and *c*_*i,s,j,k*_ and *d*_*i,s,j,k*_ are the coefficients for the father and mother wavelets at scale *j* and location *k* for summary statistic *s* in observation *i*. Note that the father and mother wavelet bases form an orthonormal basis (Daubechies, 1988).

### Penalized functional multinomial regression to classify genomic regions

After approximating functions 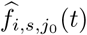 of each summary statistic *s* in observation *i* at detail level *j*_0_, we then use these functions (*i.e.*, their associated coefficients) as the independent variables to model multinomial regression. Denote the vector of length *p* = 2^*J*^ containing estimated father and mother basis coefficients for summary statistics *s* in observation *i* at detail level *j*_0_ as

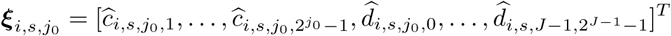

Furthermore, define the concatenated vector of length *m* × *p* of such coefficients across all *m* summary statistics for observation *i* by

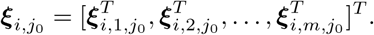

As in Mousavi and Sørensen (2017), we model

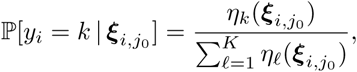

where

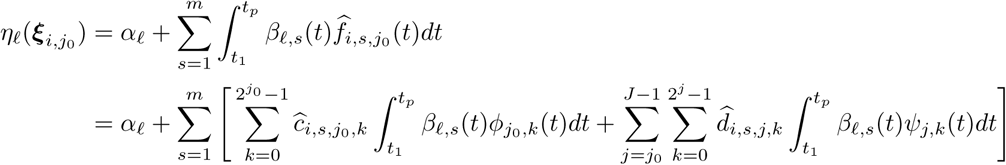

for *ℓ* = 1, 2, …, *K*. This is similar to other multinomial regression models, with the the caveat that we replaced the summation with an integration across the interval [*t*_1_, *t*_*p*_] for position *t*. Here *i* is the index for the observation number, *y*_*i*_ is the categorical response variable with values *y*_*i*_ = *ℓ* for class *ℓ*, for *ℓ* = 1, 2, …, *K, α*_*ℓ*_ is the intercept parameter for class *ℓ*, and *β*_*ℓ,s*_(*t*) is the function for summary statistic *s* of class *ℓ*.

To learn the functions *β*_*ℓ,s*_(*t*), we can note that we may also approximate them with the same set of basis functions as we did for approximating *f*_*i,s*_(*t*). That is, we can approximate the function *β*_*ℓ,s*_(*t*) using wavelets at a detail level of *j*_0_ as

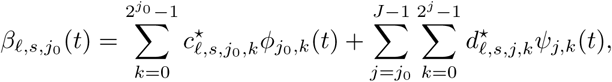

where 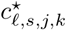 and 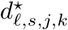 are the coefficients for the father and mother wavelets at scale *j* and location *k* for summary statistic *s* in class *ℓ*. Denote the vector of length *p* = 2^*J*^ containing father and mother basis coefficients for summary statistic *s* for class *ℓ* at detail level *j*_0_ as

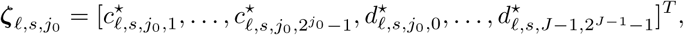

and further define the concatenated vector of length *m* × *p* of such coefficients across all *m* summary statistics for class *ℓ* by

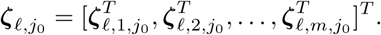

Plugging in this approximation, and using the orthnormality of the set of basis functions, we obtain

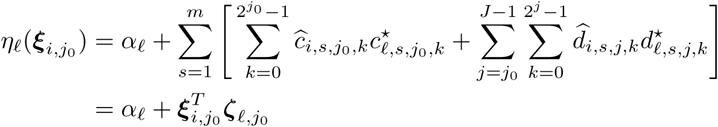

which yields

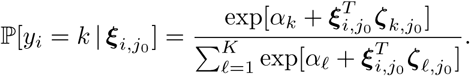

Let ***α*** = [*α*_1_, *α*_2_, …, *α*_*K*_]^*T*^ denote the vector of intercept terms for each of the *K* classes and define the matrix 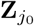 containing *m* × *p* rows and *K* columns by

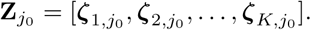

The log likelihood of observing the set of model parameters 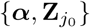 given the collection of data points 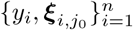 is

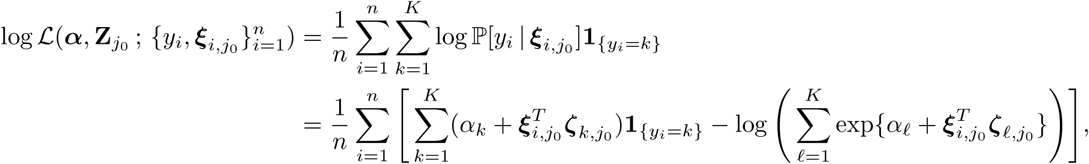

where 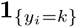 is an indicator random variable that takes the values one if *y*_*i*_ = *k* and zero otherwise.

From this likelihood function, we wish to estimate the intercept terms ***α*** and the coefficients 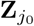. Define 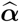 as an estimate of ***α*** and 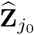 an estimate of 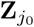. Moreover, as our model is over-parameterized, we need to maximize a penalized log likelihood function. Denoting ‖ · ‖_1_ and ‖ · ‖_2_ as the *ℓ*_1_ and *ℓ*_2_ norms, respectively, define

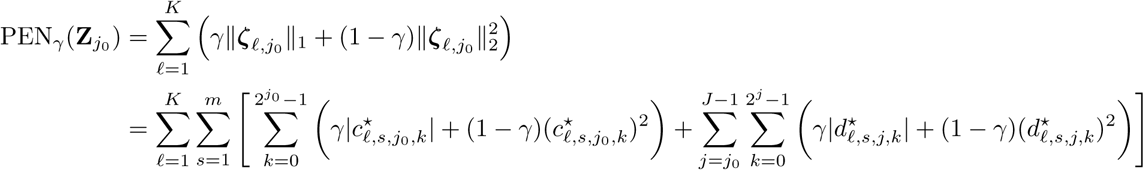

to be the elastic-net penalty (Zou and Hastie, 2005) controlled by parameter *γ* ∈ [0, 1] on the coefficients for the basis functions of the regression coefficient functions, and let *λ* denote a tuning parameter associated with this penalty. A value of *γ* = 0 leads to the standard ridge regression penalty, and *γ* = 1 leads to the lasso penalty. We can therefore estimate the coefficient functions as

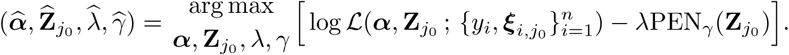

To perform this estimation, we first learn the underlying functions *f*_*i,s*_(*t*) based on orthonormal wavelet basis functions at detail level *j*_0_, yielding the estimated set of coefficients 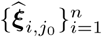 and hence estimated functions

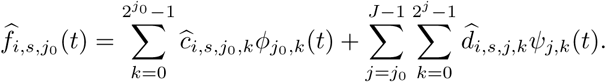

These basis function coefficients are then employed as input covariates to the penalized regression model, for which ten-fold cross validation is used to estimate the tuning parameter *λ*, the tuning parameter *γ* controlling the elastic-net penalty, and associated parameters ***α*** and 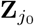. This process is repeated for different detail levels *j*_0_ = 0, 1, …, *J* − 1 to estimate the *j*_0_ that minimizes the ten-fold cross validation error, and the best fitting values of regression model parameters 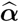 and 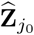 are estimated. These estimates lead to a classifier for future input data, as well as learned functions

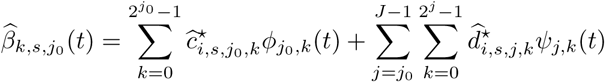

for summary statistic *s, s* = 1, 2, …, *m*, in class *k, k* = 1, 2, …, *K*. After parameter inference, the most likely class 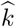 is estimated as

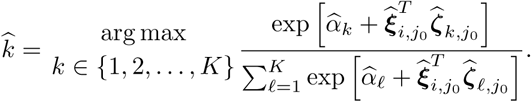

### Penalized functional linear regression to infer evolutionary parameters

Once identifying the most likely class 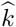, we then estimate the underlying evolutionary parameters ***σ*** = [*σ*_1_, *σ*_2_, …, *σ*_*q*_]^*T*^ that gave rise to patterns within the genomic region provided that it was estimated to be non-neutral, where *σ*_1_, *σ*_2_, …, *σ*_*q*_ represent the *q* evolutionary parameters we are estimating for class 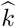.

Consider again the approximated functions 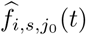 of each summary statistic *s* in observation *i* at detail level *j*_0_. We will use these functions (and as in the preceding section, their associated coefficients) as the independent variables to model multivariate linear regression as

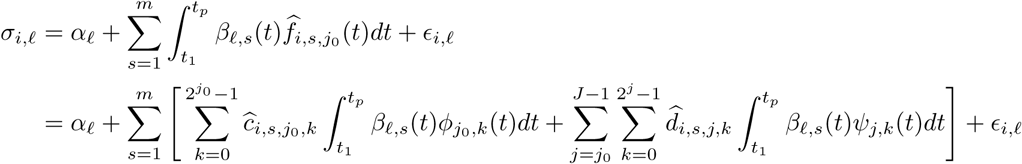

for *ℓ* = 1, 2, …, *q*. Here *i* is the index for the observation number, *σ*_*i,ℓ*_ is the response value for evolutionary parameter *σ*_*ℓ*_ of observation *i, α*_*ℓ*_ is the intercept for evolutionary parameter *σ*_*ℓ*_, *β*_*ℓ,s*_(*t*) is the function for summary statistic *s* of evolutionary parameter *σ*_*ℓ*_, and *ϵ*_*i,ℓ*_ is the error associated with observation *i* of evolutionary parameter *σ*_*ℓ*_. Moreover, define the vector of length *q* containing the evolutionary parameters that generated observation *i* by

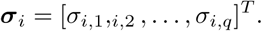

As in the preceding section, to learn the functions *β*_*ℓ,s*_(*t*) we can approximate them using wavelets at a detail level of *j*_0_ as

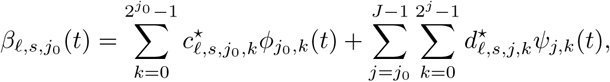

where 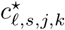 and 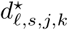 are the coefficients for the father and mother wavelets at scale *j* and location *k* for summary statistic *s* of evolutionary parameter *σ*_*ℓ*_. Denote the vector of length *p* = 2^*J*^ containing father and mother basis coefficients for summary statistics *s* for evolutionary parameter *σ*_*ℓ*_ at detail level *j*_0_ as

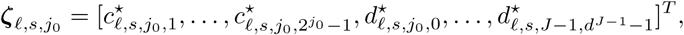

and further define the concatenated vector of length *m* × *p* of such coefficients across all *m* summary statistics for evolutionary parameter *σ*_*ℓ*_ by

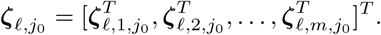

Plugging in this approximation, and using the orthonormality of the set of basis functions, we obtain

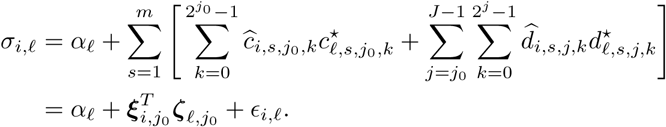

Let ***α*** = [*α*_1_, *α*_2_, …, *α*_*q*_]^*T*^ denote the vector of intercept terms for each of the *q* evolutionary parameters and define the matrix 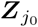 containing *m* × *p* rows and *q* columns by

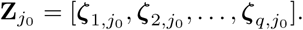

The loss function of the collection of data points 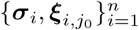 given the set of model parameters 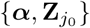 is

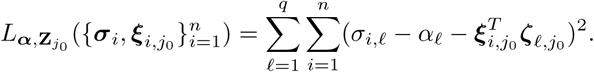

From this loss function, we wish to estimate the intercept terms ***α*** and the coefficients 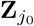. Define 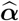 as an estimate of *α* and 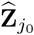 as an estimate of 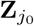. Similarly to the previous section, define

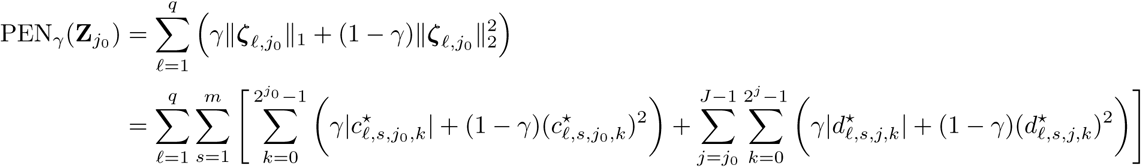

to be the elastic-net penalty (Zou and Hastie, 2005) controlled by parameter *γ* ∈ [0, 1] on the coefficients for the basis functions of the regression coefficient functions, and let *λ* denote a tuning parameter associated with this penalty. We can therefore estimate the coefficient functions as

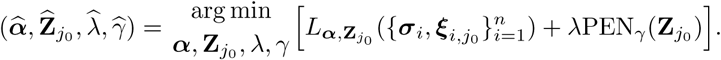

As in preceding section, we perform this estimation, we first learn the underlying functions *f*_*i,s*_(*t*) based on orthonormal wavelet basis functions at detail level *j*_0_,yielding the estimated set of coefficients 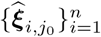 and hence estimated functions 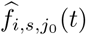. These basis function coefficients are then input as covariates to the penalized regression model, for which ten-fold cross validation is used to estimate the tuning parameter *λ*, the tuning parameter *γ* controlling the elastic-net penalty, and associated parameters ***α*** and 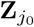. This process is repeated for different detail levels *j*_0_ = 0, 1, …, *J* − 1 to estimate the *j*_0_ that minimizes the ten-fold cross validation error, and the best fitting values of regression model parameters 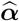 and 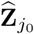 are estimated. These estimates lead to an estimator for the *q* underlying evolutionary parameters for future input data, as well as learned functions 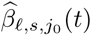 for summary statistic *s, s* = 1, 2, …, *m* of evolutionary parameter *σ*_*ℓ*_, *ℓ* = 1, 2, …, *q*. After parameter inference, evolutionary parameter *σ*_*ℓ*_ is estimated as

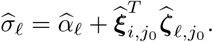

### Calculating summary statistics

Informative summary statistics are likely the most important aspect of developing prediction models. In this manuscript we discuss the use of several sets of summary statistics. For our initial comparison, we utilize a similar set of summary statistics as discussed in Mughal and DeGiorgio (2019) including the mean pairwise sequence difference 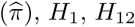, and *H*_2_ */H*_1_. As shown in Mughal and DeGiorgio (2019) *r*^2^ was not informative for classification rates, and for this reason, we omit *r*^2^ as applied in (Mughal and DeGiorgio, 2019) from our model. Moreover, Mughal and DeGiorgio (2019) found that haplotype-based statistics were often more informative than site frequency-based statistics, and for this reason we include the frequencies of the first, second, third, fourth, and fifth most common haplotypes. We also removed all sites with minor allele count less than three because this dramatically reduced the differences between simulated and empirical site frequency spectra (Figure S35). To keep our performance evaluation consistent with the empirical assessment, we also removed these sites for tests with simulations of a constant-size demographic history, although models trained with these simulations were not applied to empirical data.

We calculated each of these *m* = 9 summary statistics in *p* genomic windows across the region of interest, where each window consists of 10 SNPs and overlaps with its neighbors for five SNPs. Because wavelet transformation requires that the number of observations *p* be a power of two, we investigated *p* = 128, leading to 645 SNPs overall used for classification of a genomic region. The classified SNP is the SNP that falls in center of the overlap of windows *p/*2 and *p/*2 + 1, and is taken as the putative location of the site under selection. A schematic illustrating how summary statistics are calculated in *SURFDAWave* is given in Figure S37.

We also explore the use of other statistics that may better differentiate more complex types of selection, such as the set of mean values of *r*^2^ between each window pair. Each one-dimensional statistic (*i.e.*, 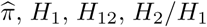, and the frequencies of first to fifth most common haplotypes) is computed using *p* = 128 overlapping windows, of which 64 are non-overlapping. Because the *r*^2^ statistic calculated between each pair of windows is more time consuming to compute than the one-dimensional statistics, we only compute the *r*^2^ statistic between the *p* = 64 non-overlapping 10-SNP windows. In addition to the mean of *r*^2^, we also use the set of values for the variance, skewness, and kurtosis of *r*^2^ computed at each window pair.

### Simulations to test method performance

We designed *SURFDAWave* to learn about adaptation in human populations, so for this reason we focus our simulations on human based parameters. All simulation results use the software SLiM (Haller and Messer, 2017). We use demographic estimates from Terhorst et al. (2017) to model the bottlenecks and expansions experienced by human populations. In addition we also simulate a constant-size demographic history with an effective population size of *N* = 10^4^ (Takahata, 1993) diploid individuals. For all demographic histories we use a mutation rate of 1.25 × 10^−8^ (Scally and Durbin, 2012) and recombination rate drawn from an exponential distribution with mean 3 × 10^−9^ and truncated at three times the mean (Schrider and Kern, 2017) to simulate genomic regions of length two Mb. For selection simulations we let a mutation occur in a generation drawn uniformly at random between 1,020 and 3,000 generations ago, and set this mutation as beneficial with selection strength *s* ∈ [0.005, 0.5] (drawn uniformly at random on a log scale) once it reached frequency *f* ∈ [1*/*(2*N*), 0.1] (drawn uniformly at random on a log scale). This results in our selection simulations containing both hard and soft sweeps. Some combinations of selection parameters are difficult to achieve and for this reason may be under-represented in our simulations (compared to our input parameters) (Figure S38).

To test the performance of *SURFDAWave* on more complex selection scenarios we simulate adaptive introgression. To do this, we simulate a single population that splits into two populations (a recipient and a donor) at a time randomly selected between 13,000 and 32,000 generations ago. This range captures the predicted split times among human, Neanderthal, and Denisovan populations (Kuhlwilm et al., 2016). After allowing the two populations to evolve in isolation, we then simulate a neutral mutation in the center of the two Mb chromosome to occur between 1,020 and 3,000 generations ago in the simulated donor population. Following this, the donor population admixes into the recipient population in which the donor replace between 1 to 10% of the recipient population. After admixture, we treat the simulation as a regular sweep setting and follow the protocol described for sweep simulations.

We also simulated background selection following the protocol described in Schrider and Kern (2017), in which purifying selection is simulated by setting a negative selection coefficient if mutations fall within simulated coding regions. The distribution of coding regions is drawn from both the phastCons (Siepel et al., 2005) and GENCODE (Harrow et al., 2012) databases. Uniformly choosing a random starting point as a SNP in the human genome, we simulate 10^3^, two Mb chromosomes with 75% of mutations falling within coding regions to have a selection coefficient drawn from a gamma distribution with mean −0.0294 with the remaining 25% as neutral, which models the distribution of fitness effects consistent with the human genome (Boyko et al., 2008). As in Mughal and DeGiorgio (2019), we simulated missing data by removing thirty percent of the simulated SNPs in blocks, with each of ten non-overlapping blocks containing 3% of the total data. This process simulates the effects of filters that remove regions of low mappability or alignability (Mallick et al., 2009). To test the accuracy of our prediction models we also simulate 1000 sweeps with background selection.

Because we are unsure of how much training data is required to adequately fit models with our noisy data, we conducted an experiment to see how different numbers of training data points per class affect classification rates (Figure S36). We include in our training data either 1000, 3000, 5000, or 7000 feature vectors for each class (adaptive introgression, neutrality, or sweep). As we increase the number of training data points per class we see an increase in the number of simulations classified correctly for both sweep and adaptive introgression classes. However, we also observe a slight decrease in the number of neutral simulations classified correctly.

### Comparison to *Trendsetter*

Mughal and DeGiorgio (2019) compared the method *Trendsetter* to other recently-developed methods for classifying selective sweeps (Lin et al., 2011; Schrider and Kern, 2016; Kern and Schrider, 2018) and found *Trendsetter* to be more robust to some confounding factors than previous methods, while remaining comparable in power to detect selective sweeps. For this reason, and because *Trendsetter* models the function of importance of summary statistics, we compare the performance of the *SURFDAWave* classifier only to *Trendsetter* in this article. We use the summary statistics 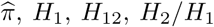, and frequency of the first, second, third, fourth, and fifth most common haplotypes calculated at 128 windows as input data for training both in *Trendsetter* and *SURFDAWave*. Though implementation differs from the original discussed in Mughal and DeGiorgio (2019), we use this new set of statistics (see *Calculating summary statistics*) to directly compare the performances of the two methods. We compare classification rates of *SURFDAWave* using both Haar and Daubechies’ least-asymmetric wavelets with elastic net penalty *γ* chosen through cross validation to *Trendsetter* using both constant (*d* = 1) and linear (*d* = 2) trend penalties.

### Application to empirical data

To locate regions of selection in human genomes, we conducted scans using phased haplotype data from the central European (CEU) and sub-Saharan African Yoruban (YRI) populations in the 1000 Genomes Project dataset (The 1000 Genomes Project Consortium, 2015). Because some genomic regions are difficult to sequence, map, or align, and result in low quality data that is prone to errors, we split the genomes into 100 kb non-overlapping segments, and removed those with mean CRG100 score less than 0.9 (Derrien et al., 2012). As described in *Calculating summary statsitics*, we also removed all sites with minor allele frequency less than three. We then split the remaining data for each chromosome into windows of 10 SNPs where each window overlaps its neighbor for five SNPs, and computed summary statistics discussed in section *Calculating summary statistics* for each window. As we are investigating *p* = 128, each set of statistics for 128 windows comprises a feature vector. When scanning the genome, we shift one window at a time, so that the putative site of selection (the middle SNP falling in the overlap of windows *p/*2 and *p/*2 + 1) will shift by five SNPs each iteration. These feature vectors are used as input to both the *SURFDAWave* classifier and predictor. As we value the correct classification of neutral genomic regions, we use 5000 simulated replicates of each class to train classifiers, because we notice a decrease in the number of correctly classified neutral regions when we use more (Figure S36).

## Acknowledgments

This research was funded by National Institutes of Health grant R35GM128590, National Science Foundation grant DEB-1753489, the Alfred P. Sloan Foundation, a NIGMS funded training grant on Computation, Bioinformatics, and Statistics (Predoctoral Training Program T32GM102057), the NASA Pennsylvania Space Grant Graduate Fellowship, and the Graduate Research Innovation Grant from the Huck Institutes of the Life Sciences. Portions of this research were conducted with Advanced CyberInfrastructure computational resources provided by the Institute for CyberScience at Pennsylvania State University.

