## Supplement for "Learning the properties of adaptive regions with functional data analysis"

### Supplementary materials

Table S1: RMSE values when predicting selection coefficient ($s$), initial frequency ($f$), and time of selection ($T_{\text{sel}}$) for YRI and CEU populations. The values show RMSE measured between standarized log-scaled predicted and actual parameters in simulated data.

| Population | RMSE($s$) | RMSE($f$) | RMSE($T_{\text{sel}}$) |
|---|---|---|---|
| CEU | 0.91 | 0.98 | 0.67 |
| YRI | 0.95 | 0.96 | 0.76 |

Table S2: RMSE values when predicting selection coefficient ($s$), initial frequency ($f$), and time of selection ($T_{\text{sel}}$) for YRI and CEU populations. The values show RMSE measured between log-scaled predicted and actual parameters after unstandarizing.

| Population | RMSE($s$) | RMSE($f$) | RMSE($T_{\text{sel}}$) |
|---|---|---|---|
| CEU | 0.49 | 0.43 | 20.34 |
| YRI | 0.47 | 0.45 | 24.49 |

Table S3: RMSE values when predicting selection coefficient ($s$), initial frequency ($f$), and time of selection ($T_{\text{sel}}$) for YRI and CEU populations tested on simulations of missing data. The values show RMSE measured between standarized log-scaled predicted and actual parameters in simulated data.

| Population | RMSE($s$) | RMSE($f$) | RMSE($T_{\text{sel}}$) |
|---|---|---|---|
| CEU | 0.93 | 1.03 | 0.87 |
| YRI | 1.12 | 1.11 | 1.21 |

Table S4: RMSE values when predicting selection coefficient ($s$), initial frequency ($f$), and time of selection ($T_{\text{sel}}$) for YRI and CEU populations tested on simulations of missing data. The values show RMSE measured between log-scaled predicted and actual parameters after unstandarizing.

| Population | RMSE($s$) | RMSE($f$) | RMSE($T_{\text{sel}}$) |
|---|---|---|---|
| CEU | 0.51 | 0.49 | 23.82 |
| YRI | 0.56 | 0.50 | 37.07 |

Table S5: RMSE values when predicting selection coefficient ($s$), initial frequency ($f$), and time of selection ($T_{\text{sel}}$) for YRI and CEU populations when tested with models trained with simulations of the opposite demography. The values show RMSE measured between standarized log-scaled predicted and actual parameters in simulated data.

| Population | RMSE($s$) | RMSE($f$) | RMSE($T_{\text{sel}}$) |
|---|---|---|---|
| CEU | 0.94 | 1.06 | 0.92 |
| YRI | 0.99 | 1.18 | 1.12 |

Table S6: RMSE values when predicting selection coefficient ($s$), initial frequency ($f$), and time of selection ($T_{\text{sel}}$) for YRI and CEU populations when tested with models trained with simulations of the opposite demography. The values show RMSE measured between log-scaled predicted and actual parameters after unstandarizing.

| Population | RMSE($s$) | RMSE($f$) | RMSE($T_{\text{sel}}$) |
| --- | --- | --- | --- |
| CEU | 0.50 | 0.51 | 26.8 |
| YRI | 0.51 | 0.53 | 34.86 |

Table S7: RMSE values when predicting selection coefficient ($s$), initial frequency ($f$), and time of selection ($T_{\mathrm{sel}}$) for YRI and CEU populations when tested with simulations of selective sweeps plus background selection. The values show RMSE measured between standarized log-scaled predicted and actual parameters in simulated data.

| Population | RMSE($s$) | RMSE($f$) | RMSE($T_{\mathrm{sel}}$) |
|---|---|---|---|
| CEU | 0.87 | 01.3 | 0.65 |
| YRI | 0.96 | 0.99 | 0.71 |

Table S8: RMSE values when predicting selection coefficient ($s$), initial frequency ($f$), and time of selection ($T_{\text{sel}}$) for YRI and CEU populations when tested with simulations of selective sweeps plus background selection. The values show RMSE measured between log-scaled predicted and actual parameters after unstandarizing.

| Population | RMSE($s$) | RMSE($f$) | RMSE($T_{\text{sel}}$) |
|---|---|---|---|
| CEU | 0.36 | 0.51 | 20.23 |
| YRI | 0.45 | 0.49 | 22.10 |

Table S9: RMSE values when predicting selection coefficient ($s$), initial frequency ($f$), and time of donor-recipient split ($T_{\text{split}}$) under adaptive introgression scenarios for YRI and CEU populations. The values show RMSE measured between standarized log-scaled predicted and actual parameters in simulated data.

| Population | RMSE($s$) | RMSE($f$) | RMSE($T_{\text{split}}$) |
|---|---|---|---|
| CEU | 0.93 | 1.00 | 0.98 |
| YRI | 0.97 | 0.91 | 0.99 |

Table S10: Classification of CEU data with classifier trained to differentiate sweeps and neutrality, $\gamma = 1$, Level 1 chosen through cross validation, Daubechies' least asymmetic wavelets

| Chromosome | Neutral | Sweep | $\mathbb{P}$[Sweep] > 0.7 |
|---|---|---|---|
| 1 | 88.0 | 12.0 | 6.9 |
| 2 | 85.8 | 14.2 | 7.8 |
| 3 | 88.8 | 11.2 | 6.3 |
| 4 | 86.6 | 13.4 | 8.5 |
| 5 | 90.2 | 9.8 | 5.1 |
| 6 | 89.8 | 10.2 | 5.3 |
| 7 | 87.5 | 12.5 | 7.0 |
| 8 | 85.7 | 14.3 | 7.7 |
| 9 | 83.9 | 16.1 | 8.9 |
| 10 | 87.2 | 12.8 | 6.9 |
| 11 | 88.4 | 11.6 | 7.2 |
| 12 | 86.2 | 13.8 | 7.9 |
| 13 | 91.6 | 8.4 | 4.2 |
| 14 | 86.4 | 13.6 | 7.7 |
| 15 | 79.4 | 20.6 | 11.8 |
| 16 | 80.6 | 19.4 | 11.0 |
| 17 | 86.4 | 13.6 | 7.5 |
| 18 | 85.1 | 14.9 | 8.7 |
| 19 | 79.4 | 20.6 | 10.7 |
| 20 | 82.8 | 17.2 | 10.0 |
| 21 | 89.4 | 10.6 | 6.2 |
| 22 | 84.4 | 15.6 | 6.5 |

Table S11: Classification of YRI data with classifier trained to differentiate sweeps and neutrality, $\gamma = 1$, Level 1 chosen through cross validation, Daubechies' least asymmetic wavelets

| Chromosome | Neutral | Sweep | $\mathbb{P}$[Sweep] > 0.7 |
|---|---|---|---|
| 1 | 97.1 | 2.9 | 2.0 |
| 2 | 97.8 | 2.2 | 1.4 |
| 3 | 98.2 | 1.8 | 1.1 |
| 4 | 96.6 | 3.4 | 2.0 |
| 5 | 97.2 | 2.8 | 1.7 |
| 6 | 95.2 | 4.8 | 3.4 |
| 7 | 96.1 | 3.9 | 2.5 |
| 8 | 97.5 | 2.5 | 1.7 |
| 9 | 97.2 | 2.8 | 1.9 |
| 10 | 96.9 | 3.1 | 1.9 |
| 11 | 96.7 | 3.3 | 1.8 |
| 12 | 97.4 | 2.6 | 1.9 |
| 13 | 98.2 | 1.8 | 0.9 |
| 14 | 97.2 | 2.8 | 1.3 |
| 15 | 97.9 | 2.1 | 1.3 |
| 16 | 95.8 | 4.2 | 3.0 |
| 17 | 97.7 | 2.3 | 1.5 |
| 18 | 98.2 | 1.8 | 1.1 |
| 19 | 94.3 | 5.7 | 3.8 |
| 20 | 97.5 | 2.5 | 2.0 |
| 21 | 97.6 | 2.4 | 1.2 |
| 22 | 98.5 | 1.5 | 1.1 |

Table S12: Classification of CEU data with classifier trained to differentiate adaptive introgression, sweeps, and neutrality, $\gamma = 1$, Level 1 chosen through cross validation, Daubechies' least asymmetic wavelets

| Chromosome | Neutral | Introgression sweep | Sweep | $\mathbb{P}$[Introgression sweep] > 0.6 | $\mathbb{P}$[Sweep] > 0.6 |
|---|---|---|---|---|---|
| 1 | 87.4 | 4.0 | 8.6 | 0.3 | 1.6 |
| 2 | 85.7 | 4.7 | 9.6 | 0.4 | 0.4 |
| 3 | 88.4 | 3.9 | 7.8 | 0.3 | 0.5 |
| 4 | 85.2 | 6.3 | 8.5 | 0.9 | 1.1 |
| 5 | 90.2 | 2.7 | 7.1 | 0.1 | 0.5 |
| 6 | 88.9 | 3.8 | 7.4 | 0.2 | 0.6 |
| 7 | 87.8 | 3.1 | 9.2 | 0.1 | 0.6 |
| 8 | 84.4 | 5.6 | 10.0 | 0.2 | 1.2 |
| 9 | 85.2 | 3.5 | 11.2 | 0.2 | 0.7 |
| 10 | 86.7 | 5.1 | 8.2 | 0.5 | 0.7 |
| 11 | 88.5 | 3.3 | 8.1 | 0.1 | 0.9 |
| 12 | 86.8 | 3.3 | 9.9 | 0.3 | 0.4 |
| 13 | 91.4 | 3.3 | 5.3 | 0.0 | 0.3 |
| 14 | 86.9 | 3.2 | 9.9 | 0.0 | 0.5 |
| 15 | 79.4 | 5.7 | 14.9 | 0.3 | 1.6 |
| 16 | 81.6 | 5.0 | 13.4 | 0.2 | 2.1 |
| 17 | 87.0 | 5.2 | 7.8 | 0.2 | 0.3 |
| 18 | 85.3 | 5.8 | 8.9 | 0.1 | 0.9 |
| 19 | 82.5 | 4.0 | 13.5 | 0.0 | 0.9 |
| 20 | 83.6 | 4.5 | 11.9 | 0.0 | 0.6 |
| 21 | 88.3 | 6.2 | 5.6 | 0.2 | 0.1 |
| 22 | 86.1 | 2.9 | 11.0 | 0.0 | 0.1 |

Table S13: Classification of CEU data with classifier trained to differentiate adaptive introgression, sweeps, and neutrality, $\gamma = 1$, Level 1 chosen through cross validation, Daubechies' least asymmetic wavelets, including two-dimensional statistics

| Chromosome | Neutral | Introgression sweep | Sweep | $\mathbb{P}$[Introgression sweep] $> 0.6$ | $\mathbb{P}$[Sweep] $> 0.6$ |
|---|---|---|---|---|---|
| 1 | 90.5 | 3.5 | 6.1 | 0.5 | 2.3 |
| 2 | 90.3 | 4.7 | 5.1 | 1.0 | 1.4 |
| 3 | 91.4 | 4.3 | 4.3 | 0.9 | 1.1 |
| 4 | 87.4 | 6.0 | 6.6 | 1.5 | 2.1 |
| 5 | 93.2 | 3.4 | 3.4 | 0.7 | 0.7 |
| 6 | 92.8 | 2.9 | 4.3 | 0.6 | 1.3 |
| 7 | 92.8 | 2.7 | 4.4 | 0.4 | 0.9 |
| 8 | 88.0 | 4.9 | 7.0 | 0.8 | 1.7 |
| 9 | 90.9 | 3.8 | 5.3 | 0.7 | 1.1 |
| 10 | 92.0 | 3.9 | 4.2 | 1.1 | 0.8 |
| 11 | 92.2 | 3.8 | 4.0 | 0.6 | 1.3 |
| 12 | 92.2 | 2.7 | 5.1 | 0.6 | 1.1 |
| 13 | 93.7 | 2.9 | 3.3 | 0.3 | 0.3 |
| 14 | 91.3 | 3.7 | 5.0 | 0.7 | 1.9 |
| 15 | 89.4 | 4.7 | 5.9 | 0.8 | 1.4 |
| 16 | 93.0 | 3.1 | 3.9 | 0.3 | 0.7 |
| 17 | 93.4 | 3.0 | 3.6 | 0.4 | 0.8 |
| 18 | 90.1 | 5.5 | 4.4 | 1.4 | 0.8 |
| 19 | 89.9 | 3.2 | 6.9 | 0.3 | 1.2 |
| 20 | 90.3 | 4.6 | 5.1 | 0.3 | 0.9 |
| 21 | 90.9 | 4.4 | 4.7 | 0.7 | 0.8 |
| 22 | 94.4 | 3.8 | 1.8 | 0.3 | 0.0 |

Table S14: Classification of YRI data with classifier trained to differentiate adaptive introgression, sweeps, and neutrality, $\gamma = 1$, Level 1 chosen through cross validation, Daubechies' least asymmetic wavelets

| Chromosome | Neutral | Introgression sweep | Sweep | $\mathbb{P}$[Introgression sweep] > 0.6 | $\mathbb{P}$[Sweep] > 0.6 |
|---|---|---|---|---|---|
| 1 | 96.9 | 0.3 | 2.8 | 0.1 | 0.7 |
| 2 | 96.9 | 0.1 | 2.9 | 0.0 | 0.8 |
| 3 | 97.7 | 0.3 | 2.0 | 0.0 | 0.6 |
| 4 | 96.3 | 0.5 | 3.2 | 0.2 | 0.6 |
| 5 | 96.6 | 0.2 | 3.2 | 0.0 | 1.0 |
| 6 | 95.5 | 1.8 | 2.8 | 0.8 | 0.5 |
| 7 | 96.1 | 0.3 | 3.6 | 0.0 | 1.0 |
| 8 | 97.3 | 0.1 | 2.7 | 0.0 | 0.8 |
| 9 | 96.8 | 0.3 | 2.9 | 0.1 | 0.5 |
| 10 | 97.0 | 0.7 | 2.3 | 0.0 | 0.3 |
| 11 | 96.5 | 0.4 | 3.1 | 0.0 | 0.9 |
| 12 | 97.3 | 0.2 | 2.4 | 0.0 | 0.7 |
| 13 | 98.2 | 0.3 | 1.6 | 0.0 | 0.3 |
| 14 | 96.9 | 0.1 | 3.0 | 0.0 | 0.9 |
| 15 | 97.5 | 0.0 | 2.5 | 0.0 | 0.9 |
| 16 | 95.4 | 0.7 | 3.9 | 0.2 | 0.9 |
| 17 | 97.7 | 0.7 | 1.6 | 0.2 | 0.3 |
| 18 | 97.9 | 0.6 | 1.5 | 0.0 | 0.4 |
| 19 | 94.8 | 0.8 | 4.4 | 0.0 | 1.2 |
| 20 | 97.5 | 0.6 | 1.9 | 0.0 | 0.5 |
| 21 | 98.5 | 0.4 | 1.2 | 0.0 | 0.0 |
| 22 | 98.3 | 0.6 | 1.1 | 0.0 | 0.2 |

Table S15: Classification of YRI data with classifier trained to differentiate adaptive introgression, sweeps, and neutrality, $\gamma = 1$, Level 1 chosen through cross validation, Daubechies' least asymmetic wavelets, including two-dimensinoal statistics

| Chromosome | Neutral | Introgression sweep | Sweep | $\mathbb{P}$[Introgression sweep] $> 0.6$ | $\mathbb{P}$[Sweep] $> 0.6$ |
|---|---|---|---|---|---|
| 1 | 95.8 | 0.5 | 3.7 | 0.0 | 0.8 |
| 2 | 96.9 | 0.3 | 2.8 | 0.0 | 0.3 |
| 3 | 97.4 | 0.4 | 2.2 | 0.1 | 0.8 |
| 4 | 96.2 | 0.9 | 2.9 | 0.1 | 0.4 |
| 5 | 96.7 | 0.3 | 2.9 | 0.0 | 0.7 |
| 6 | 95.6 | 2.3 | 2.1 | 1.1 | 0.4 |
| 7 | 96.0 | 0.9 | 3.0 | 0.1 | 0.5 |
| 8 | 96.4 | 0.8 | 2.8 | 0.1 | 0.7 |
| 9 | 97.3 | 0.5 | 2.2 | 0.0 | 0.6 |
| 10 | 97.7 | 0.9 | 1.4 | 0.0 | 0.4 |
| 11 | 96.7 | 0.8 | 2.5 | 0.1 | 0.3 |
| 12 | 96.1 | 0.7 | 3.2 | 0.0 | 0.7 |
| 13 | 98.1 | 0.7 | 1.3 | 0.2 | 0.3 |
| 14 | 97.8 | 0.3 | 1.9 | 0.0 | 0.6 |
| 15 | 97.0 | 0.7 | 2.3 | 0.0 | 0.8 |
| 16 | 96.3 | 1.1 | 2.5 | 0.2 | 0.5 |
| 17 | 97.8 | 0.8 | 1.4 | 0.2 | 0.1 |
| 18 | 97.8 | 0.4 | 1.7 | 0.0 | 0.4 |
| 19 | 94.7 | 2.0 | 3.3 | 0.2 | 0.7 |
| 20 | 97.2 | 1.1 | 1.7 | 0.0 | 0.4 |
| 21 | 97.9 | 0.9 | 1.2 | 0.1 | 0.1 |
| 22 | 98.7 | 1.0 | 0.4 | 0.3 | 0.0 |

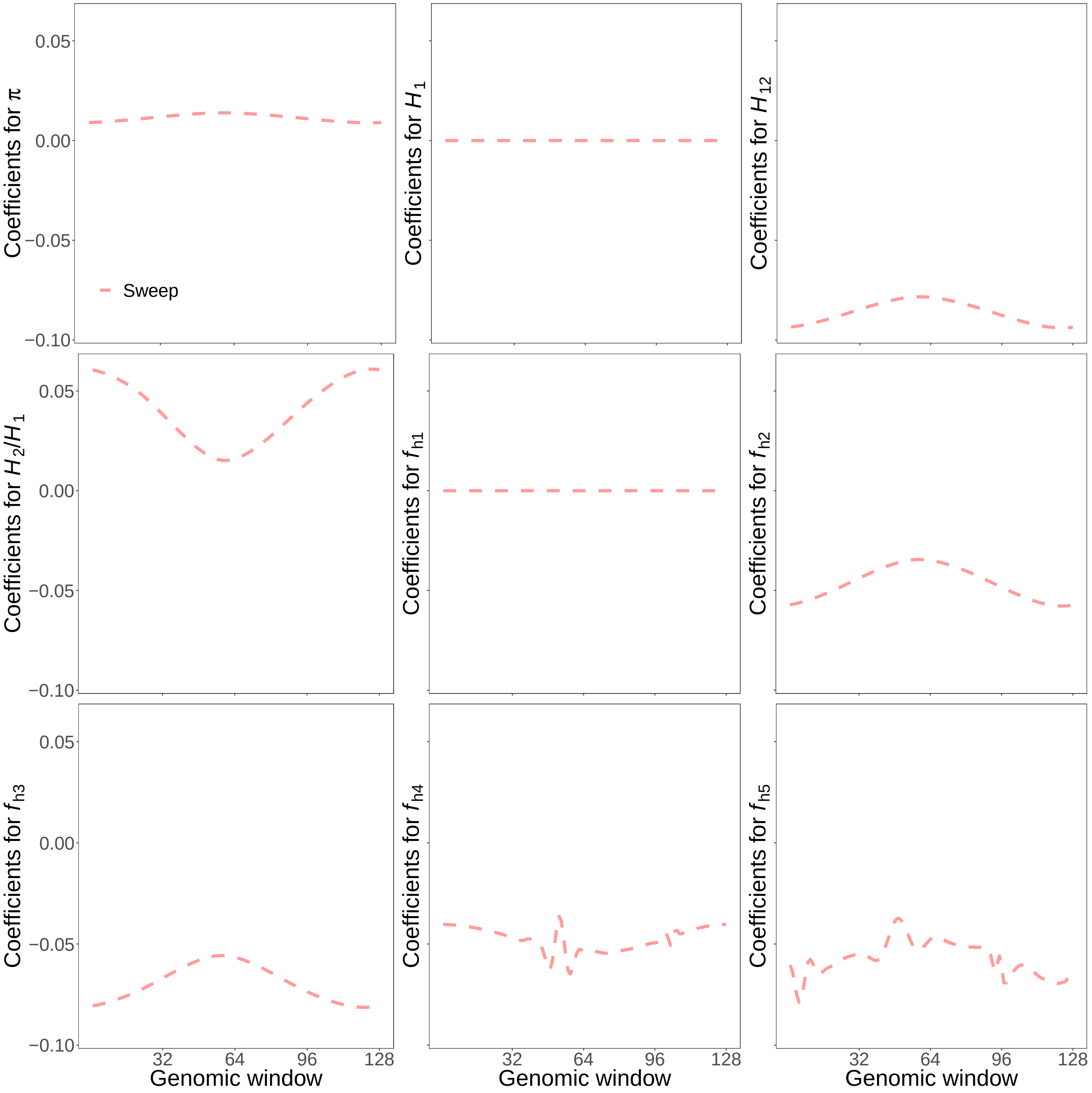


Figure S1: Reconstructed wavelets from regression coefficients ($\beta$s) in sweep versus neutrality scenarios for summary statistics $\widehat{\pi}$, $H_1$, $H_{12}$, $H_2/H_1$, and frequencies of first to fifth most common haplotypes for *SURFDAWave* when $\gamma = 1$. *SURFDAWave* was trained on simulations of scenarios simulated under demographic specifications for European CEU demographic history. Note that the wavelet reconstructions for all summary statistics are plotted on the same scale, thereby making the distributions of some summaries difficult to decipher as their magnitudes are relatively small. *SURFDAWave* results shown are using Daubechies' least-asymmetric wavelets to estimate spatial distributions of summary statistics. Level 1 chosen through cross-validation.

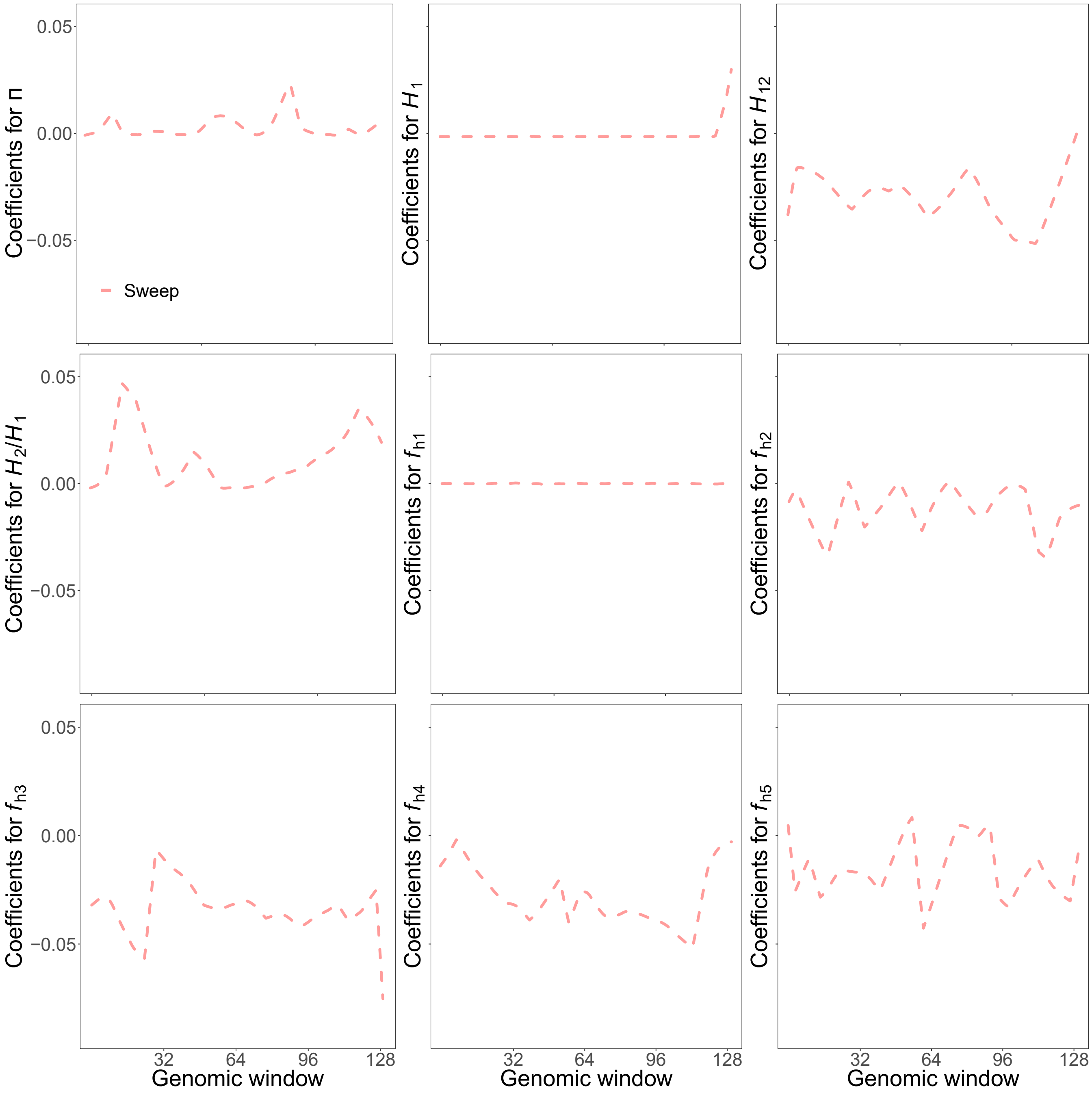


Figure S2: Spatial distribution of regression coefficients ($\beta$s) in sweep scenarios for summary statistics $H_1$, $H_{12}$, $H_2/H_1$, and frequencies of first to sixth most common haplotypes for *Trendsetter* with a linear $d = 2$ trend penalty. *Trendsetter* was trained on simulations of constant demographic history.

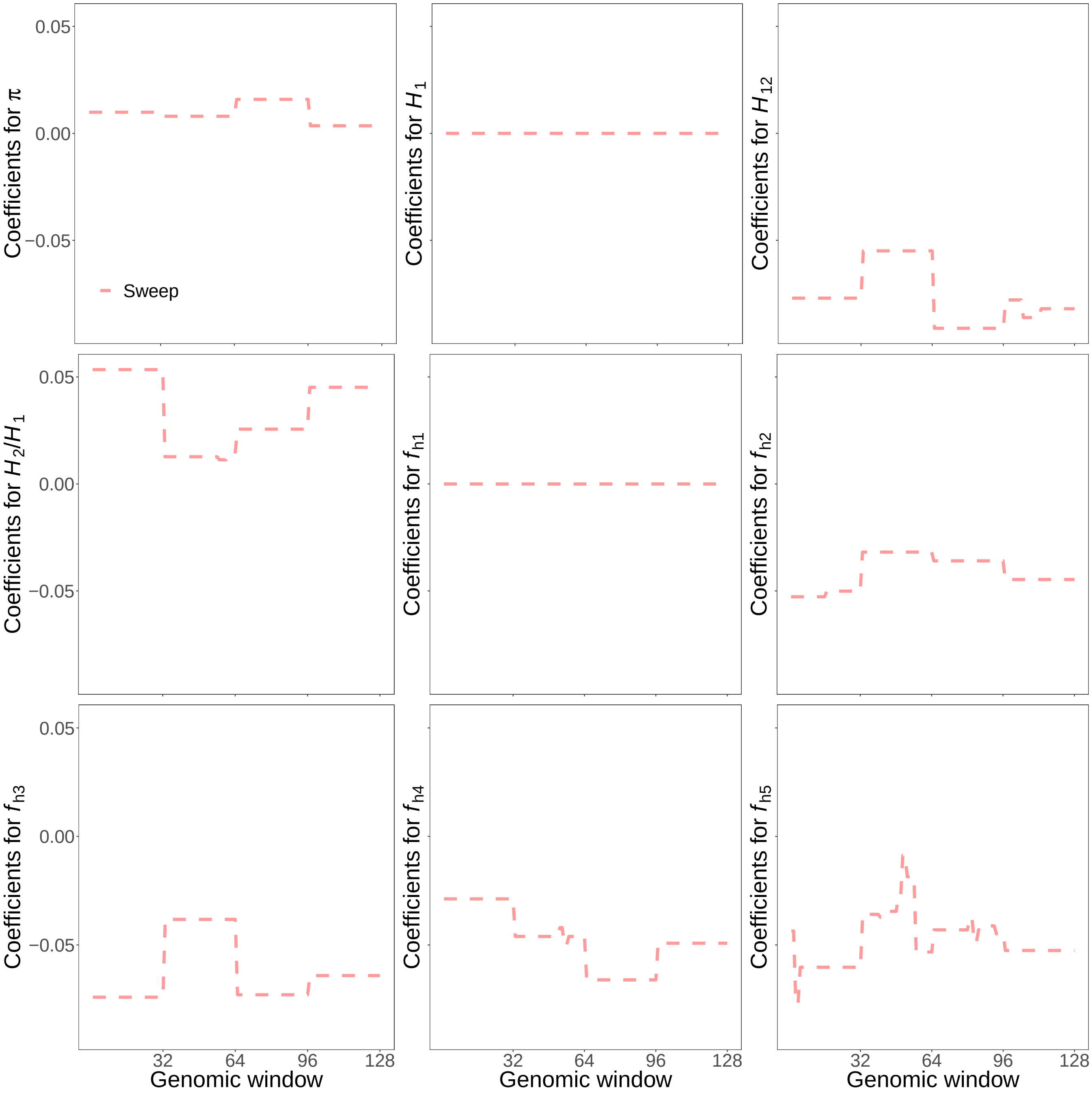


Figure S3: Reconstructed wavelets from regression coefficients ($\beta$s) in sweep versus neutrality scenarios for summary statistics $\widehat{\pi}$, $H_1$, $H_{12}$, $H_2/H_1$, and frequencies of first to fifth most common haplotypes for *SURFDAWave* when $\gamma = 1$. *SURFDAWave* was trained on simulations of scenarios simulated under demographic specifications for European CEU demographic history. Note that the wavelet reconstructions for all summary statistics are plotted on the same scale, thereby making the distributions of some summaries difficult to decipher as their magnitudes are relatively small. *SURFDAWave* results shown are using Haar wavelets to estimate spatial distributions of summary statistics. Level 2 chosen through cross-validation.

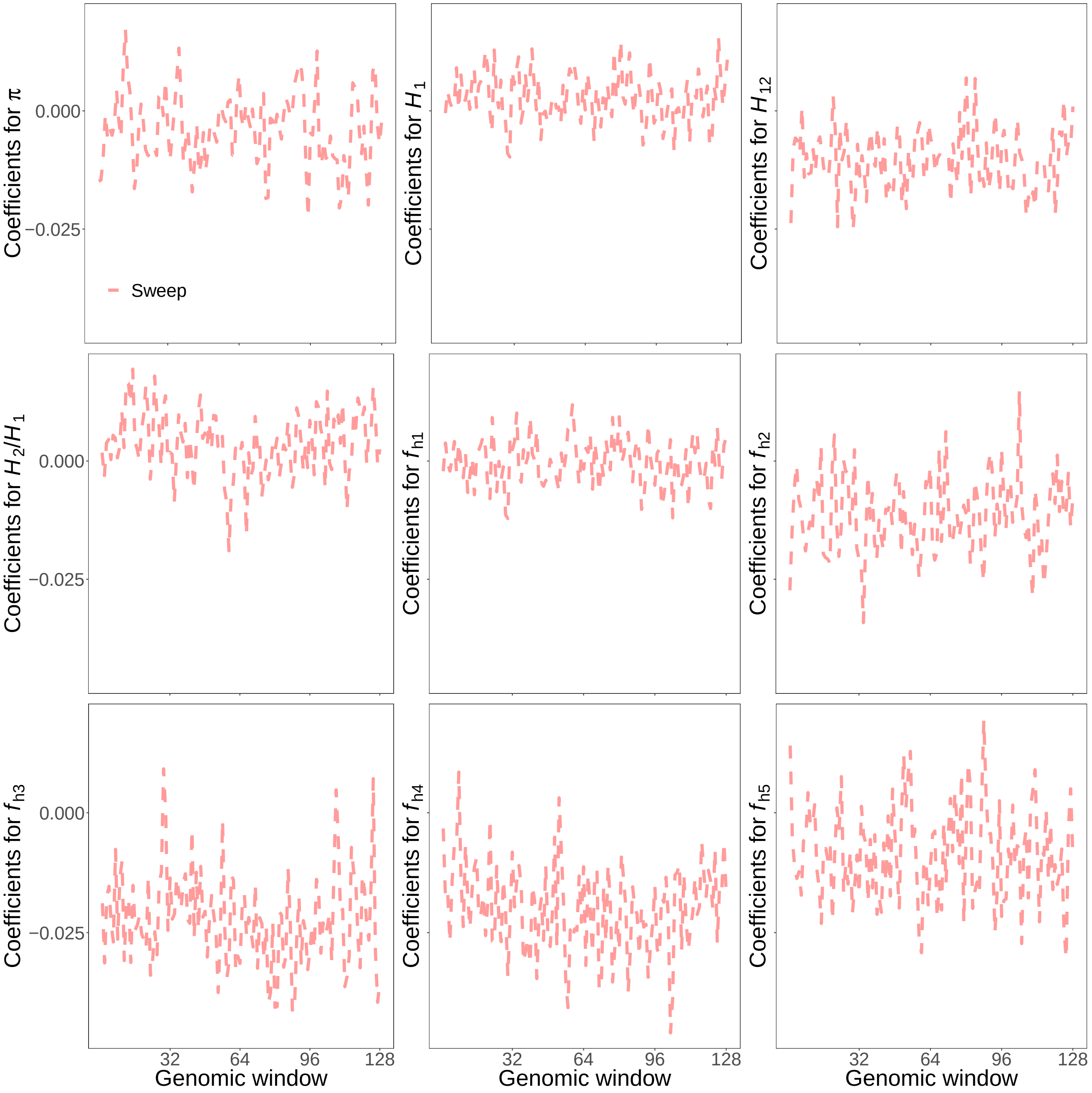


Figure S4: Reconstructed wavelets from regression coefficients ($\beta$s) in sweep versus neutrality scenarios for summary statistics $\widehat{\pi}$, $H_1$, $H_{12}$, $H_2/H_1$, and frequencies of first to fifth most common haplotypes for *SURFDAWave* when $\gamma = 0$. *SURFDAWave* was trained on simulations of scenarios simulated under demographic specifications for European CEU demographic history. Note that the wavelet reconstructions for all summary statistics are plotted on the same scale, thereby making the distributions of some summaries difficult to decipher as their magnitudes are relatively small. *SURFDAWave* results shown are using Daubechies' least-asymmetric wavelets wavelets to estimate spatial distributions of summary statistics. Level 4 chosen through cross-validation.

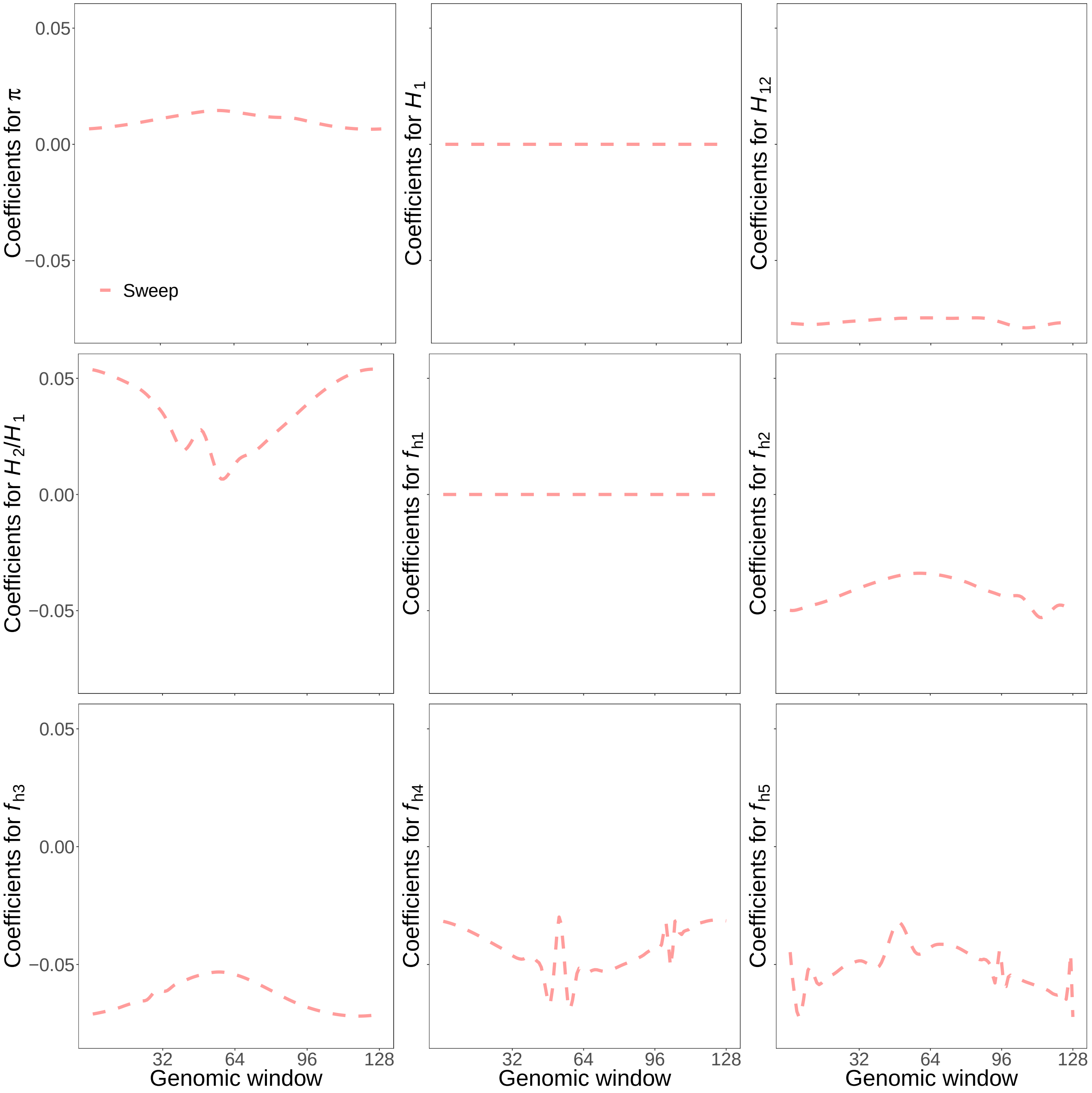


Figure S5: Reconstructed wavelets from regression coefficients ($\beta$s) in sweep versus neutrality scenarios for summary statistics $\widehat{\pi}$, $H_1$, $H_{12}$, $H_2/H_1$, and frequencies of first to fifth most common haplotypes for *SURFDAWave* when $\gamma = 0.7$. *SURFDAWave* was trained on simulations of scenarios simulated under demographic specifications for European CEU demographic history. Note that the wavelet reconstructions for all summary statistics are plotted on the same scale, thereby making the distributions of some summaries difficult to decipher as their magnitudes are relatively small. *SURFDAWave* results shown are using Daubechies' least-asymmetric wavelets wavelets to estimate spatial distributions of summary statistics. Level 4 and $\gamma = 0.7$ chosen through cross-validation.

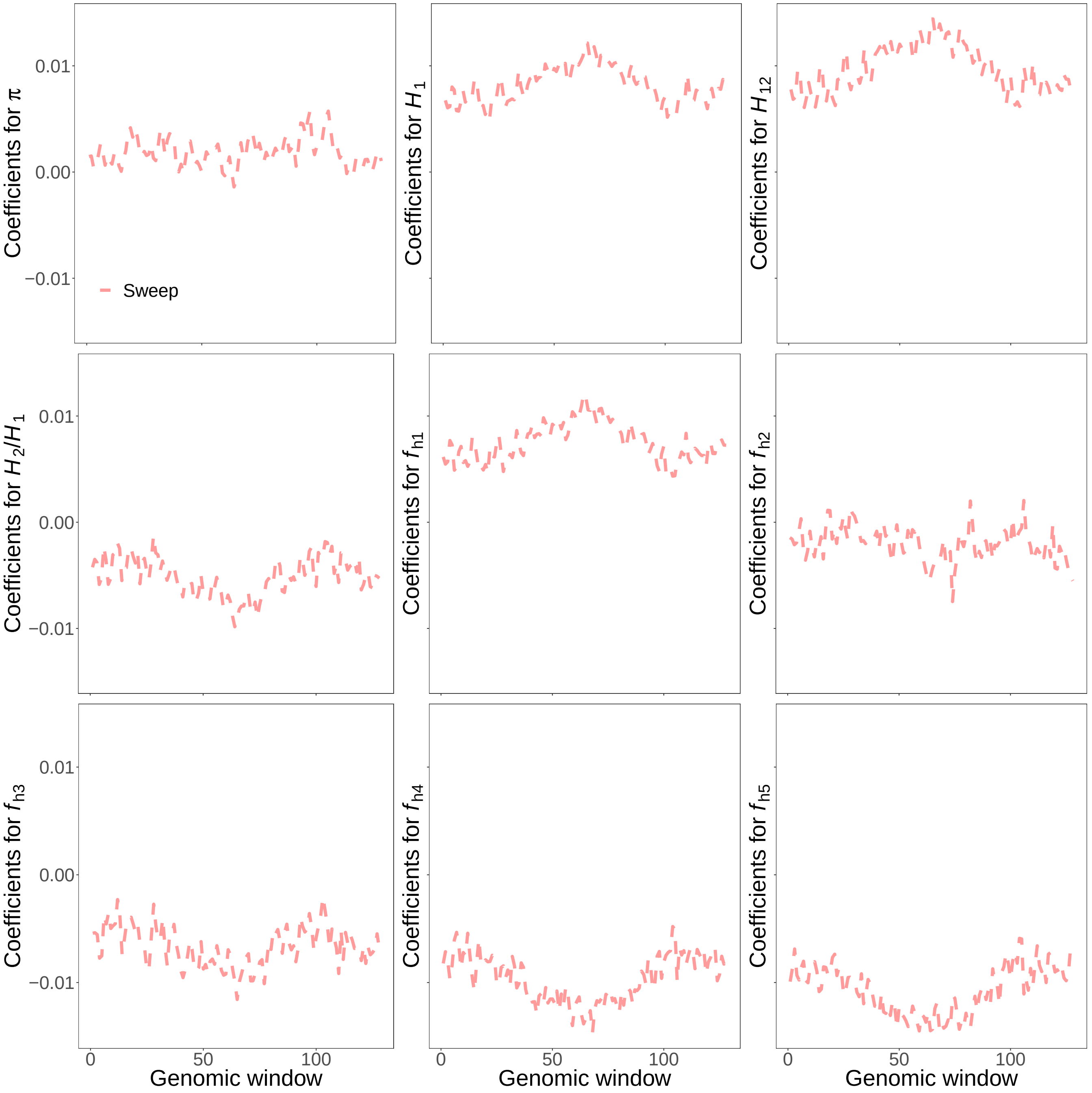


Figure S6: Reconstructed wavelets from regression coefficients ($\beta$s) in sweep versus neutrality scenarios for summary statistics $\widehat{\pi}$, $H_1$, $H_{12}$, $H_2/H_1$, and frequencies of first to fifth most common haplotypes for *SURFDAWave* when $\gamma = 0$. *SURFDAWave* was trained on simulations of scenarios simulated under demographic specifications for sub-Saharan African YRI demographic history. Note that the wavelet reconstructions for all summary statistics are plotted on the same scale, thereby making the distributions of some summaries difficult to decipher as their magnitudes are relatively small. *SURFDAWave* results shown are using Daubechies' least-asymmetric wavelets wavelets to estimate spatial distributions of summary statistics. Level 4 chosen through cross-validation.

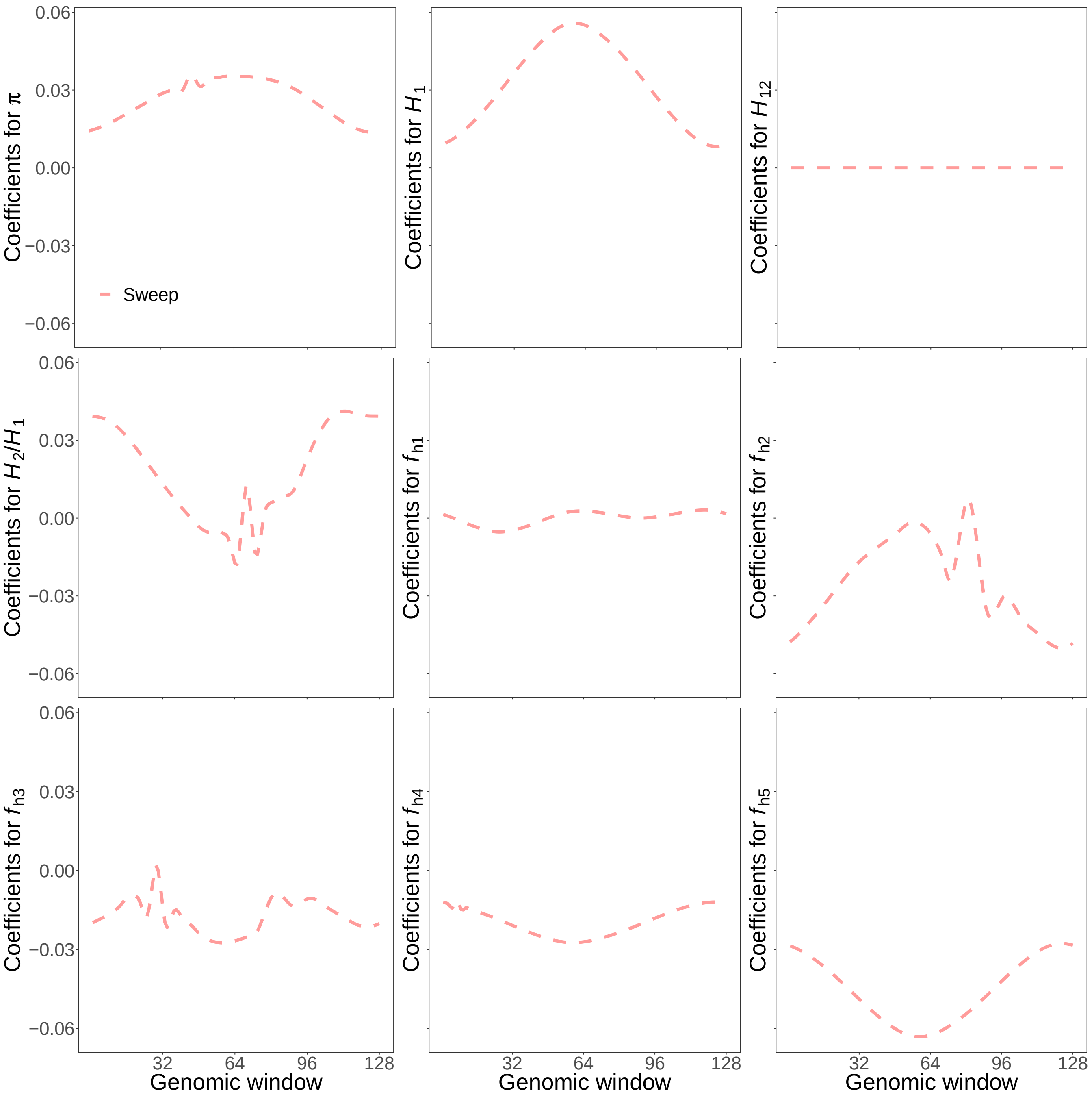


Figure S7: Reconstructed wavelets from regression coefficients ($\beta$s) in sweep versus neutrality scenarios for summary statistics $\widehat{\pi}$, $H_1$, $H_{12}$, $H_2/H_1$, and frequencies of first to fifth most common haplotypes for *SURFDAWave* when $\gamma = 0.5$. *SURFDAWave* was trained on simulations of scenarios simulated under demographic specifications for sub-Saharan African YRI demographic history. Note that the wavelet reconstructions for all summary statistics are plotted on the same scale, thereby making the distributions of some summaries difficult to decipher as their magnitudes are relatively small. *SURFDAWave* results shown are using Daubechies' least-asymmetric wavelets wavelets to estimate spatial distributions of summary statistics. Level 1 and *gamma* chosen through cross-validation.

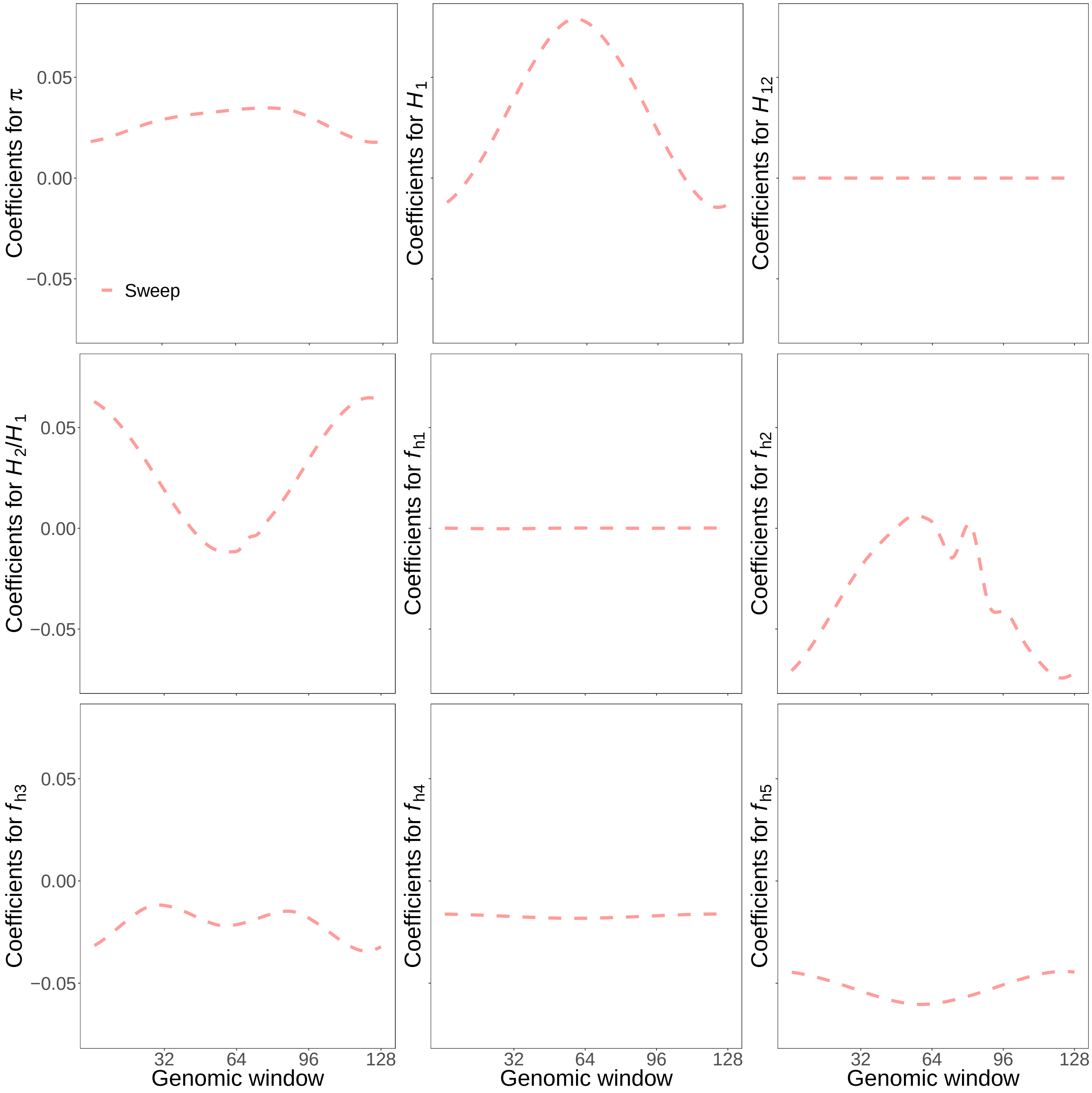


Figure S8: Reconstructed wavelets from regression coefficients ($\beta$s) in sweep versus neutrality scenarios for summary statistics $\widehat{\pi}$, $H_1$, $H_{12}$, $H_2/H_1$, and frequencies of first to fifth most common haplotypes for *SURFDAWave* when $\gamma = 1$. *SURFDAWave* was trained on simulations of scenarios simulated under demographic specifications for sub-Saharan African YRI demographic history. Note that the wavelet reconstructions for all summary statistics are plotted on the same scale, thereby making the distributions of some summaries difficult to decipher as their magnitudes are relatively small. *SURFDAWave* results shown are using Daubechies' least-asymmetric wavelets wavelets to estimate spatial distributions of summmry statistics. Level 1 chosen through cross-validation.

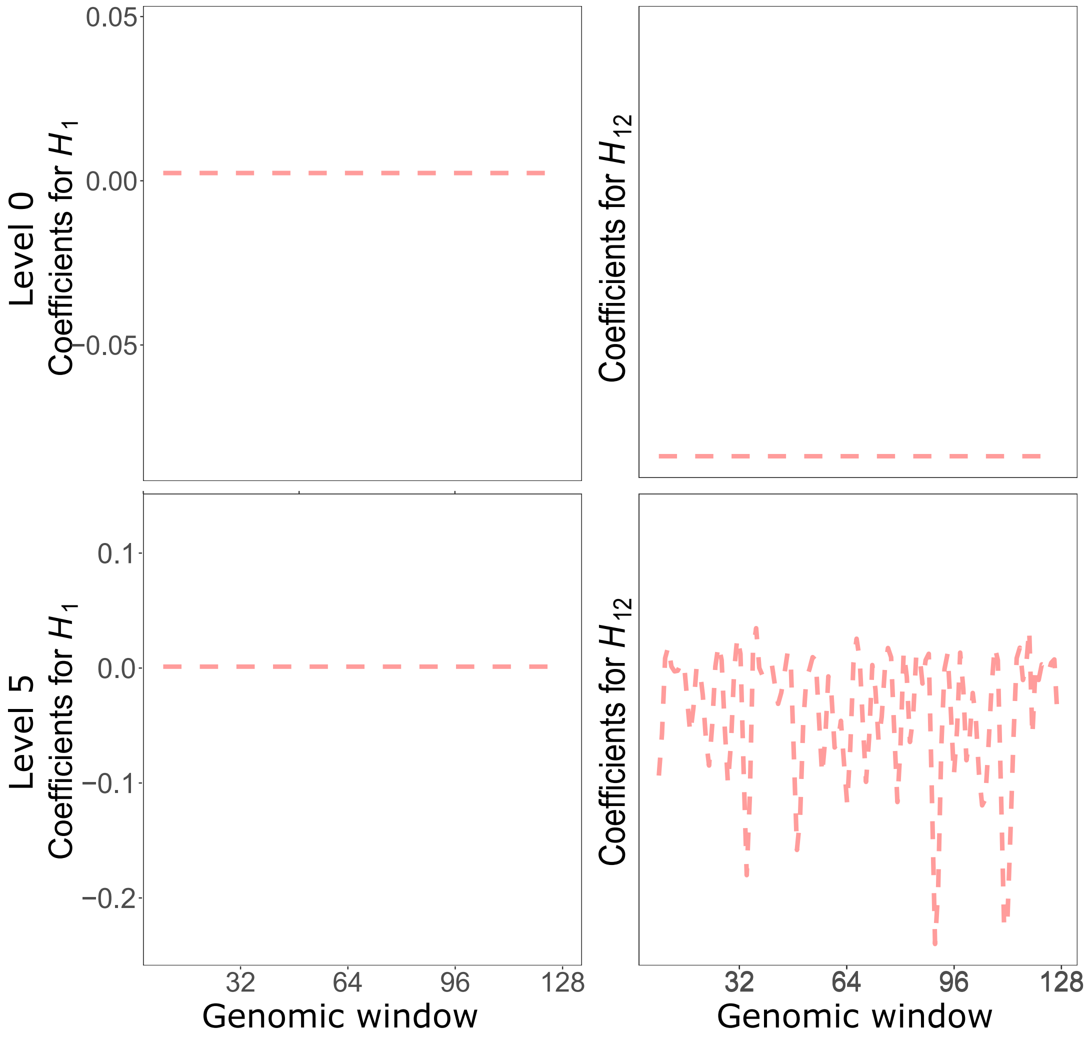


Figure S9: Reconstructed wavelets from regression coefficients ($\beta$s) in sweep vs. neutrality scenarios for summary statistics $H_1$ and $H_{12}$ showing difference between discrete wavelet transform at level 0 and level 5. Using Daubechies' least-Asymmetric wavelets and $\gamma = 1$.

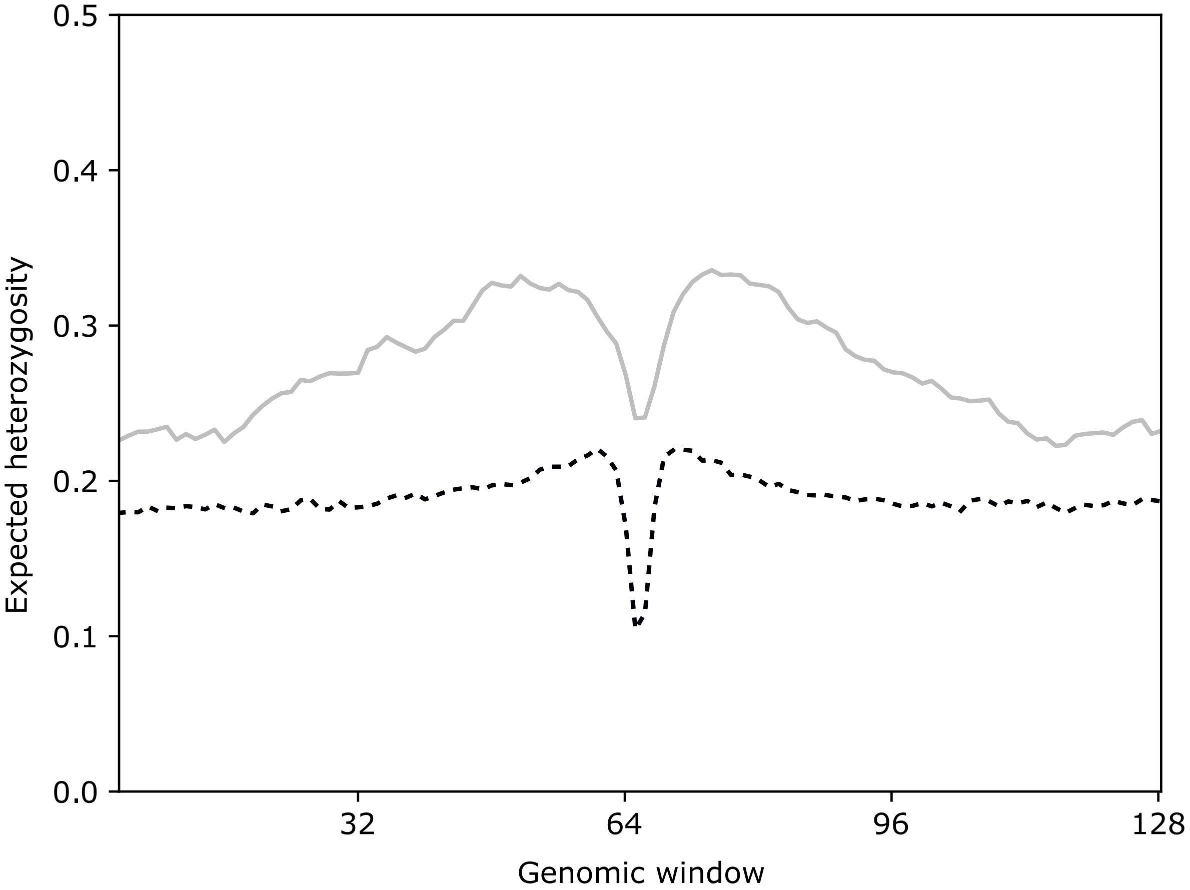


Figure S10: Value of expected heterozygosity across simulated regions of adaptive introgression with varying divergence times. The black dotted line shows the value of the statistic when the divergence time is shorter (30,000 generations ago) and the gray line shows the value when the divergence time is longer (400,000 generations ago).

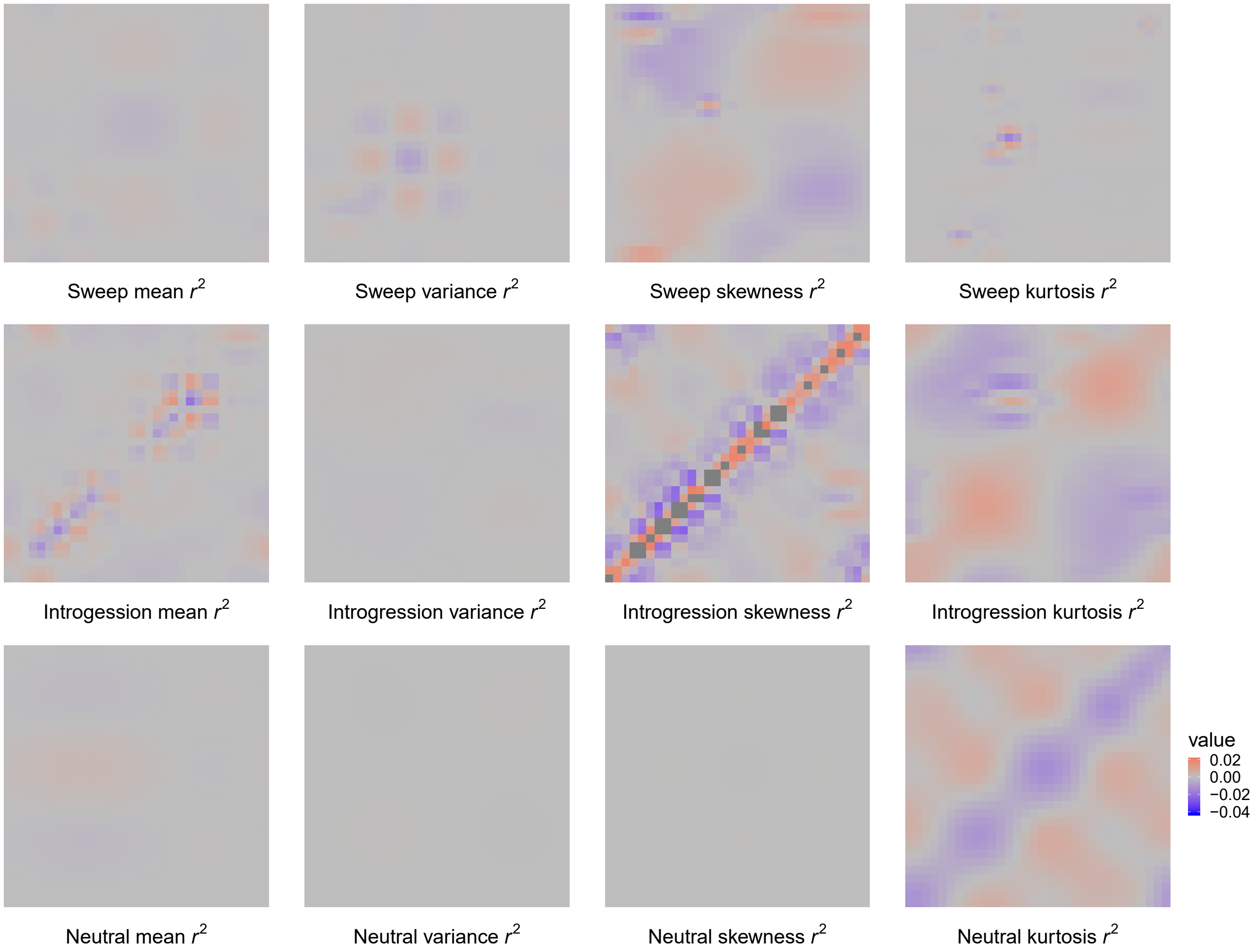


Figure S11: Reconstructed wavelets from regression coefficients ($\beta$s) when differentiating among adaptive introgression, sweeps, and neutrality scenarios for summary statistics mean, variance, skewness, and kurtosis of pairwise $r^2$ for *SURFDAWave* when $\gamma = 1$, when trained with statistics $\widehat{\pi}$, $H_1$, $H_{12}$, $H_2/H_1$, and frequencies of first to fifth most common haplotypes (Figure S13). *SURFDAWave* was trained on simulations of scenarios simulated under demographic specifications for European CEU demographic history. Note that the wavelet reconstructions for all summary statistics are plotted on the same scale, thereby making the distributions of some summaries difficult to decipher as their magnitudes are relatively small. *SURFDAWave* results shown are using Daubechies' least-asymmetric wavelets wavelets to estimate spatial distributions of summary statistics. Level 1 chosen through cross-validation.

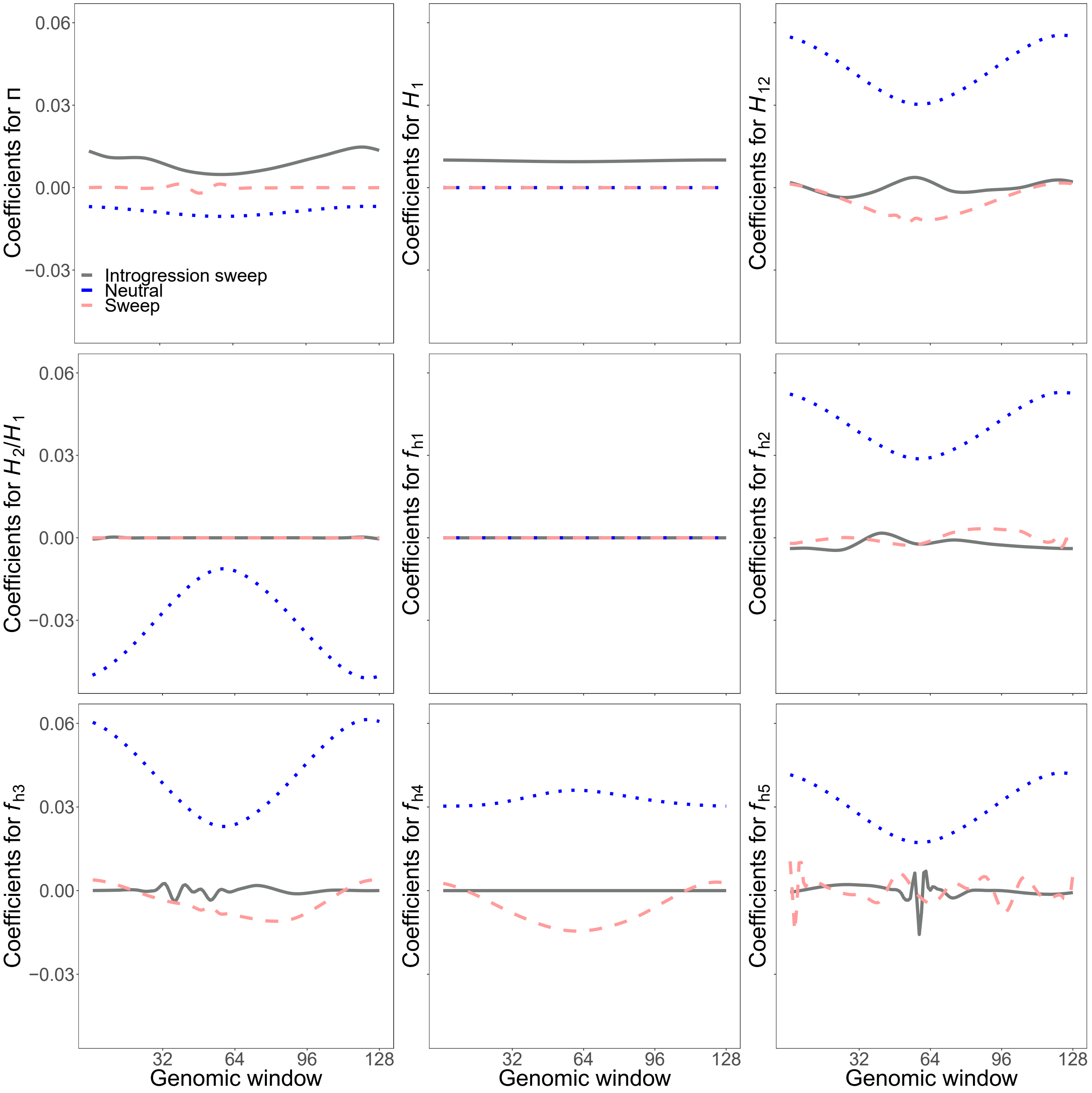


Figure S12: Reconstructed wavelets from regression coefficients ($\beta$s) when differentiating among adaptive introgression, sweeps, and neutrality scenarios for summary statistics $\widehat{\pi}$, $H_1$, $H_{12}$, $H_2/H_1$, and frequencies of first to fifth most common haplotypes for *SURFDAWave* when $\gamma = 1$. *SURFDAWave* was trained on simulations of scenarios simulated under demographic specifications for European CEU demographic history. Note that the wavelet reconstructions for all summary statistics are plotted on the same scale, thereby making the distributions of some summaries difficult to decipher as their magnitudes are relatively small. *SURFDAWave* results shown are using Daubechies' least-asymmetric wavelets wavelets to estimate spatial distributions of summary statistics. Level 1 chosen through cross-validation.

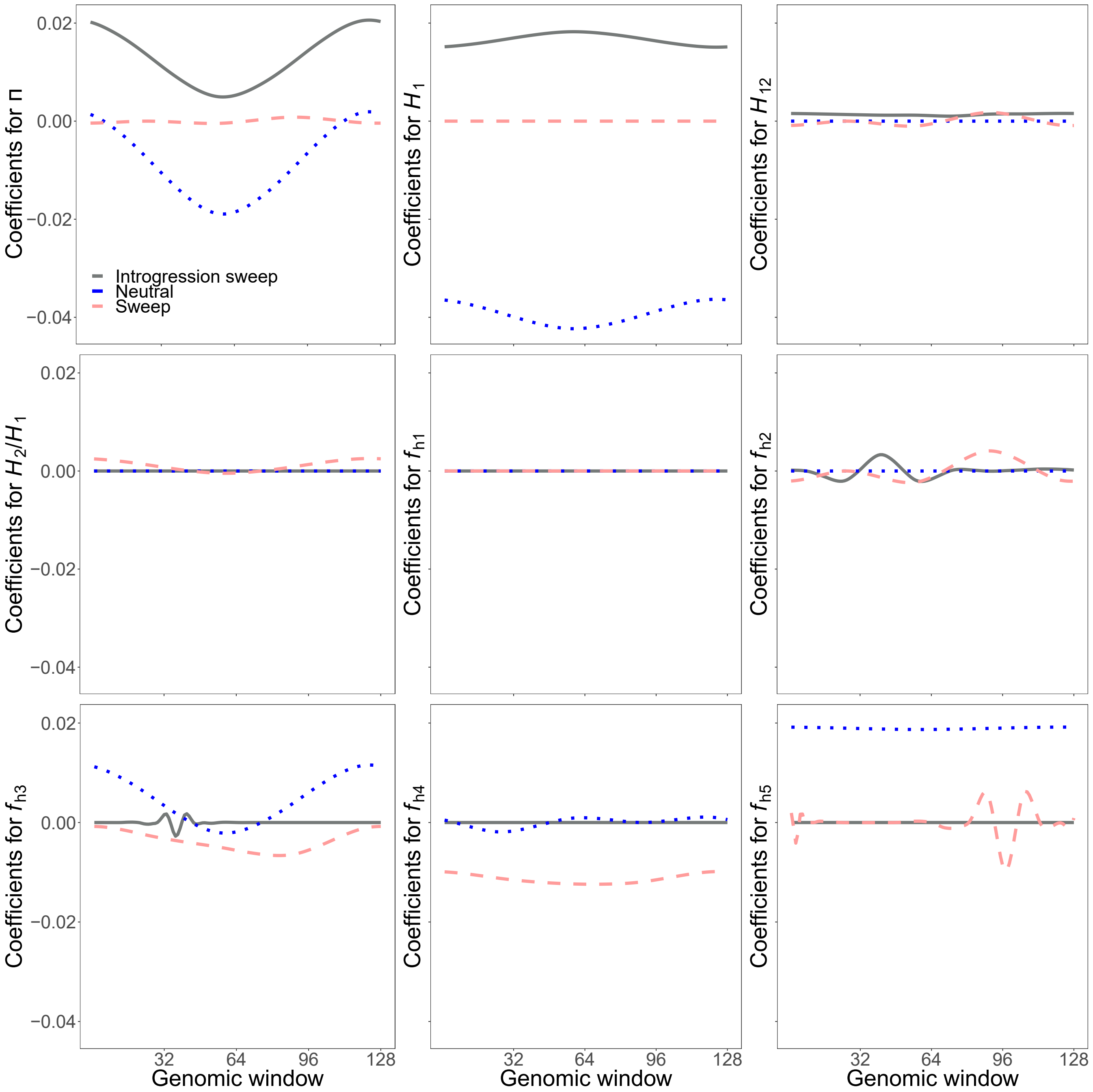


Figure S13: Reconstructed wavelets from regression coefficients ($\beta$s) when differentiating among adaptive introgression, sweeps, and neutrality scenarios for summary statistics $\widehat{\pi}$, $H_1$, $H_{12}$, $H_2/H_1$, and frequencies of first to fifth most common haplotypes for $SURFDAWave$ when $\gamma = 1$, when trained with additional statistics mean, variance, skewness, and kurtosis of pairwise $r^2$ (Figure S11). $SURFDAWave$ was trained on simulations of scenarios simulated under demographic specifications for European CEU demographic history. Note that the wavelet reconstructions for all summary statistics are plotted on the same scale, thereby making the distributions of some summaries difficult to decipher as their magnitudes are relatively small. $SURFDAWave$ results shown are using Daubechies' least-asymmetric wavelets wavelets to estimate spatial distributions of summary statistics. Level 1 chosen through cross-validation.

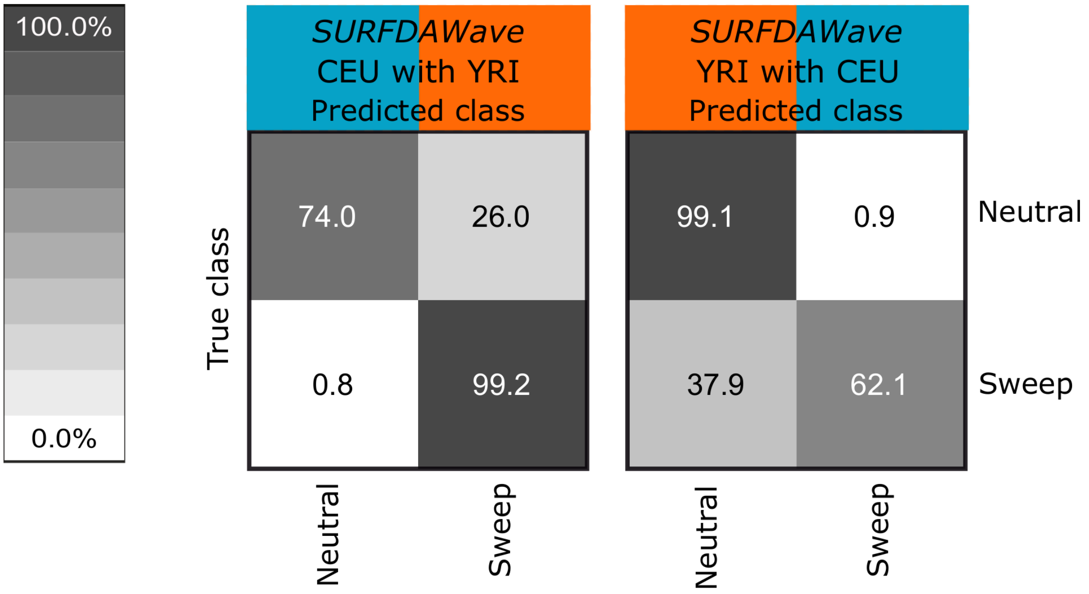


Figure S14: Confusion matrices showing classification results for demographic mis-specificaiton. (Left) Classification rates of simulations conducted under CEU European demographic specifications when the model is trained with simulations conducted under YRI African demographic specifications. (Right) Classification rates of simulations conducted under YRI African demographic specifications when the model is trained with simulations conducted under CEU European demographic specifications.

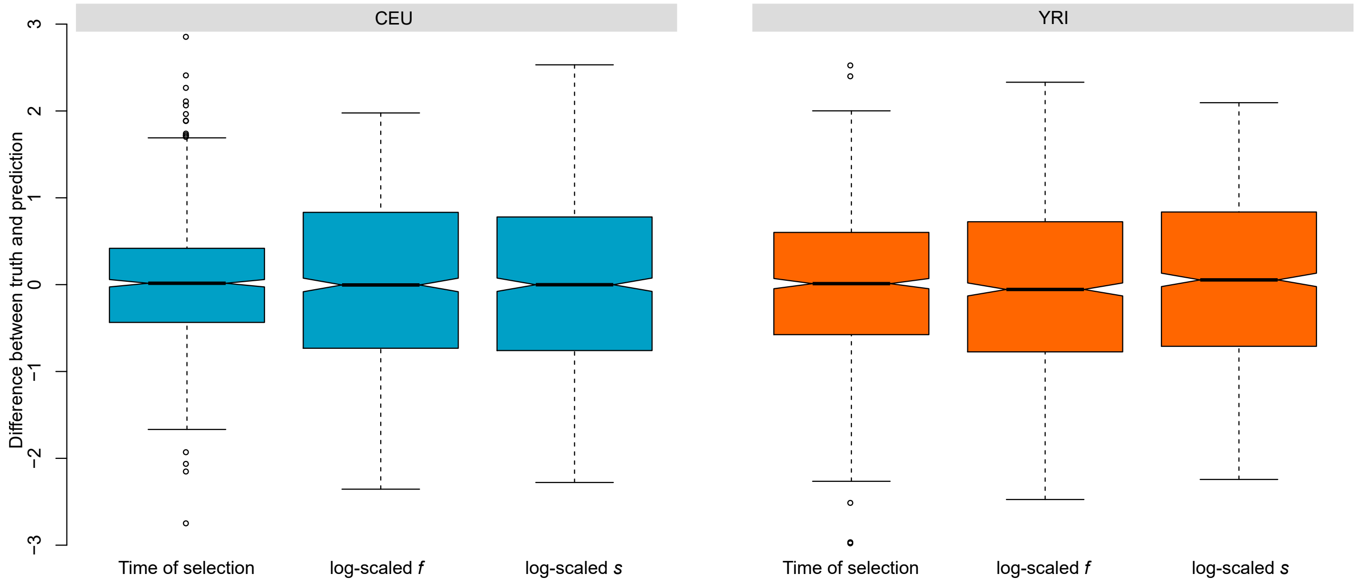


Figure S15: Difference between standardized predicted and actual selection parameters with *SURFDAWave* for the CEU and YRI demographic models. (Left box plot) Difference in prediction and truth of log scaled time at which mutation became beneficial. (Middle box plot) Difference in prediction and truth of log scaled frequency reached by mutation prior to it becoming beneficial ($f$). (Right box plot) Difference in prediction and truth of log scaled selection coefficient ($s$).

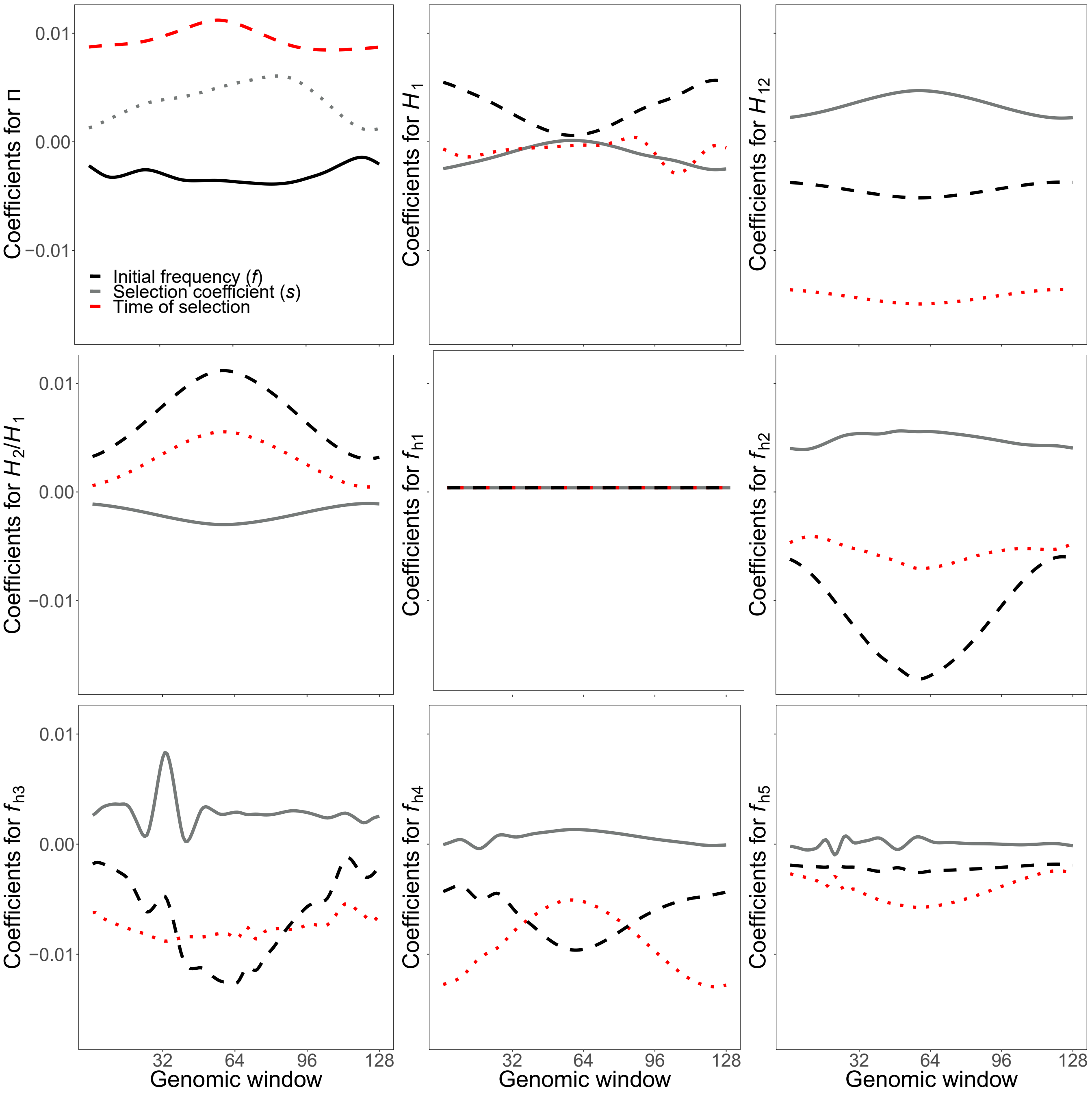


Figure S16: Reconstructed wavelets from regression coefficients ($\beta$s) in predicting time at which mutation became beneficial, frequency reached by mutation before becoming beneficial, and selection strength for summary statistics $\widehat{\pi}$, $H_1$, $H_{12}$, $H_2/H_1$, and frequencies of first to fifth most common haplotypes for *SURFDAWave* when $\gamma = 0.6$. *SURFDAWave* was trained on simulations of scenarios simulated under demographic specifications for European CEU demographic history. Note that the wavelet reconstructions for all summary statistics are plotted on the same scale, thereby making the distributions of some summaries difficult to decipher as their magnitudes are relatively small. *SURFDAWave* results shown are using Daubechies' least-asymmetric wavelets wavelets to estimate spatial distributions of summary statistics. Level 1 and $\gamma = 0.6$ chosen through cross-validation.

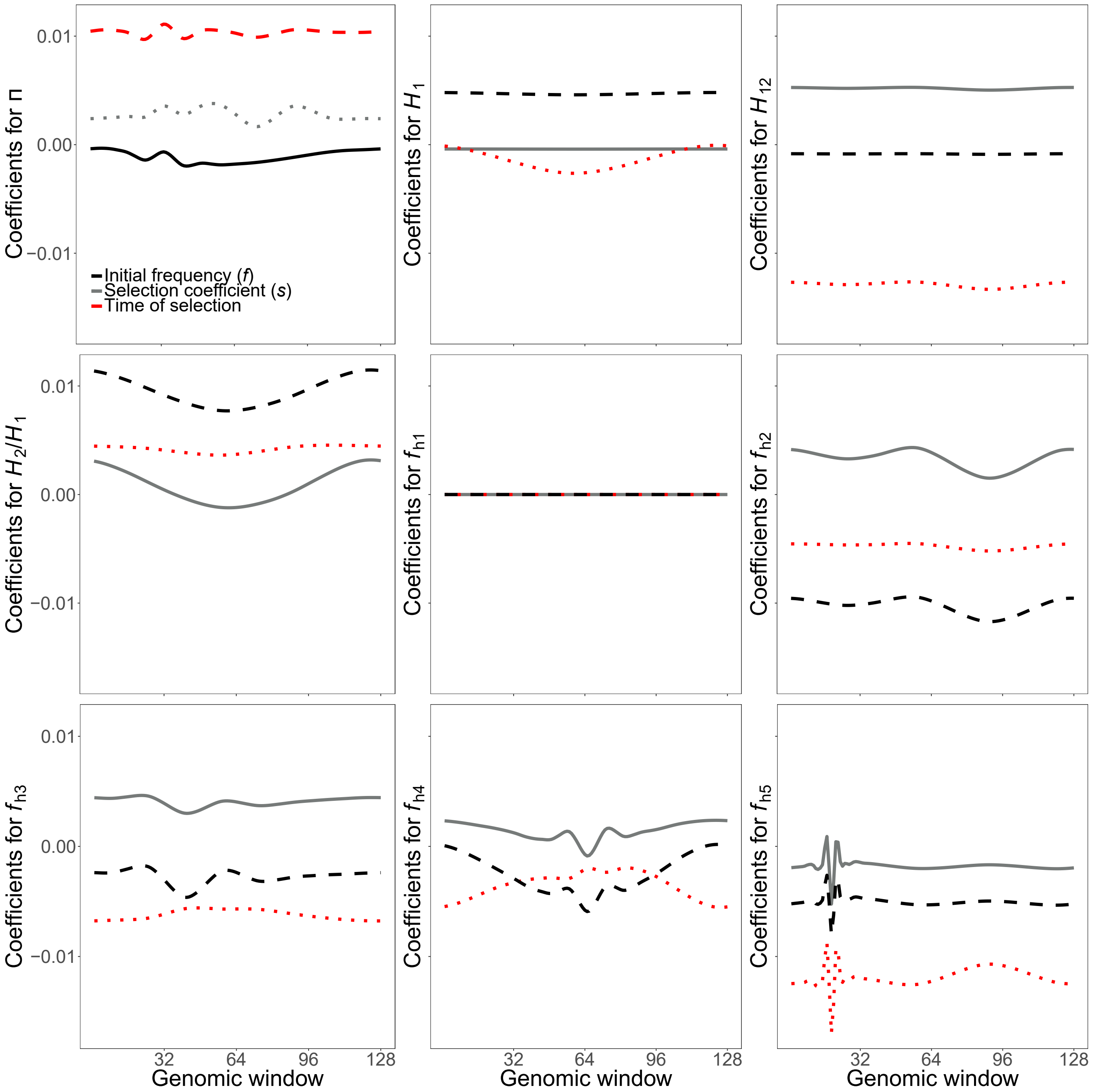


Figure S17: Reconstructed wavelets from regression coefficients ($\beta$s) in predicting time at which mutation became beneficial, frequency reached by mutation before becoming beneficial, and selection strength for summary statistics $\widehat{\pi}$, $H_1$, $H_{12}$, $H_2/H_1$, and frequencies of first to fifth most common haplotypes for *SURFDAWave* when $\gamma = 0.7$. *SURFDAWave* was trained on simulations of scenarios simulated under demographic specifications for sub-Sarahan African YRI demographic history. Note that the wavelet reconstructions for all summary statistics are plotted on the same scale, thereby making the distributions of some summaries difficult to decipher as their magnitudes are relatively small. *SURFDAWave* results shown are using Daubechies' least-asymmetric wavelets wavelets to estimate spatial distributions of summary statistics. Level 0 and $\gamma = 0.7$ chosen through cross-validation.

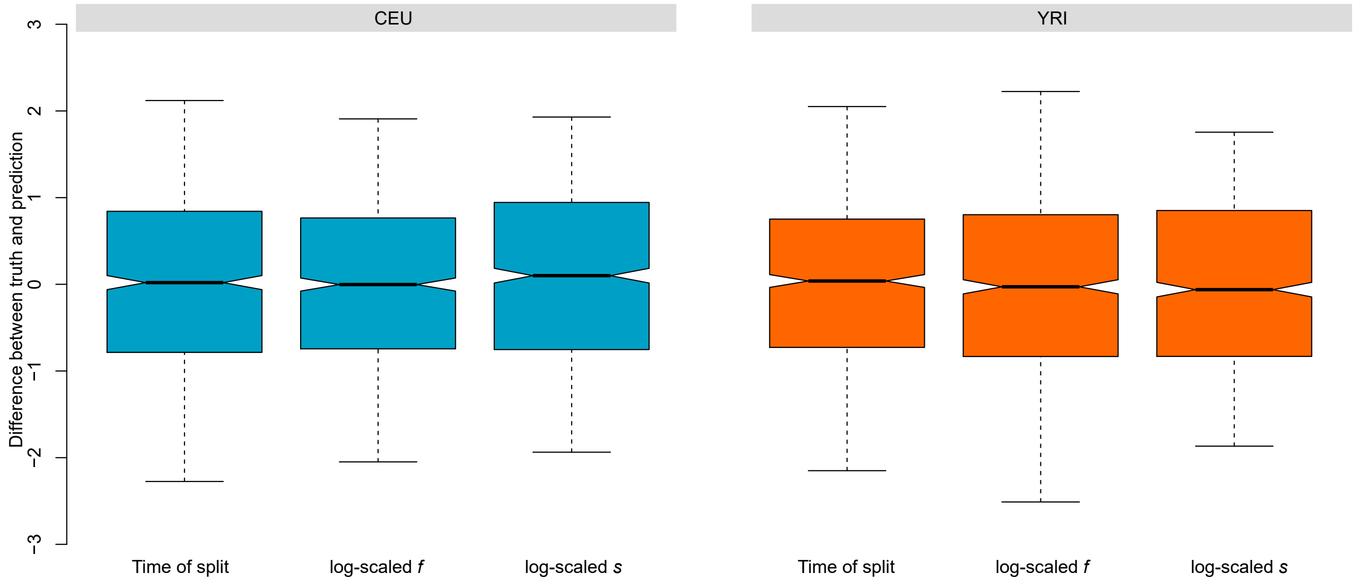


Figure S18: Difference between standardized predicted and actual selection parameters with *SURFDAWave* for the CEU and YRI demographic models. (Left box plot) Difference in prediction and truth of log scaled time at which donor and recipient populations split. (Middle box plot) Difference in prediction and truth of log scaled frequency reached by mutation prior to it becoming beneficial ($f$). (Right box plot) Difference in prediction and truth of log scaled selection coefficient ($s$).

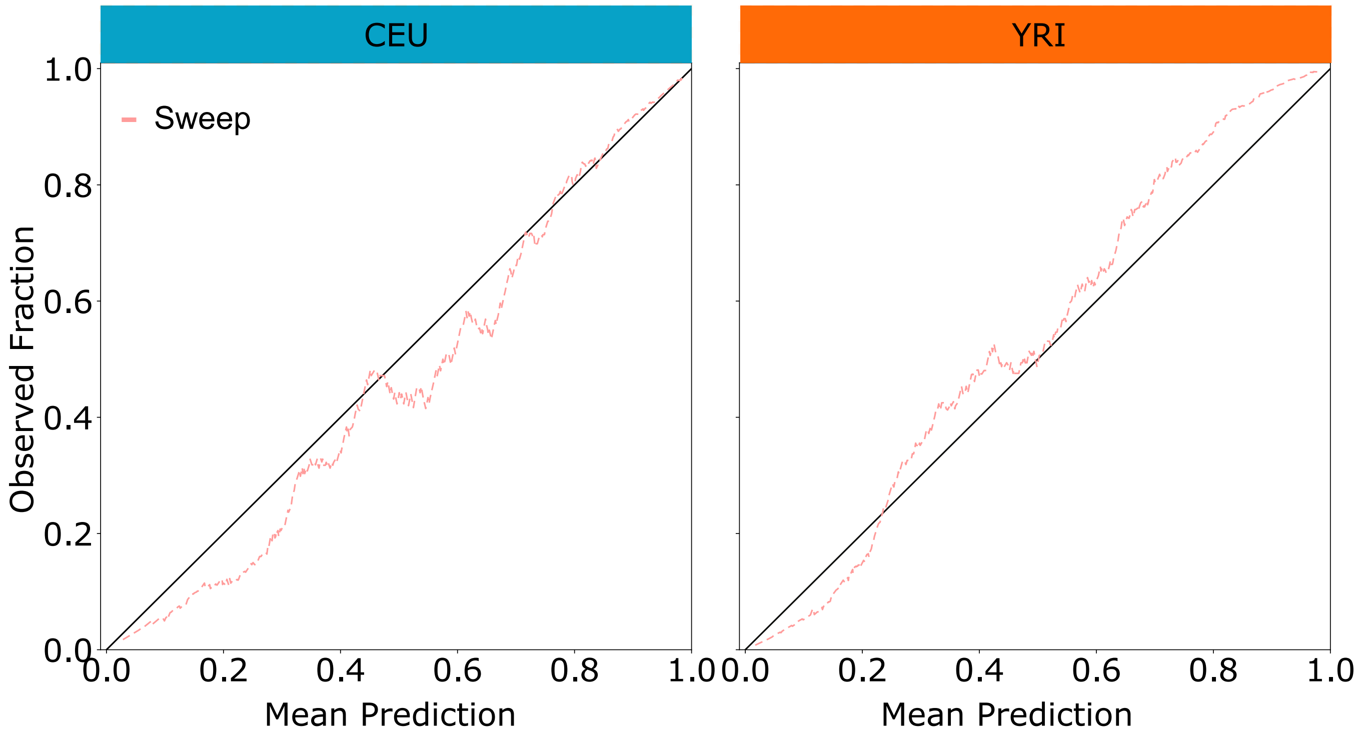


Figure S19: Reliability diagrams showing how close our predicted probabilities are to actual probabilities. For each classifier we predict the probability of sweep for test 1000 simulations. We divide the predicted probabilities into 950 overlapping windows each of length 0.5, with the first window beginning ranging from 0 to 0.05 and the second from 0.001 to 0.051 and so on with the last window ranging from 0.95 to 1.0. Using these ranges as thresholds, we calculate the mean probability of all predicted probabilities within this range (Mean Prediction) along with the fraction of these cases that are classified as sweep (Observed Fraction).

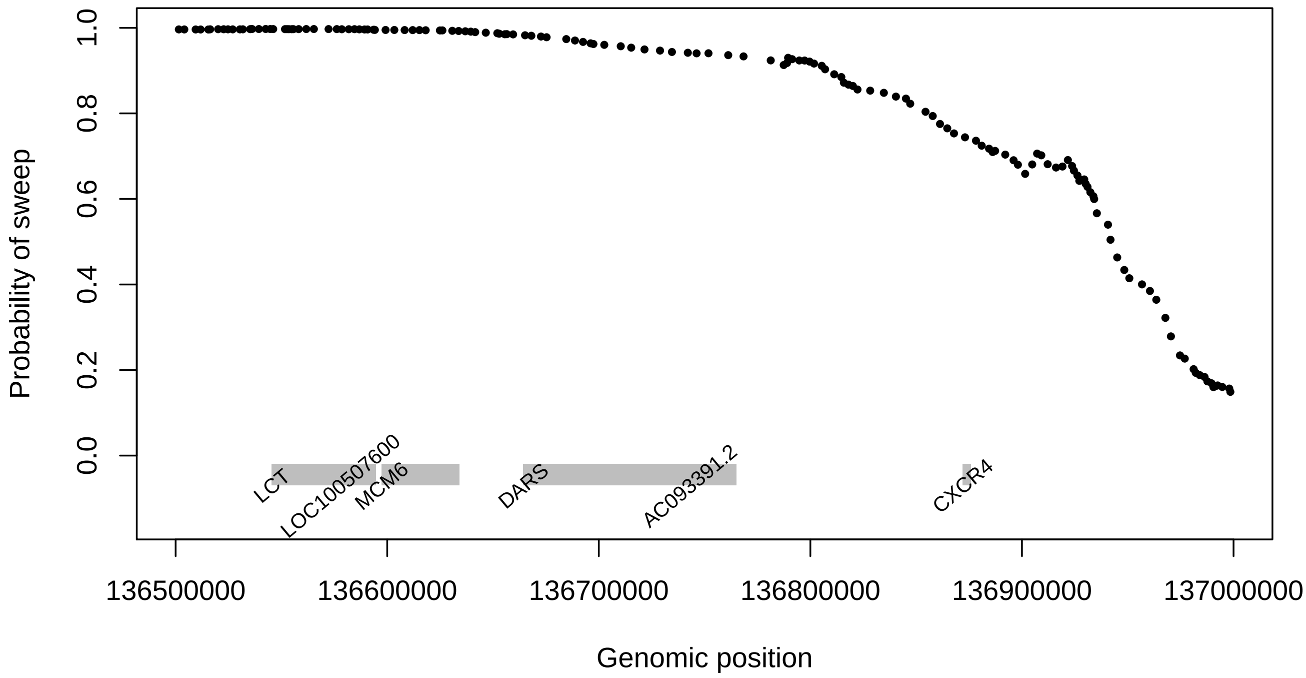


Figure S20: Probability of sweep in the CEU dataset across the genomic region of chromosome 2 containing the *LCT*, *MCM6*, and *DARS* genes. *SURFDAWave* is trained to differentiate between selective sweeps and neutrality with simulations conducted under demographic specifications of the CEU demographic history. The black dots show the predicted probability of sweep and the gray bars show the positions of the labeled genes.

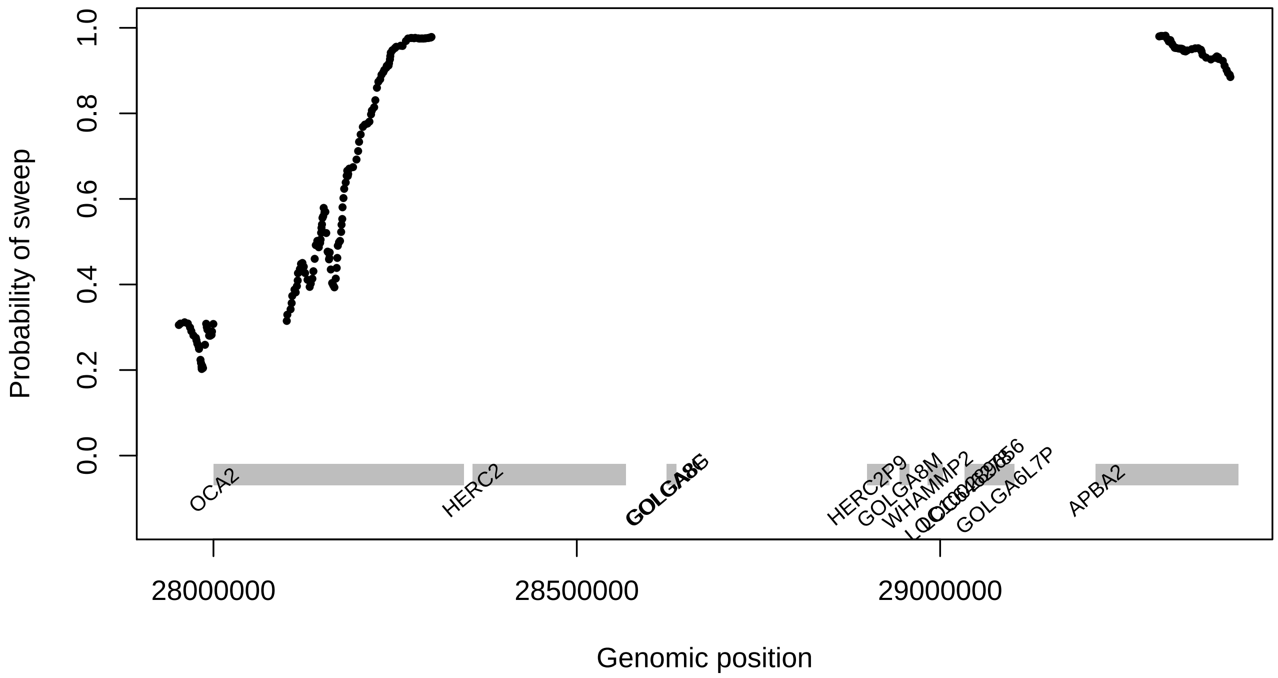


Figure S21: Probability of sweep in the CEU dataset across the genomic region of chromosome 15 containing the *OCA2* and *HERC2* genes. *SURFDAWave* is trained to differentiate between selective sweeps and neutrality with simulations conducted under demographic specifications of the CEU demographic history. The black dots show the predicted probability of sweep and the gray bars show the positions of the labeled genes.

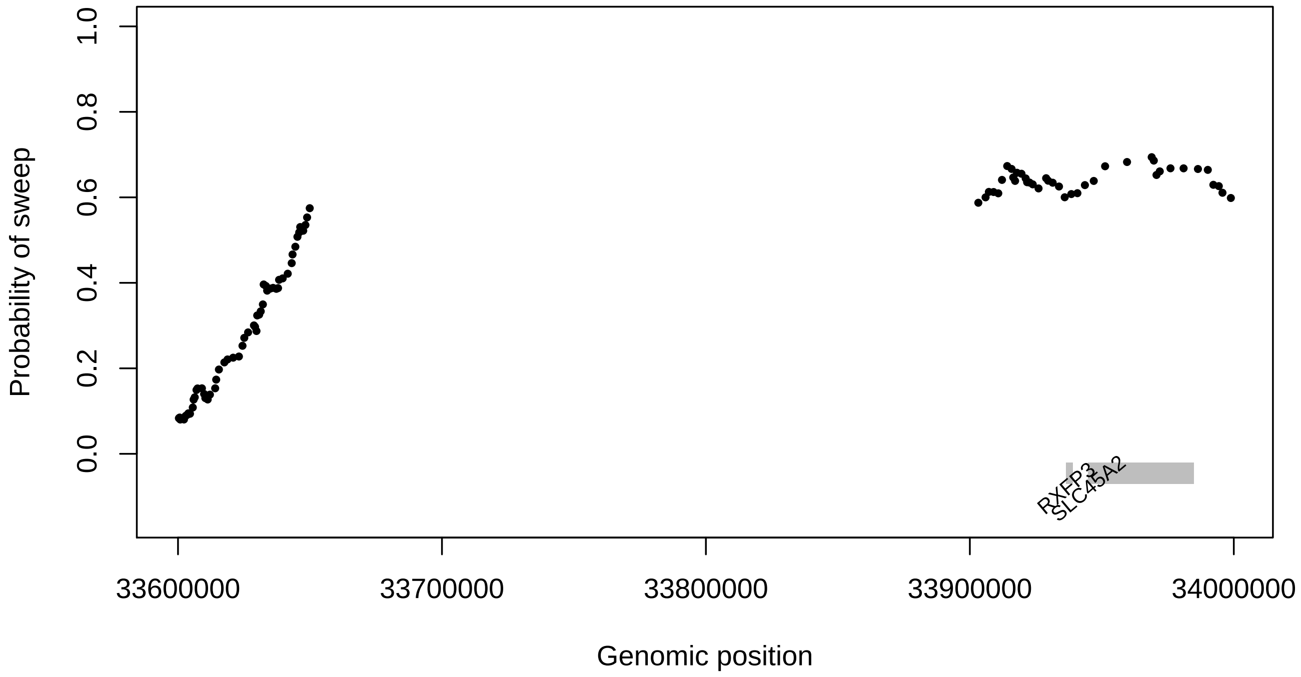


Figure S22: Probability of sweep in the CEU dataset across the genomic region of chromosome 5 containing the *SLC45A2* gene. *SURFDAWave* is trained to differentiate between selective sweeps and neutrality with simulations conducted under demographic specifications of the CEU demographic history. The black dots show the predicted probability of sweep and the gray bars show the positions of the labeled genes.

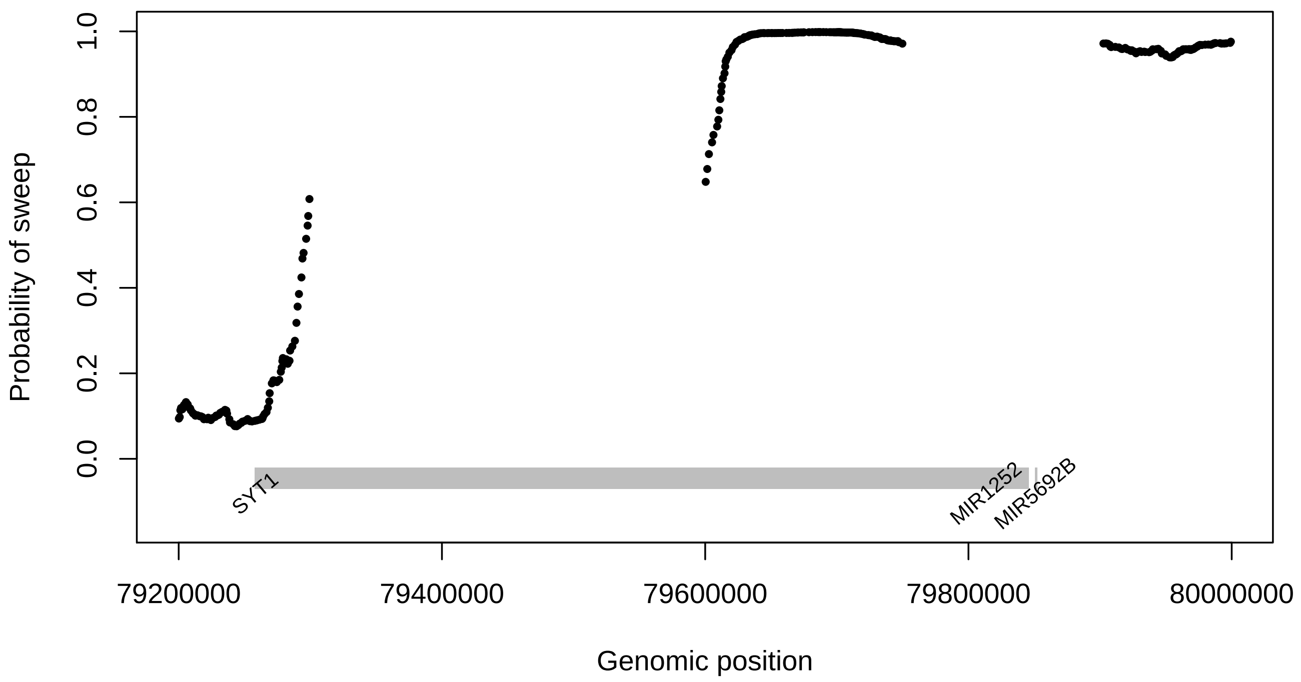


Figure S23: Probability of sweep in the YRI dataset across the genomic region of chromosome 12 containing the *SYT1* gene. *SURFDAWave* is trained to differentiate between selective sweeps and neutrality with simulations conducted under demographic specifications of the YRI demographic history. The black dots show the predicted probability of sweep and the gray bars show the positions of the labeled genes.

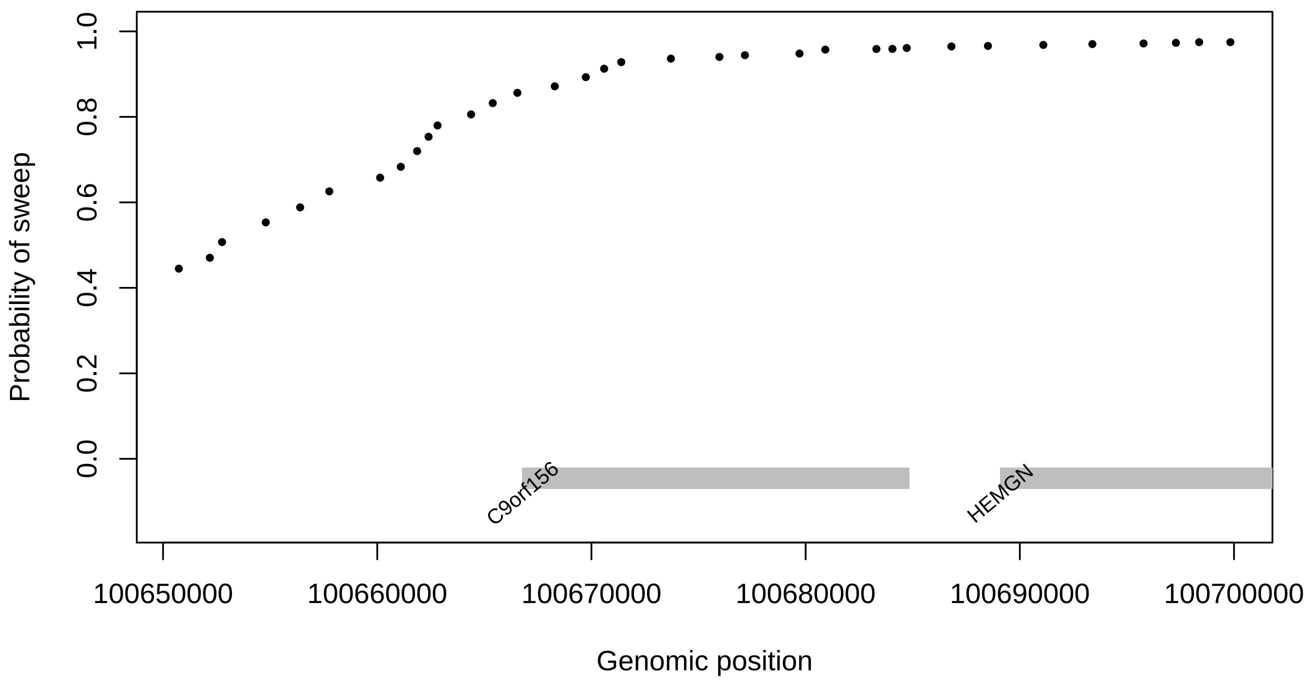


Figure S24: Probability of sweep in the YRI dataset across the genomic region of chromosome 9 containing the *HEMGN* gene. *SURFDAWave* is trained to differentiate between selective sweeps and neutrality with simulations conducted under demographic specifications of the YRI demographic history. The black dots show the predicted probability of sweep and the gray bars show the positions of the labeled genes.

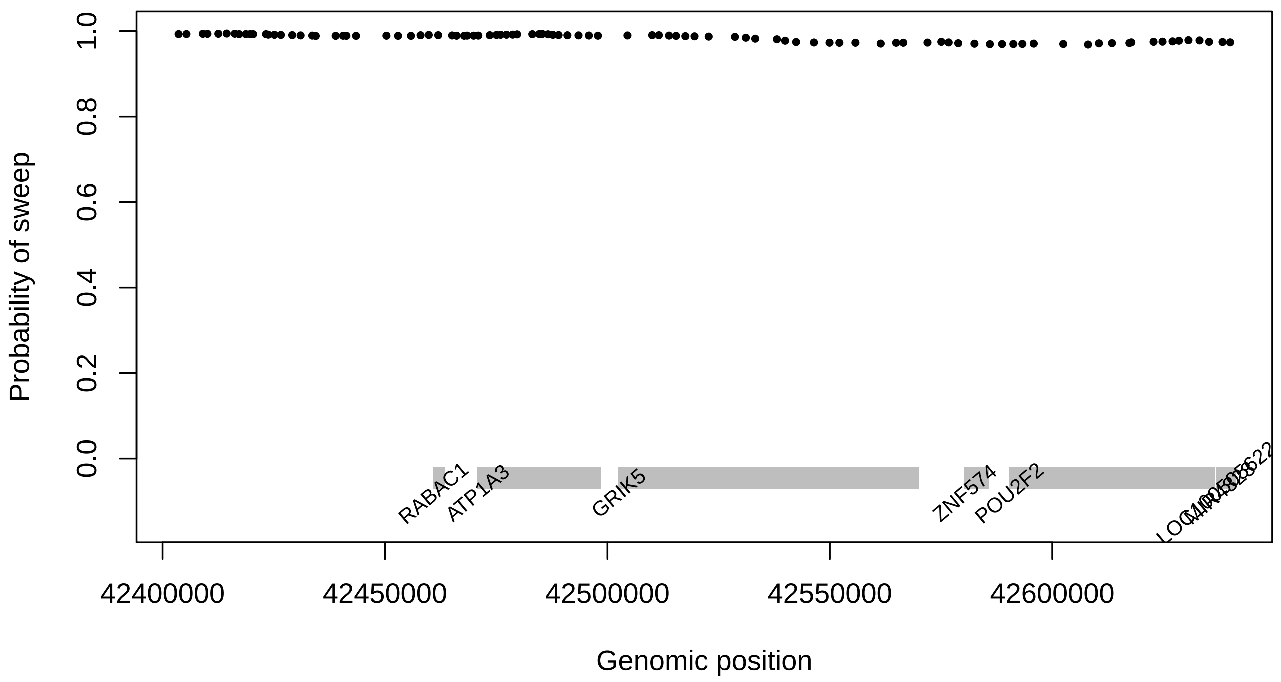


Figure S25: Probability of sweep in the YRI dataset across the genomic region of chromosome 19 containing the *GRIK5* gene. *SURFDAWave* is trained to differentiate between selective sweeps and neutrality with simulations conducted under demographic specifications of the YRI demographic history. The black dots show the predicted probability of sweep and the gray bars show the positions of the labeled genes.

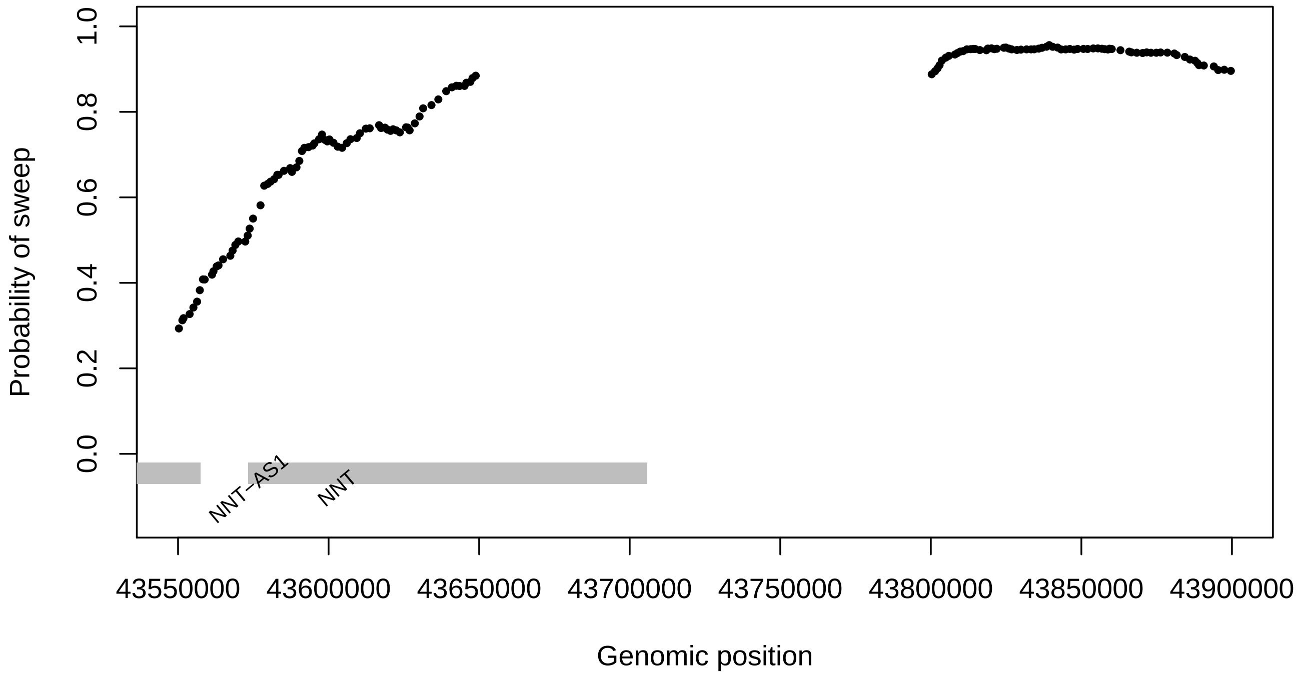


Figure S26: Probability of sweep in the YRI dataset across the genomic region of chromosome 5 containing the *NNT* gene. *SURFDAWave* is trained to differentiate between selective sweeps and neutrality with simulations conducted under demographic specifications of the YRI demographic history. The black dots show the predicted probability of sweep and the gray bars show the positions of the labeled genes.

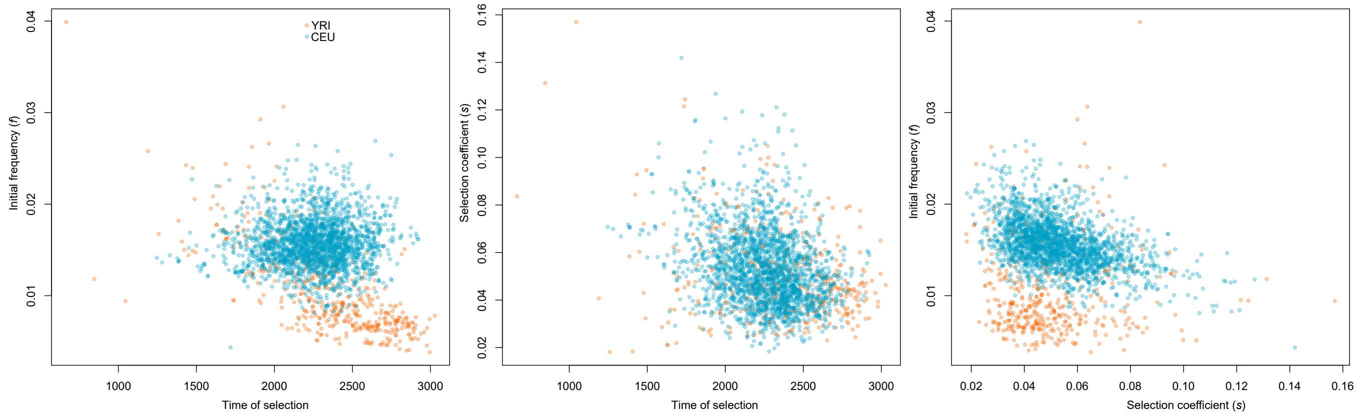


Figure S27: Predicted selection parameters for all genes in YRI and CEU with probability of being classified as sweep greater than 0.7 (Left) Scatter plot of predicted initial frequency mutation reached before becoming beneficial (Initial frequency) versus generations before present at which selection began (Time of selection). (Middle) Scatter plot of Selection coefficient versus Time of selection. (Right) Scatter plot of Initial frequency versus selection coefficient.

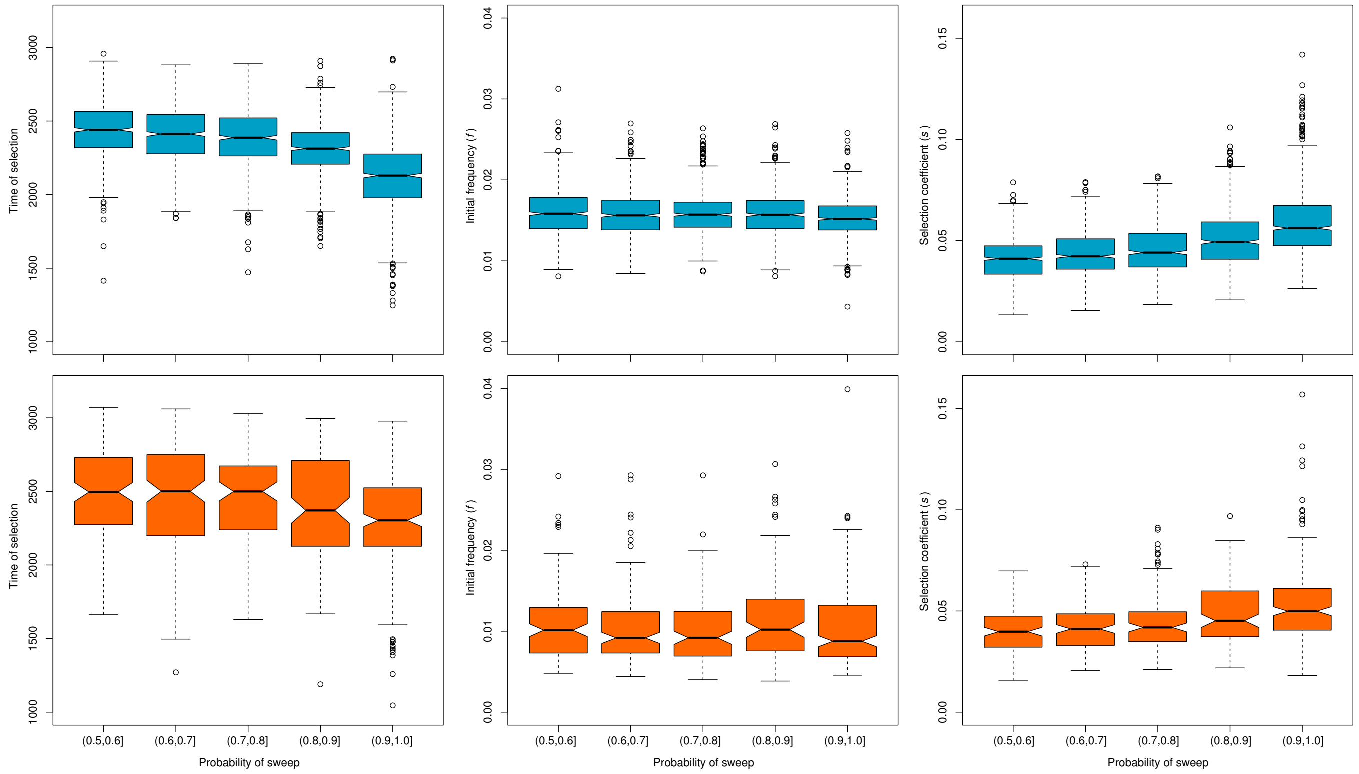


Figure S28: Predicted selection parameters for all genes in YRI (orange) and CEU (blue) with probability of being classified as sweep greater than 0.5 divided into bins of probability of sweep (Left) Predicted number of generations before present at which selection began (Time of selection) as a function of the probability of sweep. (Middle) Frequency reached by mutation before becoming beneficial ($f$) as a function of probability of sweep. (Right) Selection coefficient ($s$) as a function of probability of sweep.

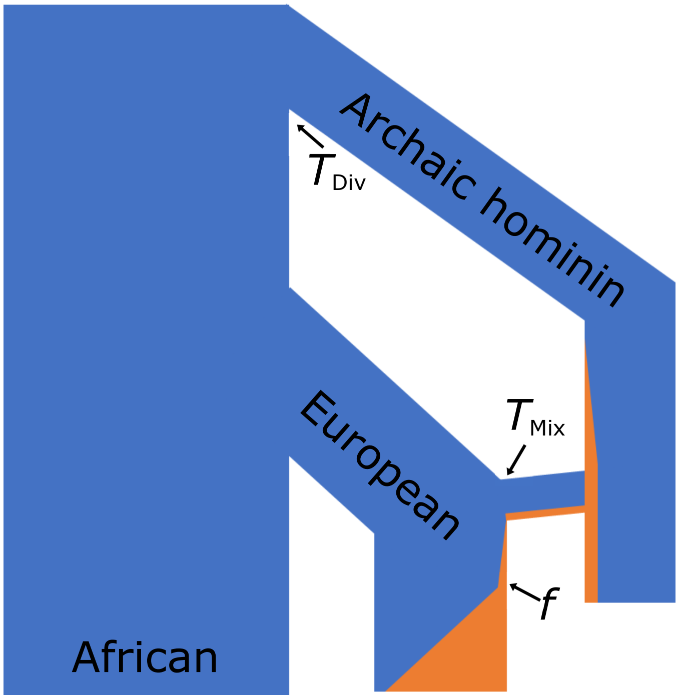


Figure S29: Demographic model depicting introgression into Europeans from an archaic homonin along with evolutionary parameters predicted by *SURFDAWave*. The width of the orange region represents the frequency of a mutation of interest, which originated in archaic homonin population, and became beneficial in Europeans. The evolutionary parameters we are trying to predict include the time at which archaic and modern human population diverged ($T_{\text{Div}}$), the time at the beneficial mutation entered the European population ($T_{\text{Mix}}$), and the initial frequency of the mutation reached before becoming beneficial in the modern humans ($f$).

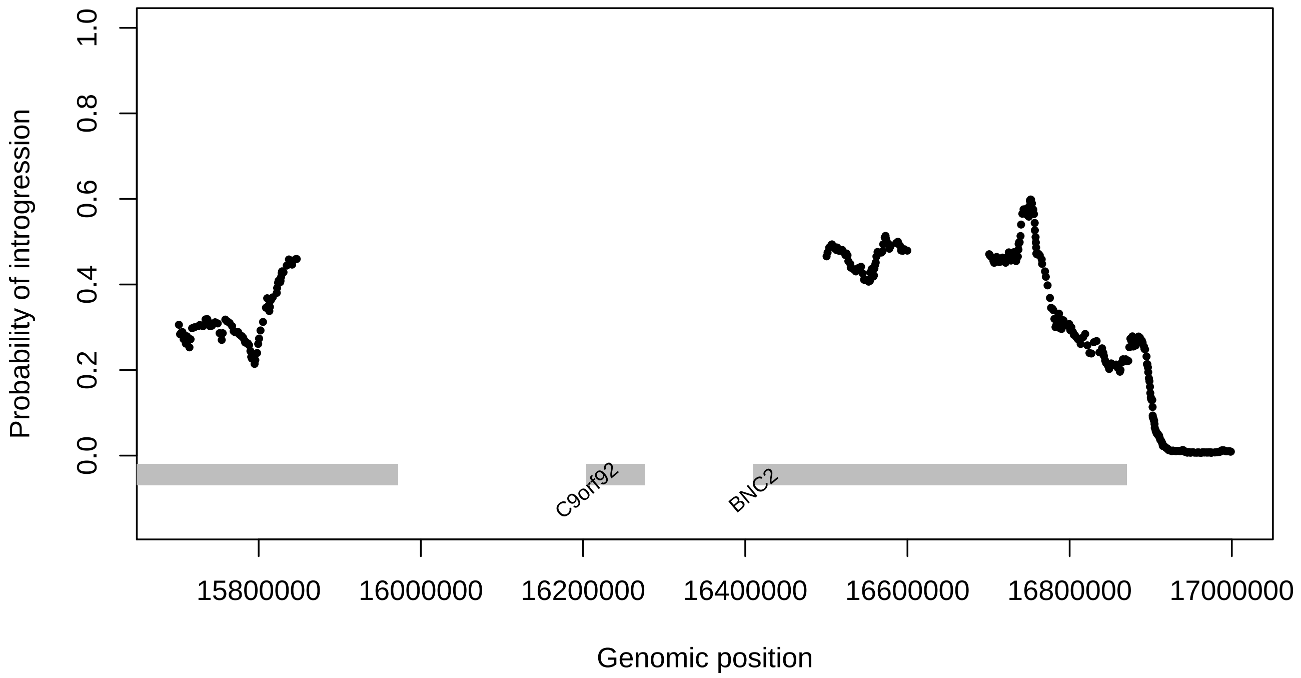


Figure S30: Probability of adaptive introgression in the CEU dataset across the genomic region of chromosome 9 containing the *BNC2* gene. *SURFDAWave* is trained to differentiate between selective sweeps, adaptive introgression, and neutrality with simulations conducted under demographic specifications of the CEU demographic history. The black dots show the predicted probability of sweep and the gray bars show the positions of the labeled genes.

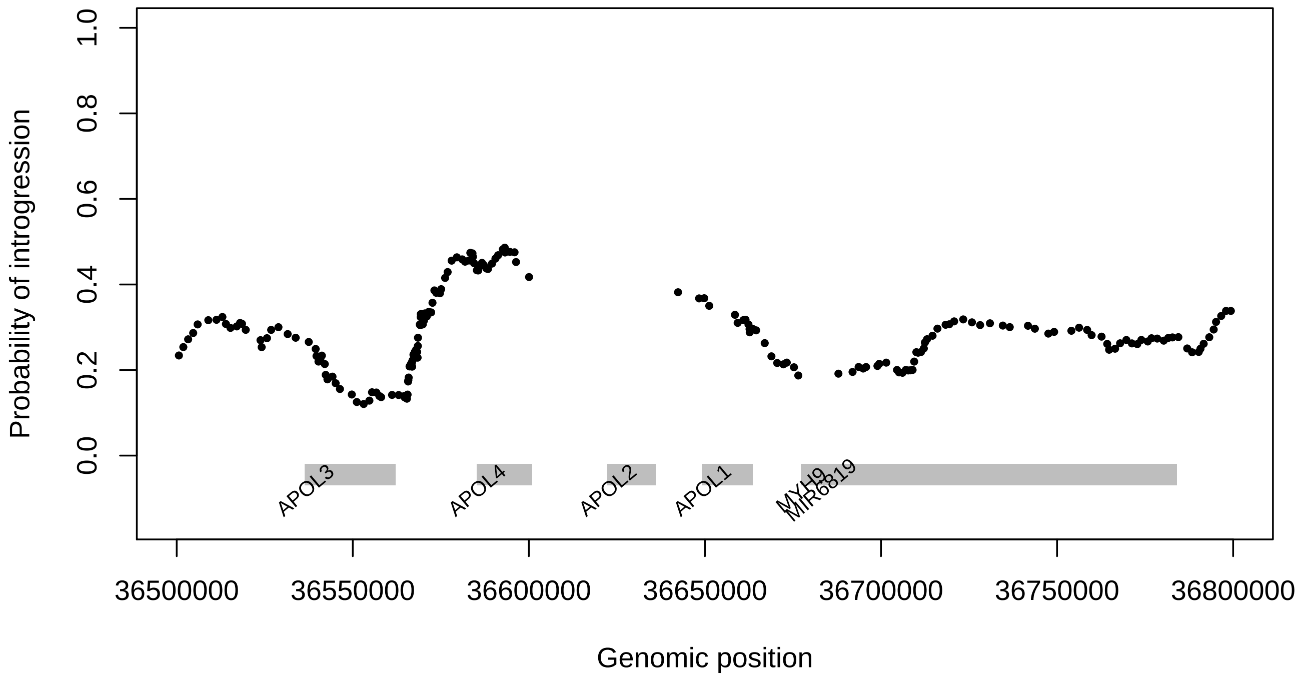


Figure S31: Probability of adaptive introgression in the CEU dataset across the genomic region of chromosome 22 containing the *APOL4* gene. *SURFDAWave* is trained to differentiate between selective sweeps, adaptive introgression, and neutrality with simulations conducted under demographic specifications of the CEU demographic history. The black dots show the predicted probability of sweep and the gray bars show the positions of the labeled genes.

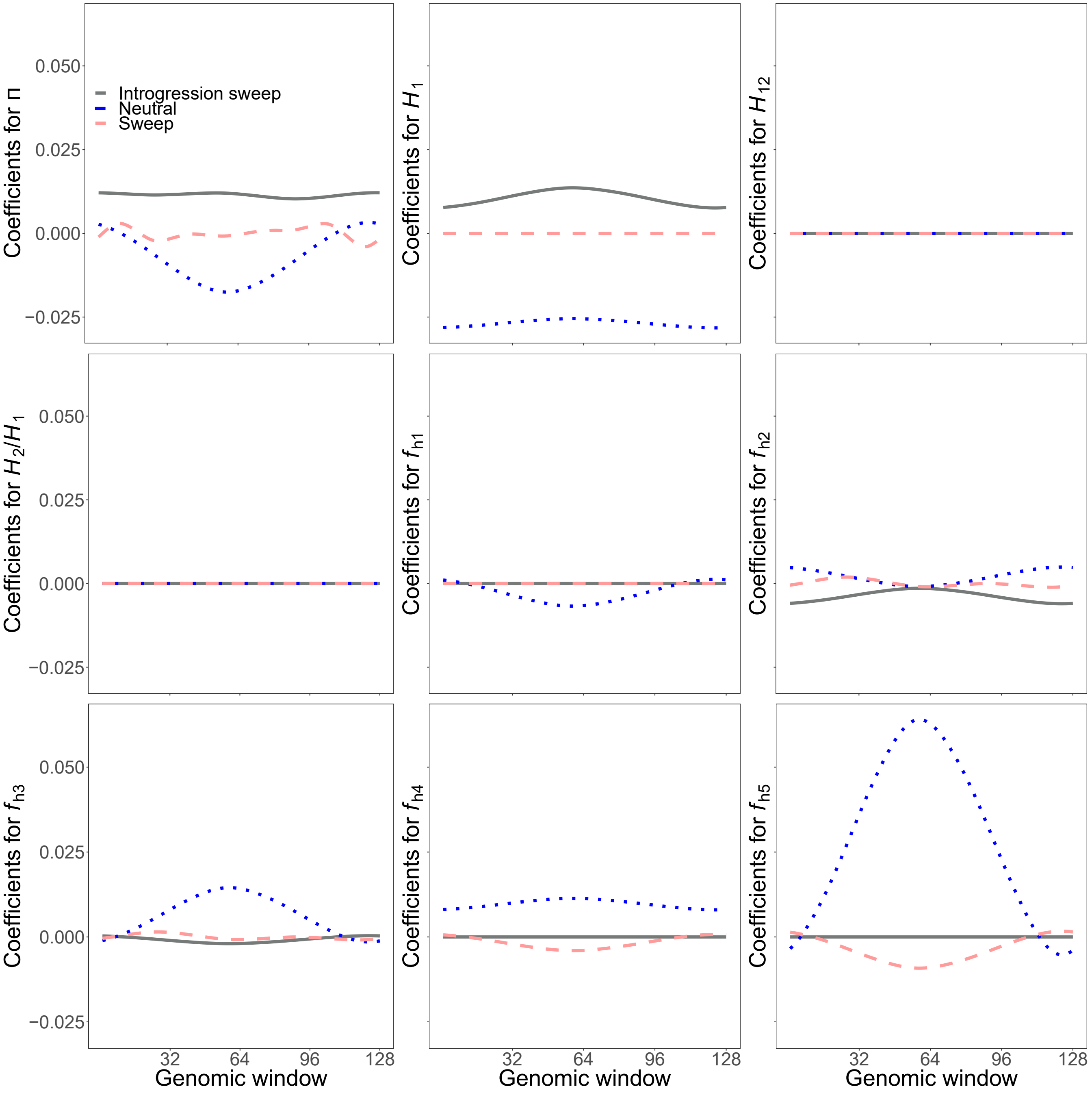


Figure S32: Reconstructed wavelets from regression coefficients ($\beta$s) when differentiating among adaptive introgression, sweeps, and neutrality scenarios for summary statistics $\widehat{\pi}$, $H_1$, $H_{12}$, $H_2/H_1$, and frequencies of first to fifth most common haplotypes for *SURFDAWave* when $\gamma = 1$. *SURFDAWave* was trained on simulations of scenarios simulated under demographic specifications for sub-Saharan African YRI demographic history. Note that the wavelet reconstructions for all summary statistics are plotted on the same scale, thereby making the distributions of some summaries difficult to decipher as their magnitudes are relatively small. *SURFDAWave* results shown are using Daubechies' least-asymmetric wavelets wavelets to estimate spatial distributions of summary statistics. Level 1 chosen through cross-validation.

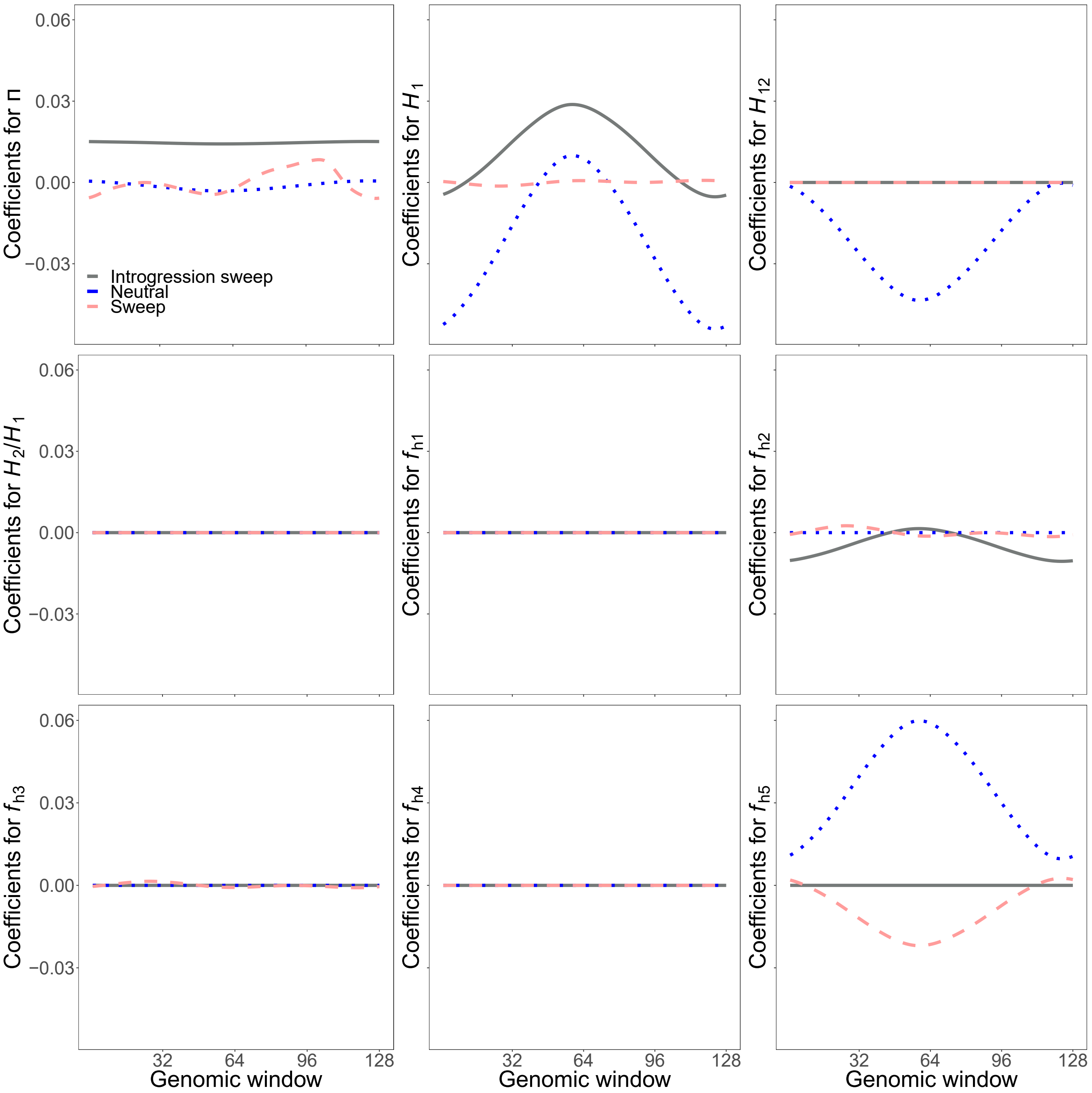


Figure S33: Reconstructed wavelets from regression coefficients ($\beta$s) when differentiating among adaptive introgression, sweeps, and neutrality scenarios for summary statistics $\widehat{\pi}$, $H_1$, $H_{12}$, $H_2/H_1$, and frequencies of first to fifth most common haplotypes for *SURFDAWave* when $\gamma = 1$, when trained with additional statistics mean, variance, skewness, and kurtosis of pairwise $r^2$ (Figure S34). *SURFDAWave* was trained on simulations of scenarios simulated under demographic specifications for sub-Saharan African YRI demographic history. Note that the wavelet reconstructions for all summary statistics are plotted on the same scale, thereby making the distributions of some summaries difficult to decipher as their magnitudes are relatively small. *SURFDAWave* results shown are using Daubechies' least-asymmetric wavelets wavelets to estimate spatial distributions of summary statistics. Level 1 chosen through cross-validation.

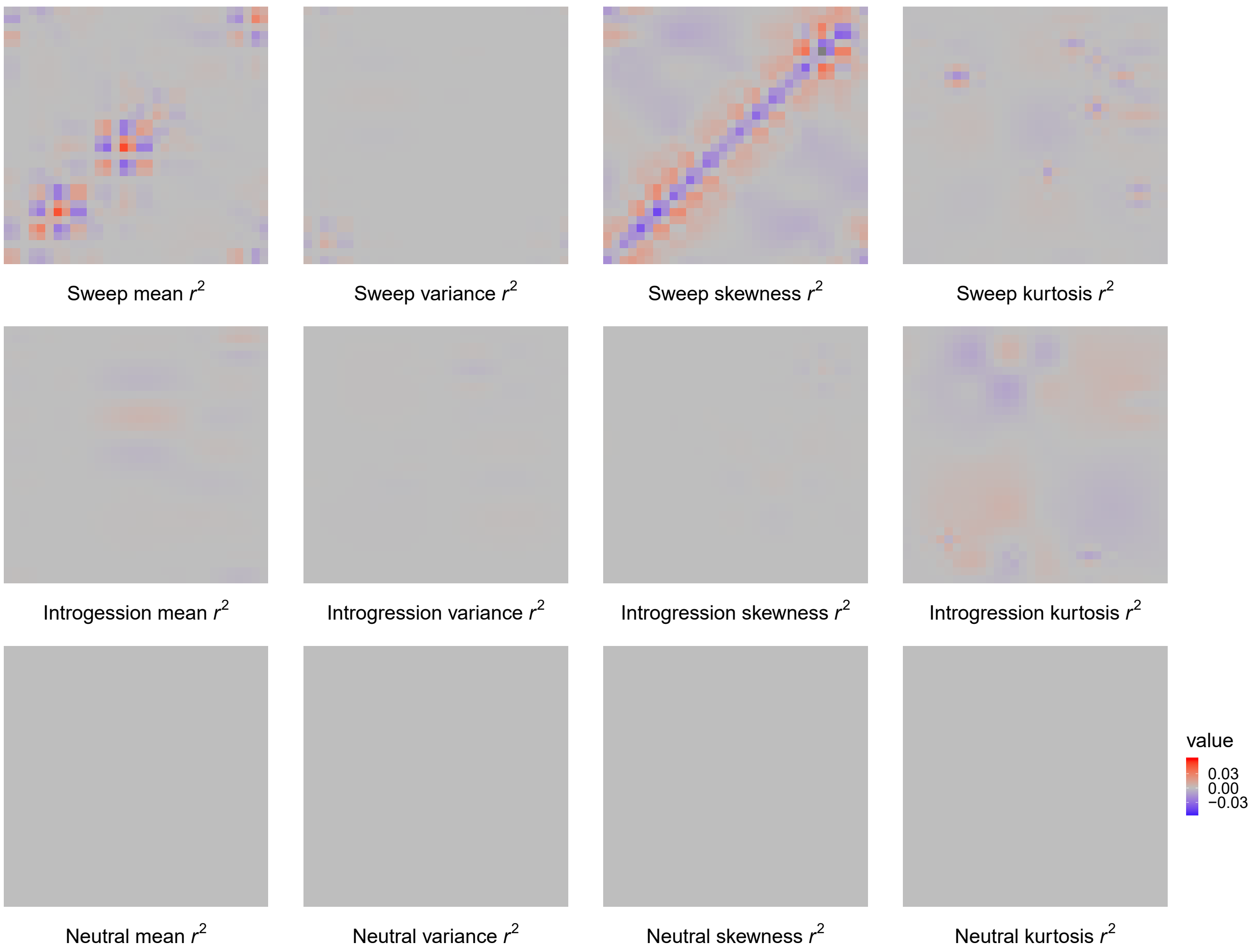


Figure S34: Reconstructed wavelets from regression coefficients ($\beta$s) when differentiating among adaptive introgression, sweeps, and neutrality scenarios for summary statistics mean, variance, skewness, and kurtosis of pairwise $r^2$ for *SURFDAWave* when $\gamma = 1$, when trained with statistics $\widehat{\pi}$, $H_1$, $H_{12}$, $H_2/H_1$, and frequencies of first to fifth most common haplotypes (Figure S33). *SURFDAWave* was trained on simulations of scenarios simulated under demographic specifications for sub-Saharan African YRI demographic history. Note that the wavelet reconstructions for all summary statistics are plotted on the same scale, thereby making the distributions of some summaries difficult to decipher as their magnitudes are relatively small. *SURFDAWave* results shown are using Daubechies' least-asymmetric wavelets wavelets to estimate spatial distributions of summary statistics. Level 1 chosen through cross-validation.

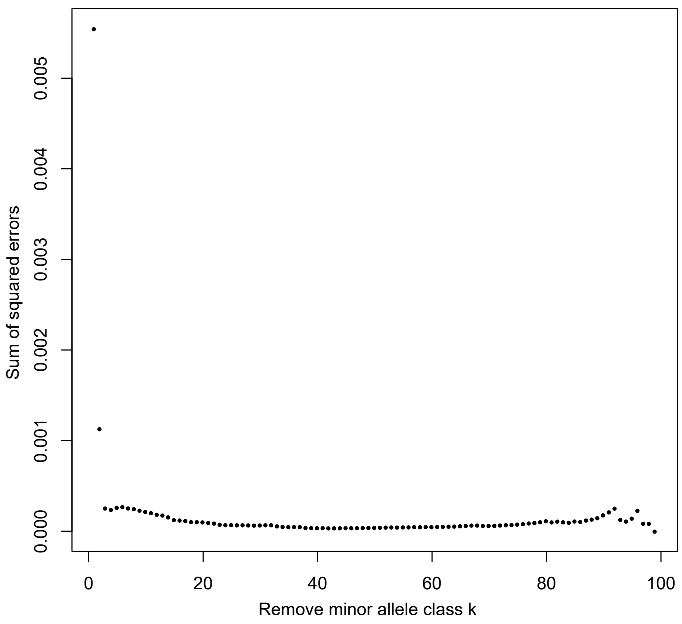


Figure S35: Sum of squared differences between the empirical CEU and the simulated neutral Terhorst normalized minor allele frequency spectra conditional on removing all minor allele classes with k or fewer minor alleles.

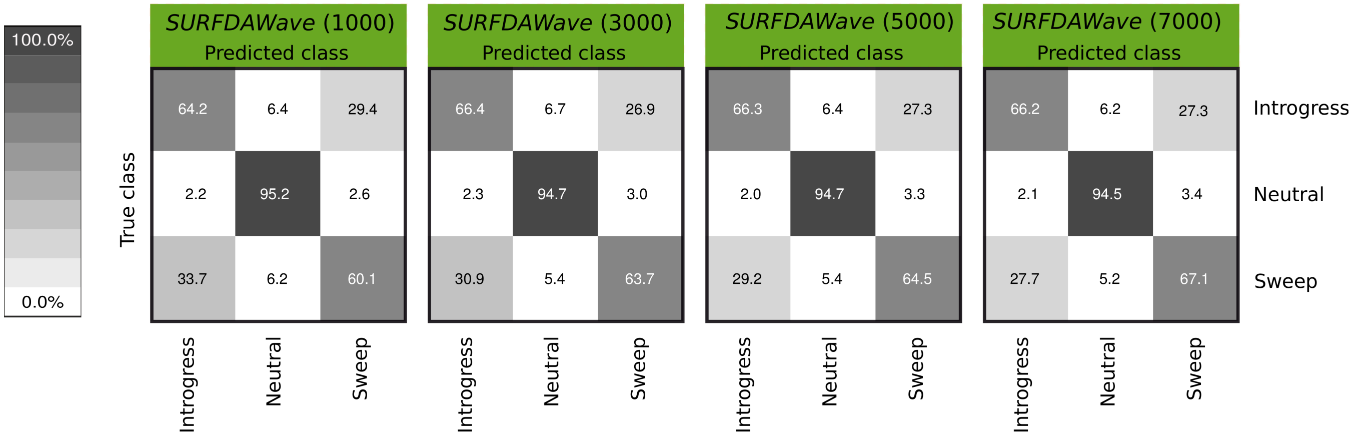


Figure S36: Confusion matrices comparing classification rates of *SURFDAWave* differentiating among non-introgressed sweeps, introgressed sweeps, and neutrality when simulated under a constant-size demographic model with non-adaptive introgression with 1000, 3000, 5000, or 7000 training samples per class.

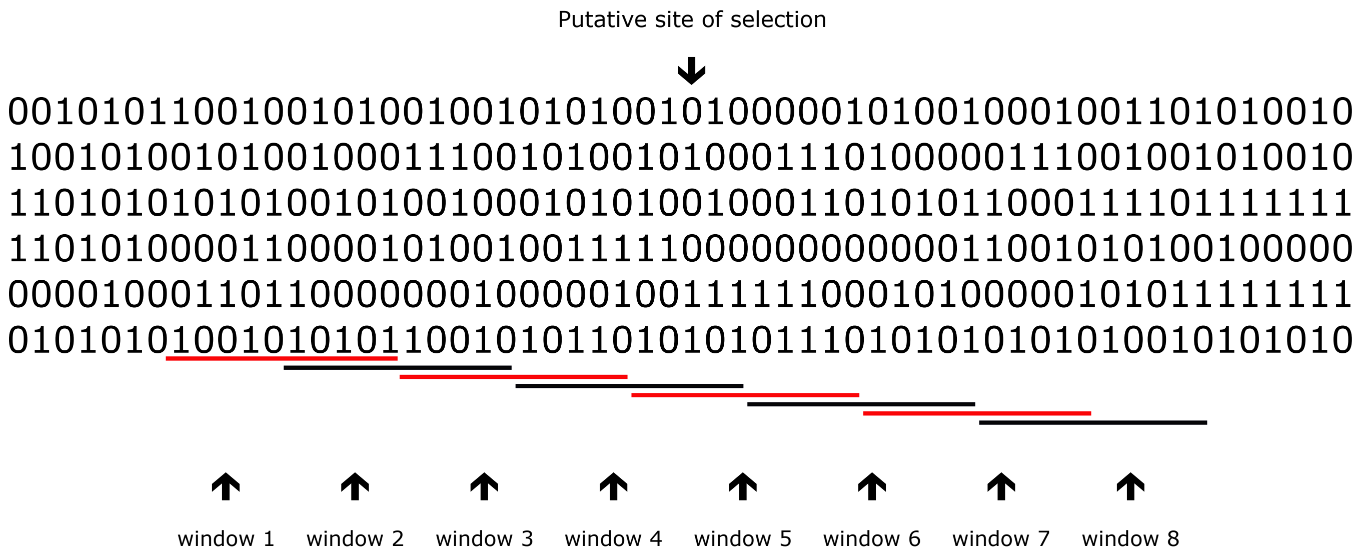


Figure S37: Schematic illustrating windows for which summary statistics are calculated in our implementation of *SURFDAWave*. Each bold black line underlines one of the eight 10-SNP long windows. Here we show a sample of six haplotypes (rows) across a string of SNPs (columns) for which we calculate summary statistics in $p = 8$ windows. Summary statistics are calculated for each 10-SNP window, with windows overlapping with each neighbor for five SNPs. The central SNP is taken to be the putative selected site and is located in the overlap of windows four and five. Here we have underlined the alternating windows used to calculate the two-dimensional statistics in red.

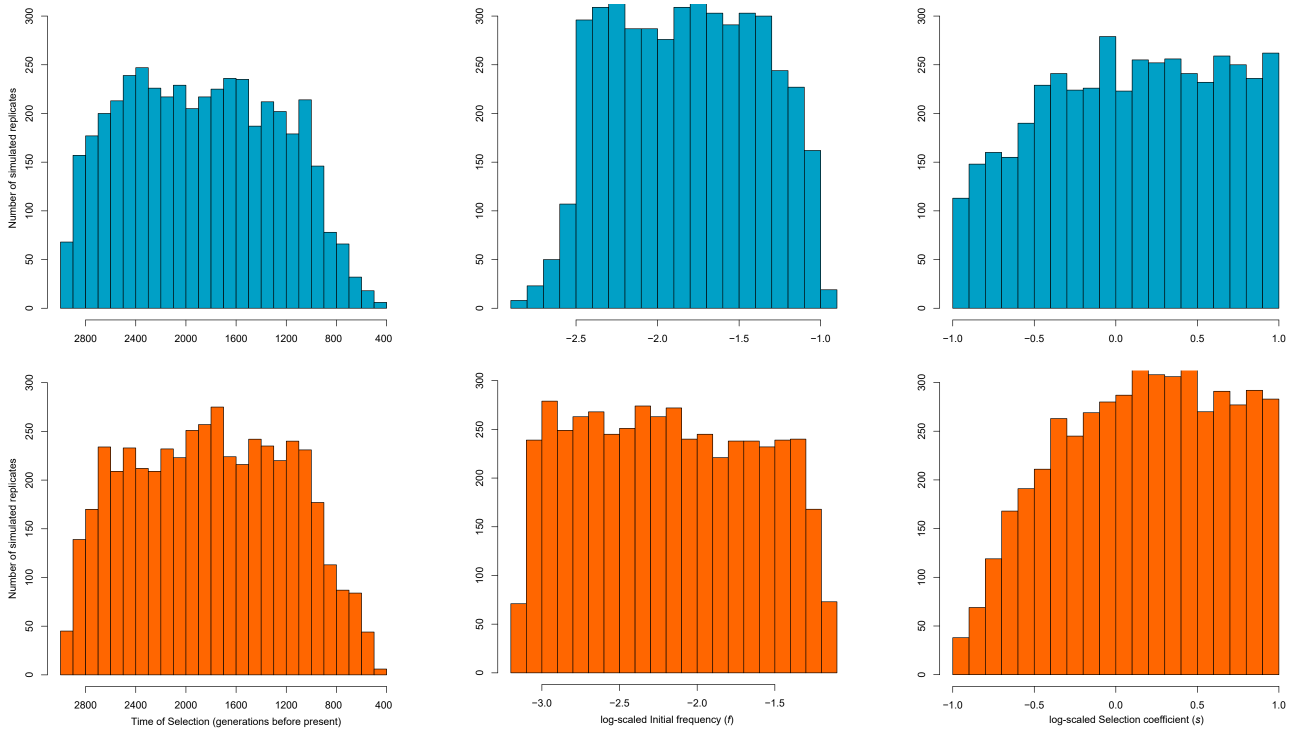


Figure S38: Distribution of selection parameters for simulations of sweeps conducted with demographic history parameters of CEU (top row) and YRI (bottom row). (Left column) Distribution of time at which tracked mutation becomes beneficial (reaches initial frequency) in simulations of selective sweeps. (Middle column) Distribution of log-scaled initial frequency (input parameter) reached by mutation before becoming beneficial in simulations of selective sweeps. (Right column) Distribution of log-scaled selection coefficient in simulations of selective sweep.
